# Revisiting the Natural History of Pulmonary Tuberculosis: a Bayesian Estimation of Natural Recovery and Mortality rates

**DOI:** 10.1101/729426

**Authors:** Romain Ragonnet, Jennifer A. Flegg, Samuel L. Brilleman, Edine Tiemersma, Yayehirad A. Melsew, Emma S. McBryde, James M. Trauer

## Abstract

**Background:** Tuberculosis (TB) natural history remains poorly characterised and new investigations are impossible as it would be unethical to follow up TB patients without treating them. Estimates of TB burden and mortality rely heavily on TB self-recovery and mortality rates, as around 40% of individuals with TB are never detected, making their prognosis entirely dependent on the disease natural history.

**Methods:** We considered the reports identified in a previous systematic review of studies from the prechemotherapy era, and extracted detailed data on mortality over time. We used a continuous-time Markov model in a Bayesian framework to estimate the rates of TB-induced mortality and self-cure. A hierarchical model was employed to allow estimates to vary by cohort. Inference was performed separately for smear-positive TB (SP-TB) and smear-negative TB (SN-TB).

**Results:** We included 41 cohorts of SP-TB patients and 19 cohorts of pulmonary SN-TB patients in the analysis. No data were available on extrapulmonary TB. The posterior median estimates of the TB-specific mortality rates were 0.390 year^−1^ (0.329-0.452, 95% credible interval) and 0.025 year^−1^ (0.016-0.036) for SP-TB and SN-TB patients, respectively. The estimates for self-recovery rates were 0.233 year^−1^ (0.179-0.293) and 0.147 year^−1^ (0.087-0.248) for SP-TB and SN-TB patients, respectively. These rates correspond to average durations of untreated TB of 1.57 years (1.37-1.81) and 5.35 years (3.42-8.23) for SP-TB and SN-TB, respectively, when assuming a natural mortality rate of 0.014 year^−1^ (i.e. a 70-year life expectancy).

**Conclusions:** TB-specific mortality rates are around 15 times higher for SP-TB than for SN-TB patients. This difference was underestimated dramatically in previous TB modelling studies that parameterised models based on the ratio of 3.3 between the 10-year case fatality of SP-TB and SN-TB. Our findings raise important concerns about the accuracy of past and current estimates of TB mortality and predicted impact of control interventions on TB mortality.

## Introduction

The TB plague has threatened mankind since prehistory. Hippocrates described the disease (then referred to as “phthisis”) as “the worst of the diseases that occurred, alone responsible for the great mortality” around 2,400 years ago (1). Today, TB is still the world’s most lethal disease from a single infectious agent (2). One obstacle to efficient TB control is the lack of knowledge concerning some of the most fundamental aspects of TB epidemiology, such as the natural history of TB. This phenomenon is impossible to investigate using modern data, as antibiotics should be systematically provided to all individuals diagnosed with TB (3). And yet, the prognosis of untreated individuals remains of critical importance, as currently around 40% of diseased individuals are never identified and this proportion reaches even higher levels in settings where TB control functions poorly and where TB is the most likely to strike (4, 5).

Characterising TB natural history accurately is all the more important because it is a central component of the methodology currently used to produce disease burden estimates. In particular, TB incidence is often estimated by dividing the disease prevalence – generally obtained from field surveys – by the estimated average duration of a TB episode, the latter relying significantly on estimates of spontaneous recovery and TB mortality rates (6). The poor characterisation of these parameters is extremely concerning, as TB incidence is used as the primary burden indicator in global TB control and by the main global health donors when allocating funding between countries (7). Moreover, as TB control policies increasingly rely on predictions based on mathematical models, it is essential to ensure that the natural history of the disease is accurately captured by such systems (8).

A systematic review aimed at better defining the prognosis of untreated TB was published in 2011 (9). The review only included TB literature from the pre-chemotherapy era. This review represents a landmark for TB epidemiology, as it provided a comprehensive overview of all the available reports on untreated TB patients and presented quantitative estimates of TB case fatality.

Specifically, this study reported a 70% 10-year case fatality for smear-positive TB (SP-TB) patients and 20% for smear-negative TB (SN-TB) patients. Since its publication, this review has constituted the main source of parameterisation for the natural history of TB in mathematical models and is used by WHO to estimate TB incidence. However, because this study was not designed to provide modelling guidance specifically, the optimal approach to interpreting these results and incorporating them into mathematical systems remains unclear.

For example, the case fatality proportions cited above provide the final outcome but do not describe mortality over time or the spontaneous recovery rates, both of which are critical in disease modelling. Moreover, while these estimates were obtained from aggregation across several cohorts, the variability around the reported values due to the heterogeneity between cohorts has not been quantified and such information is crucial for modellers to produce informative estimates. Finally, it is to be noted that these rates include both the TB-induced mortality and natural mortality, as they are based on overall fatality rates estimated from survival proportions of TB patients over time. Since the contribution from natural mortality is not explicitly reported, these estimates cannot be updated to reflect changes in natural mortality rates over time (10).

In the current paper, we propose a re-investigation of the TB prognosis data from the pre-chemotherapy era by taking a mathematical modelling approach to estimate TB-specific mortality rates and spontaneous recovery rates.

## Methods

### Literature review and data extraction

We considered the manuscripts that were identified in the previous systematic review of studies from the prechemotherapy era (9), and extracted data on mortality over time. The reports present aggregated cumulative death or survival proportions over time, with the starting time usually being the time of notification or of admission to a hospital or sanatorium. The number of reported timepoints varied between one and 31 and the follow-up duration between one and 31 years. Most studies reported on several cohorts of TB patients, often disaggregated by year of diagnosis or gender. Overall, 64 cohorts were extracted from the 15 reports that were found to contain data on case fatality. These included 41 groups of SP-TB patients (often referred to as “open TB cases”) and 23 groups of SN-TB patients (“closed TB cases”). Only 19 SN-TB cohorts could be included in the quantitative analysis, as the number of patients was not reported in four cohorts. All cohorts reported on pulmonary TB and we employed the classification that was used in the previous review to distinguish SP-TB from SN-TB (9).

All studies originated from Western Europe, with four reports from England, three from Denmark, two from the Netherlands, and one each from: Poland, Norway, Switzerland, Sweden, Germany and Iceland. Six studies reported on sanatorium patients, six on officially notified individuals with TB and three on hospital or dispensary patients. Cohort sizes varied between eight and 2,382, with a median size of 379 patients. The detailed cohort profiles are provided in the Supplement. Figure 1 presents the raw mortality proportions over time extracted from the different reports.

**Figure 1.**
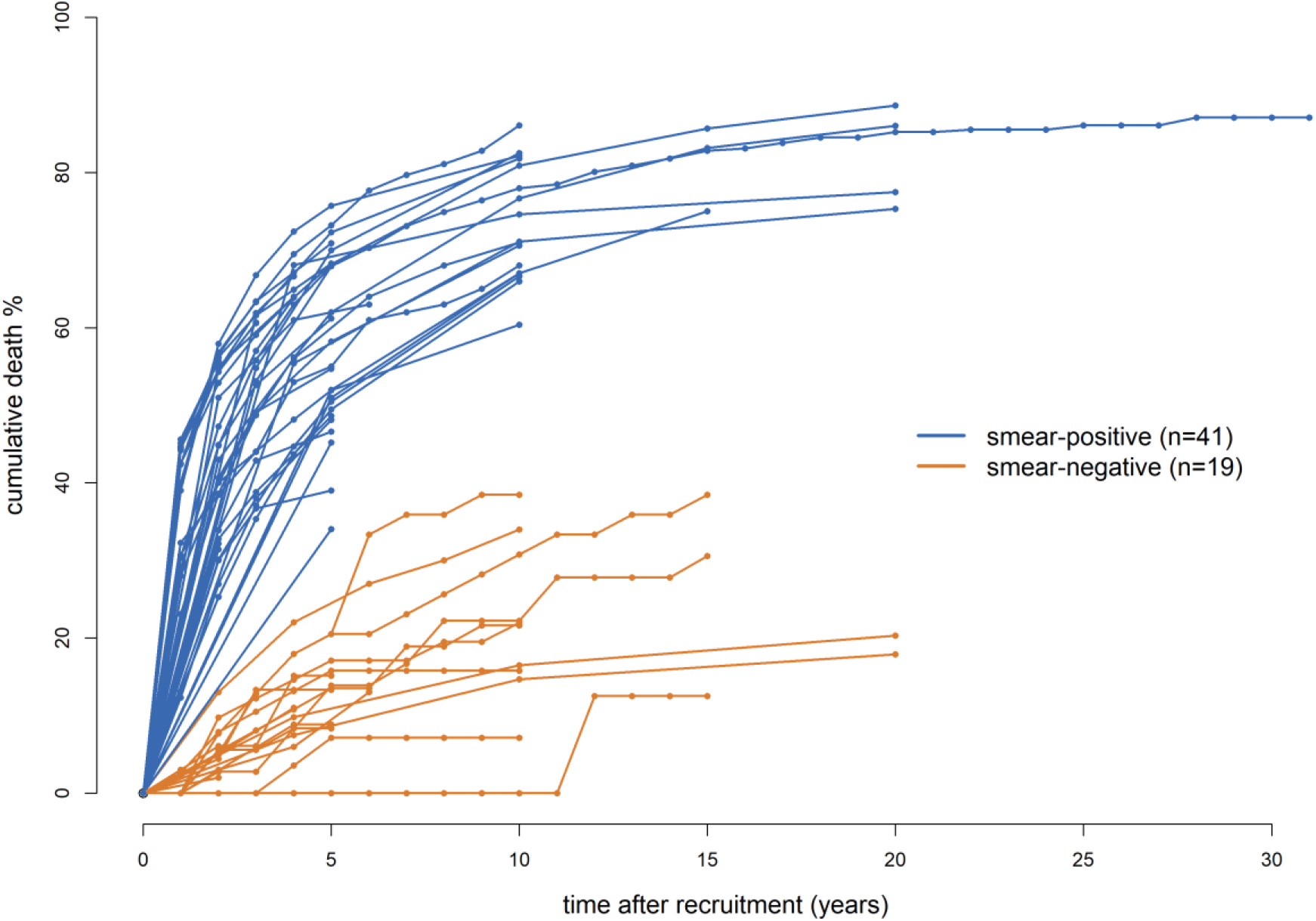
Observed cumulative percentage of death in the cohorts of TB patients identified. Each line connecting a series of dots represents a single patient cohort, with each dot representing an observation point for that cohort. Blue lines represent smear-positive patients (open TB) and orange lines represent smear-negative patients (closed TB).

### Model and parameters

To estimate the TB mortality and recovery rates, we used a continuous-time homogeneous Markov chain that mimics the progression between the main stages that comprise the natural history of TB: active TB, self-recovery and death. Using this approach, transitions between the different states are allowed at any time and governed by constant transition rates. Diseased individuals may spontaneously recover (at rate *γ*) or die (at rate *μ* + *μ_T_*) from TB, while recovered individuals die only from natural causes at rate *μ*. An illustration of this model is shown in Figure 2. The main objective of this study was to estimate the parameters *γ* and *μ_T_*.

**Figure 2.**
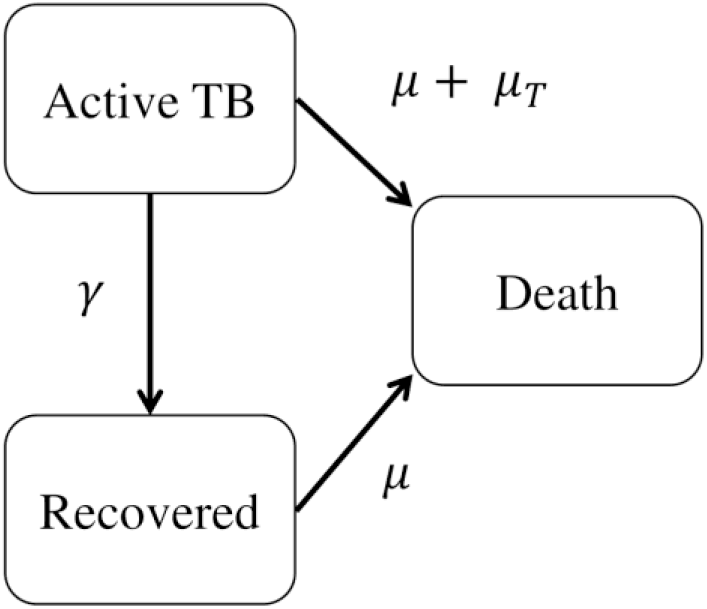
Representation of the pathways to death and the parameters characterising TB natural history. *γ* is the rate of spontaneous recovery, *μ* is the natural mortality rate and *μ_T_* is the additional mortality rate due to TB disease.

### Parameter estimation

We used a Bayesian approach to estimate the parameters characterising TB natural history. For each cohort *i*, the extracted data provide the numbers of deaths 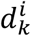 occurring during the time-interval 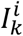 where *k* is in [and where *m^i^* is the number of time-intervals reported for cohort *i*. We also know the number of people 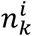 that are still alive (at-risk) at the beginning of each interval 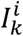. Therefore, the quantity 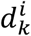 is the realisation of a binomial process 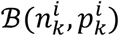, where 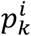 is the individual probability of death within time-interval 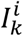, provided that the individual was alive at the beginning of 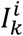. The probabilities 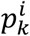 were computed analytically from the Markov process introduced in Figure 2 (see Supplement Section 5 for details). These probabilities therefore depend on the TB natural history parameters (*γ* and *μ_T_*), the natural mortality rate (*μ*), as well as the starting time and length of the intervals 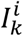.

We used a hierarchical approach to allow for *γ* and *μ_T_* to vary by cohort (11). That is, the associated parameters (then *γ^i^* and *μ_T_^i^*) were assumed to be drawn from zero-truncated normal prior distributions *γ^i^* ~ *N*(*λ_γ_, σ_γ_*) and *μ_T_^i^* ~ *N*(*λ_μT_*, *σ_μ_T__*). In a sensitivity analysis we replaced the normal priors with gamma distributions *γ^i^* ~ Gamma(*α_γ_, β_γ_*) and *μ_T_^i^* ~ Gamma(*α_μ_T__, β_μ_T__*). All hyperparameters were assigned improper uniform prior distributions.

Particular attention was paid to the estimation of the natural mortality rate (*μ*), as this parameter was anticipated to influence the results of the analysis. Although all cohorts originated from similar geographical and socio-economic settings (being all from Western Europe), variations were observed in their recruitment years, as well as in their demographic characteristics (see Figures S1 and S2 in Supplement). To account for this and to incorporate uncertainty around the parameter value, *μ* was included as an estimated cohort-specific parameter. We used more informative priors on *μ* for the cohorts with available information about demographics (see Supplement for details).

A Hamiltonian Monte Carlo algorithm (HMC) was used to generate 10,000 samples from the posterior distributions of the parameters and the estimation was made separately for SP-TB and SN-TB cohorts. The 10,000 samples used for inference were obtained from 60,000 iterations of the HMC, burning the first 10,000 draws and only retaining every fifth element of the remaining 50,000 draws (thinning). Burn-in size and convergence were assessed by inspecting the trace plots of the estimated parameters, while the performed thinning was sufficient to reach negligible auto-correlation. The reported 95% credible intervals were obtained by computing the 2.5^th^ and 97.5^th^ percentiles of the parameters’ posterior distributions. Programming was done in R (v.3.5.1) using the package ‘rstan’ (v.2.17.3) and the associated data and code are publicly available via a Github repository (github.com/romain-ragonnet/tb_natural_history) (12, 13).

## Results

### Parameter estimates

Figure 3 and Figure 4 present the results of the parameter estimation for smear-positive TB and smear-negative TB, respectively. The detailed posterior distributions of the cohort-specific parameters are available in the Supplement. The posterior median estimates of TB-specific mortality rates were 0.390 year^−1^ (0.330-0.453, 95% credible interval) and 0.025 year^−1^ (0.016-0.036) for SP-TB and SN-TB patients, respectively. The estimates of self-recovery rates were 0.234 year^−1^ (0.178-0.294) and 0.148 year^−1^ (0.085-0.242) for SP-TB and SN-TB patients, respectively. We observed important variations across the different cohorts in relation to the estimated TB mortality of smear-positive TB patients, with rates ranging from 0.151 year^−1^ (0.137-0.166) for cohort #19 to 0.812 year^−1^ (0.712-0.920) for cohort #50.

**Figure 3.**
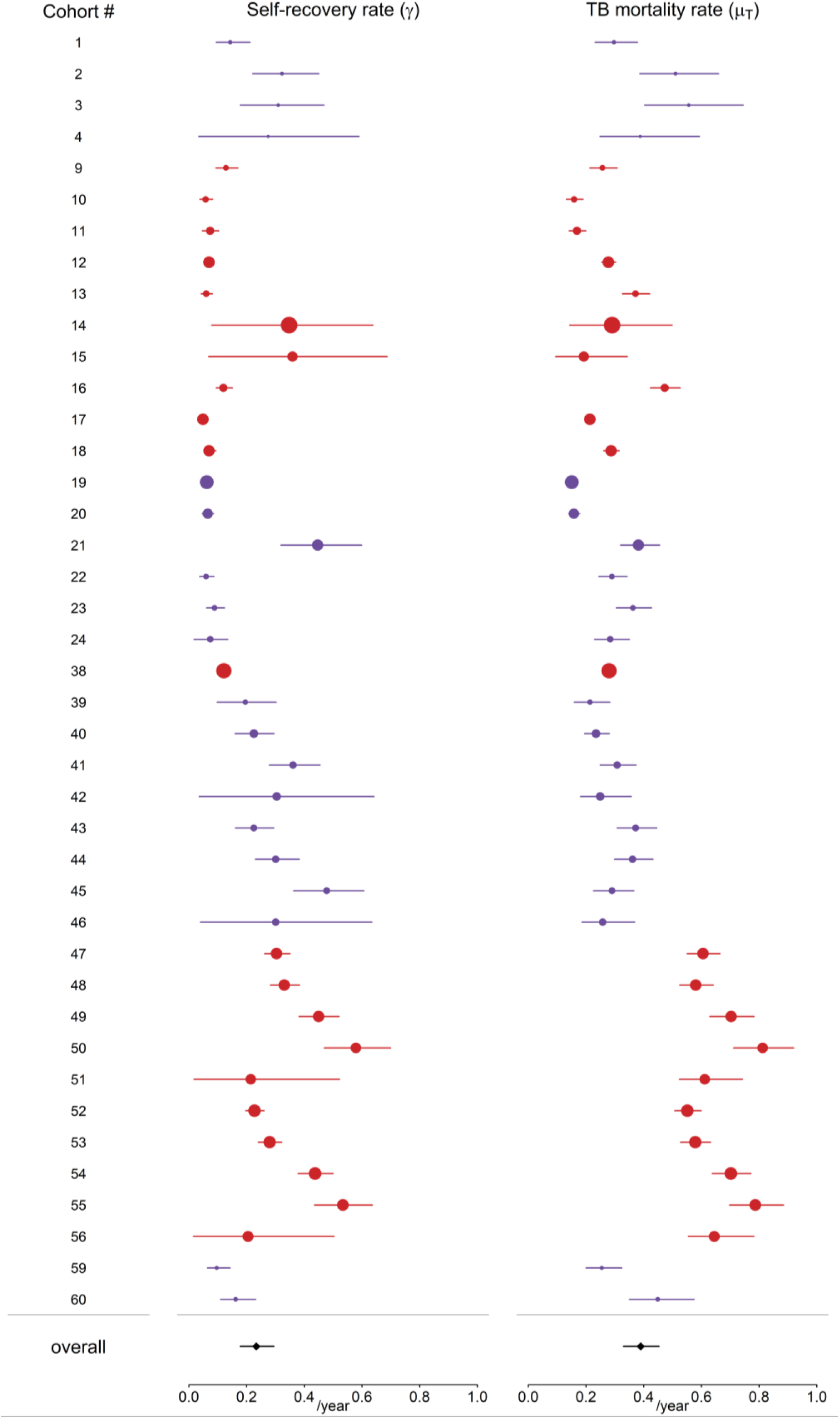
Parameter estimates for individuals with smear-positive TB. Results obtained with gaussian prior distributions. Dots represent the median estimates and horizontal bars represent the 95% credible intervals. The symbol areas are proportional to the cohort sizes. Colours represent sanatorium attendance status (purple: Yes, red: No).

**Figure 4.**
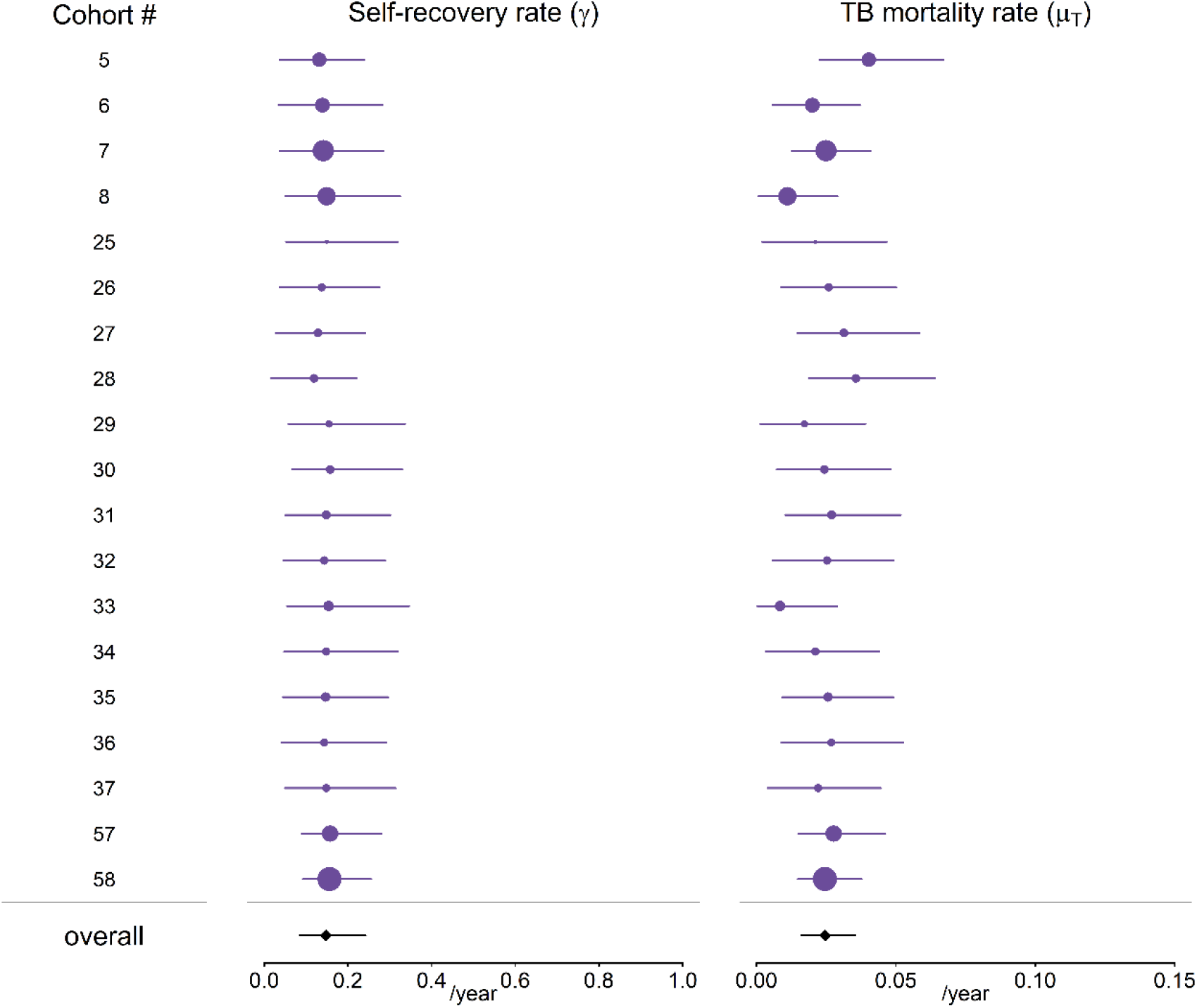
Parameter estimates for individuals with smear-negative TB. Results obtained with gaussian prior distributions. Dots represent the median estimates and horizontal bars represent the 95% credible intervals. The symbol areas are proportional to the cohort sizes. All smear-negative TB patients attended sanatoria.

The average disease duration and TB case fatality ratios in the absence of treatment also depend on the value of the natural mortality rate (*μ*). Table 1 presents their estimated values for various levels of natural mortality, with illustrative populations associated with such natural mortality rates. The average duration of active disease was obtained from 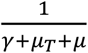 and the analytical expression of the case fatality ratio after *T* years was: 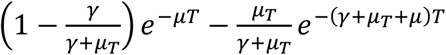 (see Supplement for details). The resulting estimates demonstrate the strong influence of natural mortality on disease duration and case fatality ratios, especially in individuals with SN-TB.

**Table 1.**
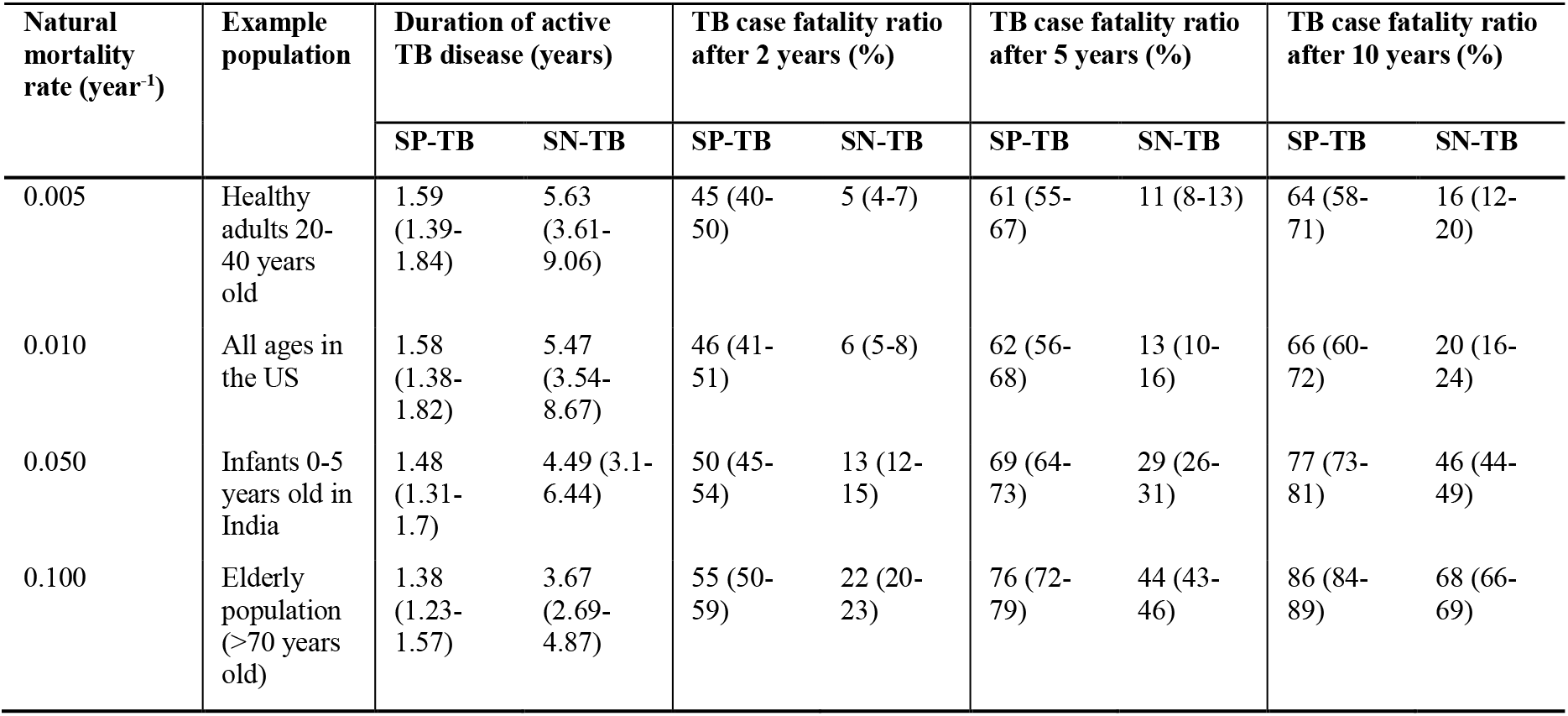
Estimated TB disease durations and case fatality ratios after 2, 5 and 10 years. Results are presented as median estimates and 95% simulation intervals.

| Natural mortality rate (year<sup>-1</sup>) | Example population | Duration of active TB disease (years) |  | TB case fatality ratio after 2 years (%) |  | TB case fatality ratio after 5 years (%) |  | TB case fatality ratio after 10 years (%) |  |
| --- | --- | --- | --- | --- | --- | --- | --- | --- | --- |
|  |  | SP-TB | SN-TB | SP-TB | SN-TB | SP-TB | SN-TB | SP-TB | SN-TB |
| 0.005 | Healthy adults 20-40 years old | 1.59<br>(1.39-1.84) | 5.63<br>(3.61-9.06) | 45 (40-50) | 5 (4-7) | 61 (55-67) | 11 (8-13) | 64 (58-71) | 16 (12-20) |
| 0.010 | All ages in the US | 1.58<br>(1.38-1.82) | 5.47<br>(3.54-8.67) | 46 (41-51) | 6 (5-8) | 62 (56-68) | 13 (10-16) | 66 (60-72) | 20 (16-24) |
| 0.050 | Infants 0-5 years old in India | 1.48<br>(1.31-1.7) | 4.49 (3.1-6.44) | 50 (45-54) | 13 (12-15) | 69 (64-73) | 29 (26-31) | 77 (73-81) | 46 (44-49) |
| 0.100 | Elderly population (>70 years old) | 1.38<br>(1.23-1.57) | 3.67<br>(2.69-4.87) | 55 (50-59) | 22 (20-23) | 76 (72-79) | 44 (43-46) | 86 (84-89) | 68 (66-69) |

Table 2 presents the estimates obtained for SP-TB patients when conducting the analysis separately according to the patients’ gender and depending on whether or not they attended a sanatorium. We observed no significant difference by gender in the self-recovery and TB mortality rates. However, sanatorium patients were found to have a significantly lower TB mortality rate than those who did not attend such TB care facilities.

**Table 2.**
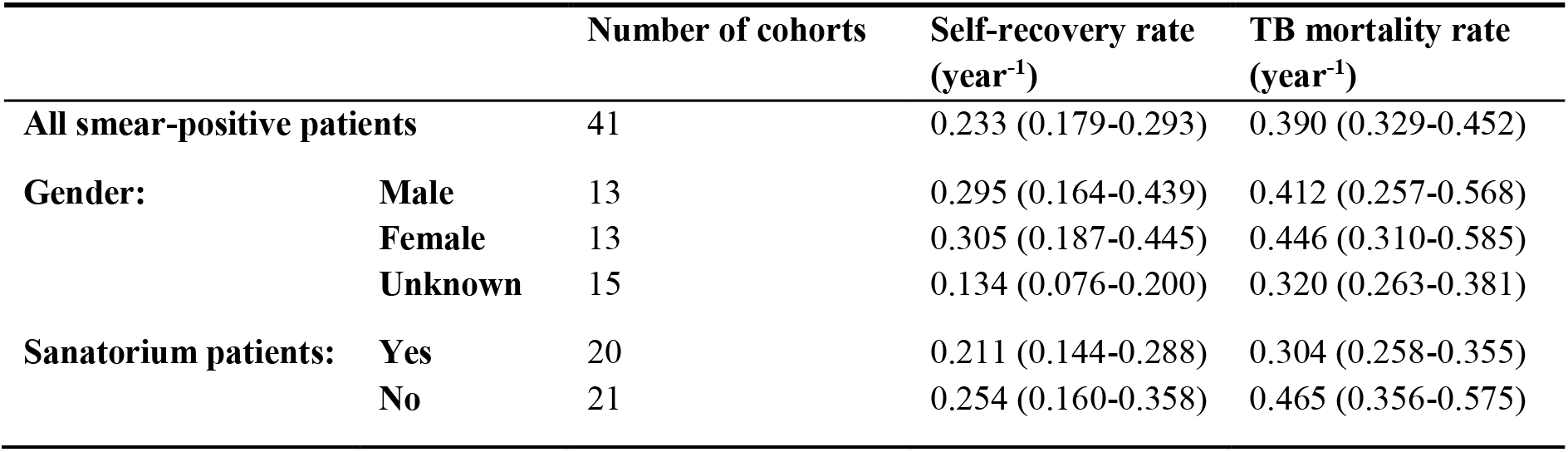
Parameter estimates for smear-positive patients with stratification by gender and sanatorium attendance status. Inference was performed separately for the different subgroups. Results are presented as median estimates and 95% simulation intervals.

|  |  | Number of cohorts | Self-recovery rate<br>(year<sup>-1</sup>) | TB mortality rate<br>(year<sup>-1</sup>) |
| --- | --- | --- | --- | --- |
| **All smear-positive patients** |  | 41 | 0.233 (0.179-0.293) | 0.390 (0.329-0.452) |
| **Gender:** | **Male** | 13 | 0.295 (0.164-0.439) | 0.412 (0.257-0.568) |
|  | **Female** | 13 | 0.305 (0.187-0.445) | 0.446 (0.310-0.585) |
|  | **Unknown** | 15 | 0.134 (0.076-0.200) | 0.320 (0.263-0.381) |
| **Sanatorium patients:** | **Yes** | 20 | 0.211 (0.144-0.288) | 0.304 (0.258-0.355) |
|  | **No** | 21 | 0.254 (0.160-0.358) | 0.465 (0.356-0.575) |

Comparison by gender and sanatorium attendance status was not possible for SN-TB patients, as all reported SN-TB patients attended sanatorium and the patients’ gender was known for only two SN-TB cohorts.

### Sensitivity analysis

The estimates obtained using gamma prior distributions for the cohort-specific parameters were very similar to estimates obtained in the primary analysis. In this sensitivity analysis, the posterior median estimates of TB-specific mortality rates were 0.386 year^−1^ (0.333-0.447) and 0.021 year^−1^ (0.015-0.031) for SP-TB and SN-TB patients, respectively. The estimates of self-recovery rates were 0.227 year^−1^ (0.179-0.291) and 0.109 year^−1^ (0.054-0.192) for SP-TB and SN-TB patients, respectively. Figures S5 and S6 (Supplement) provide the associated results by cohort.

## Discussion

This study is the first to produce quantitative estimates of disease-induced mortality and self-recovery rates for pulmonary TB patients. It suggests that TB mortality rates are around 15 times higher for SP-TB than for SN-TB patients, while self-recovery rates are comparable between the two categories of patients.

In order to compare our estimates to the parameter values previously employed, we conducted a systematic review of existing TB modelling studies and extracted the TB mortality and spontaneous recovery rates. The search strategy and selection criteria are described in Supplement and the results of the review are summarised in Figure 5. Interestingly, while most of the studies published after 2011 (11 out of 16, 69%) used Tiemersma and colleagues’ review to inform the natural history parameters, we observe important heterogeneity in the employed parameter values. This suggests that modellers have various interpretations of the findings of the previous review and that the rigorous approach used for estimation in the present analysis may allow for improvements and homogenisation in TB modelling methodology.

**Figure 5.**
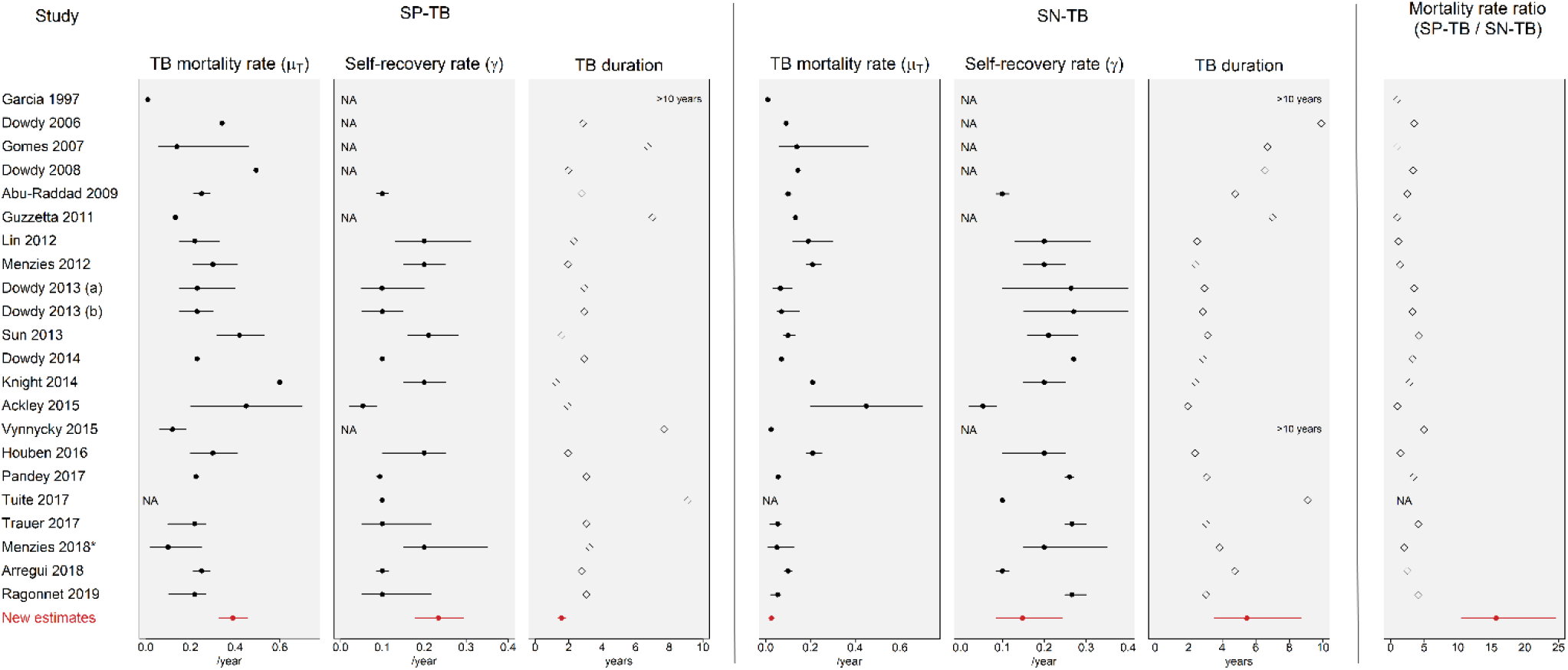
Comparison of our parameter estimates with previous TB modelling works. *Parameter values employed for the age category 0-15 years old. NA: parameter value not reported or process not simulated in the listed study. TB duration was estimated using a natural mortality rate of 0.01.

The dramatic difference of mortality rates between SP-TB and SN-TB patients was largely underestimated in previous TB modelling studies, as most of them parameterised models based on the ratio of 3.3 between the 10-year cumulative fatality of SP-TB and SN-TB (14, 15). Our findings highlight the danger represented by employing risk ratios in place of rate ratios to inform systems that are parameterised with rates. While the current study raises important concerns about the accuracy of previous modelling-based estimates, further work will be required to assess which model applications would be significantly affected by our new findings. Nevertheless, one can anticipate that the predicted effect of interventions linked to case detection are highly likely to be sensitive to natural history parameters, as the impact of such interventions strongly depends on the individuals’ prognosis in absence of treatment.

The current study is also likely to have critical implications for the estimation of local and global TB incidence and mortality (6). In particular, using our new estimates to re-estimate disease burden indicators in places with poor case detection may result in marked discrepancies with the previously reported estimates, as TB natural history is expected to play a major role in such settings.

The hierarchical approach allowed us to highlight the marked heterogeneity in cohort-specific estimates of TB mortality of smear-positive patients. This may be explained by the different diagnostic methods employed to identify TB and the various types of care used in the different cohorts. For smear positive cases, we were able to examine the impact of some factors. In particular, we found that patients who did not attend sanatoria had significantly higher mortality rates (50% increase) than those treated in such facilities. In contrast, no difference by sex was noted in the prognosis of untreated pulmonary TB patients.

An important strength of this study is that we were able to distinguish mortality that is specifically induced by TB from that linked to natural causes. That is, the rates presented in this report should be interpreted as TB-specific mortality rates that have to be added to natural mortality in order to capture the overall death rate. The main advantage of this distinction is that our estimates could be applied directly to any settings without the need for any further adjustment, such that the TB-specific mortality rate can be simply added to the setting-specific natural mortality rate.

Our study includes several limitations. First, we were unable to provide estimates for extrapulmonary TB patients, as the required data were not available. For the same reason, the analysis could not be performed separately for male versus female patients or for sanatorium versus non-sanatorium patients in the case of smear-negative patients. In addition, no age-specific estimates could be obtained due to the limited number of cohorts with information about age-distribution. Also, it is to be noted that most patients included in the analysis were aged 15 and over, such that the rates presented here are representative of adult individuals with pulmonary TB.

Another limitation is that the diagnostics used in these historical cohorts differ substantially from current state of the art diagnostics, especially in the case of SN-TB patients. As culture-based diagnostics were only invented in the early 1930s, many SN-TB patients were probably diagnosed clinically or on the basis of X-ray. Moreover, the classification into SP-TB or SN-TB patients may be unclear, as it depends very much on the quality of the microscopy, the number of slides examined or how sputum was collected. A final caveat is a possible lack of representativeness of the cohorts included, as undiagnosed patients by definition were not considered. Moreover, our finding that patients treated in sanatoria had a better prognosis is difficult to interpret, as there may be selection bias driven by factors influencing the decision of whether to admit TB patients to these institutions (16, 17).

Unfortunately, most of the limitations listed above could not be addressed by future studies, as it would be unethical to follow up TB patients without providing them with treatment. Accordingly, no other data than those used to inform the present analysis and previously reported by Tiemersma and colleagues could be used to estimate TB mortality and self-recovery rates (9).

In conclusion, this study presents detailed estimates for the parameters that govern the natural history of pulmonary TB patients. It highlights that the gap between smear-positive and smear-negative TB patients in terms of mortality rates is much higher than previously thought. The parameter values reported in this study should improve the accuracy of disease burden estimations and make future TB modelling works more reliable.

## Supporting information

Cohort profiles smear-negative

Cohort profiles smear-positive

## Acknowledgments

We would like to thank Dr Nico Nagelkerke for providing copies of the manuscripts and reports used to extract the data on TB patients’ prognosis. We are also grateful for his advice regarding the interpretation of the findings.

## References

1. Jones WHS. Hippocrates, Vol. I. Cambridge, MA: Harvard University Press. 1923.

2. WHO. Global TB Report 2018. 2018.

3. Organization WH. Ethics guidance for the implementation of the end tb strategy. 2017.

4. Storla DG, Yimer S, Bjune GA. A systematic review of delay in the diagnosis and treatment of tuberculosis. BMC Public Health. 2008;8:15.

5. Dheda K, Barry CE, 3rd, Maartens G. Tuberculosis. Lancet. 2016;387(10024):1211–26.

6. Glaziou P, Dodd PJ, Zignol M, Sismanidis C, Floyd K. Methods used by WHO to estimate the global burden of TB disease. 2018.

7. Fund TG. Detailed Explanation of the Allocation Methodology 2017-2019. 2017.

8. Dowdy DW, Dye C, Cohen T. Data needs for evidence-based decisions: a tuberculosis modeler’s ‘wish list’. Int J Tuberc Lung Dis. 2013;17(7):866–77.

9. Tiemersma EW, van der Werf MJ, Borgdorff MW, Williams BG, Nagelkerke NJ. Natural history of tuberculosis: duration and fatality of untreated pulmonary tuberculosis in HIV negative patients: a systematic review. PLoS One. 2011;6(4):e17601.

10. Bank TW. Life expectancy at birth 2019 [Available from: https://data.worldbank.org/indicator/sp.dyn.le00.in.

11. Gelman A, Hill J. Data Analysis Using Regression and Multilevel/Hierarchical Models: Cambridge University Press; 2009.

12. Team RC. R: A Language and Environment for Statistical Computing. 2019.

13. Team SD. RStan: the R interface to Stan. 2018.

14. Ragonnet R, Underwood F, Doan T, Rafai E, Trauer J, McBryde E. Strategic Planning for Tuberculosis Control in the Republic of Fiji. Trop Med Infect Dis. 2019;4(2).

15. Trauer JM, Denholm JT, Waseem S, Ragonnet R, McBryde ES. Scenario Analysis for Programmatic Tuberculosis Control in Western Province, Papua New Guinea. Am J Epidemiol. 2016;183(12):1138–48.

16. McCarthy OR. The key to the sanatoria. J R Soc Med. 2001;94(8):413–7.

17. Williams LR, Hill AM. A Study of what Tuberculosis Patients Experience Prior to their Admission to a Sanatorium. The New England Journal of Medicine. 1929;201(2):82–91.

