## Supplement for "Revisiting the Natural History of Pulmonary Tuberculosis: a Bayesian Estimation of Natural Recovery and Mortality rates"

#### Supplementary Appendix

Romain Ragonnet, Jennifer A. Flegg, Samuel L. Brilleman, Edine Tiemersma, Yayehirad A. Melsew, Emma S. McBryde, James M. Trauer

#### Contents

**1 Age distribution of TB patients 1**

**2 Cohort sizes and times of patient diagnosis 1**

**3 Estimation of the natural mortality rate $\mu$ 2**

**4 Prior distributions 4**

**5 Likelihood calculation 4**

**6 Results of sensitivity analysis using a gamma distribution as prior 6**

### 1 Age distribution of TB patients

The age-distribution of the TB patients was available for 20 cohorts. They are represented in Figure S1. Some studies reporting on several cohorts did not provide the age distribution by cohort. We used the available aggregated data to inform the analysis for these cohorts.

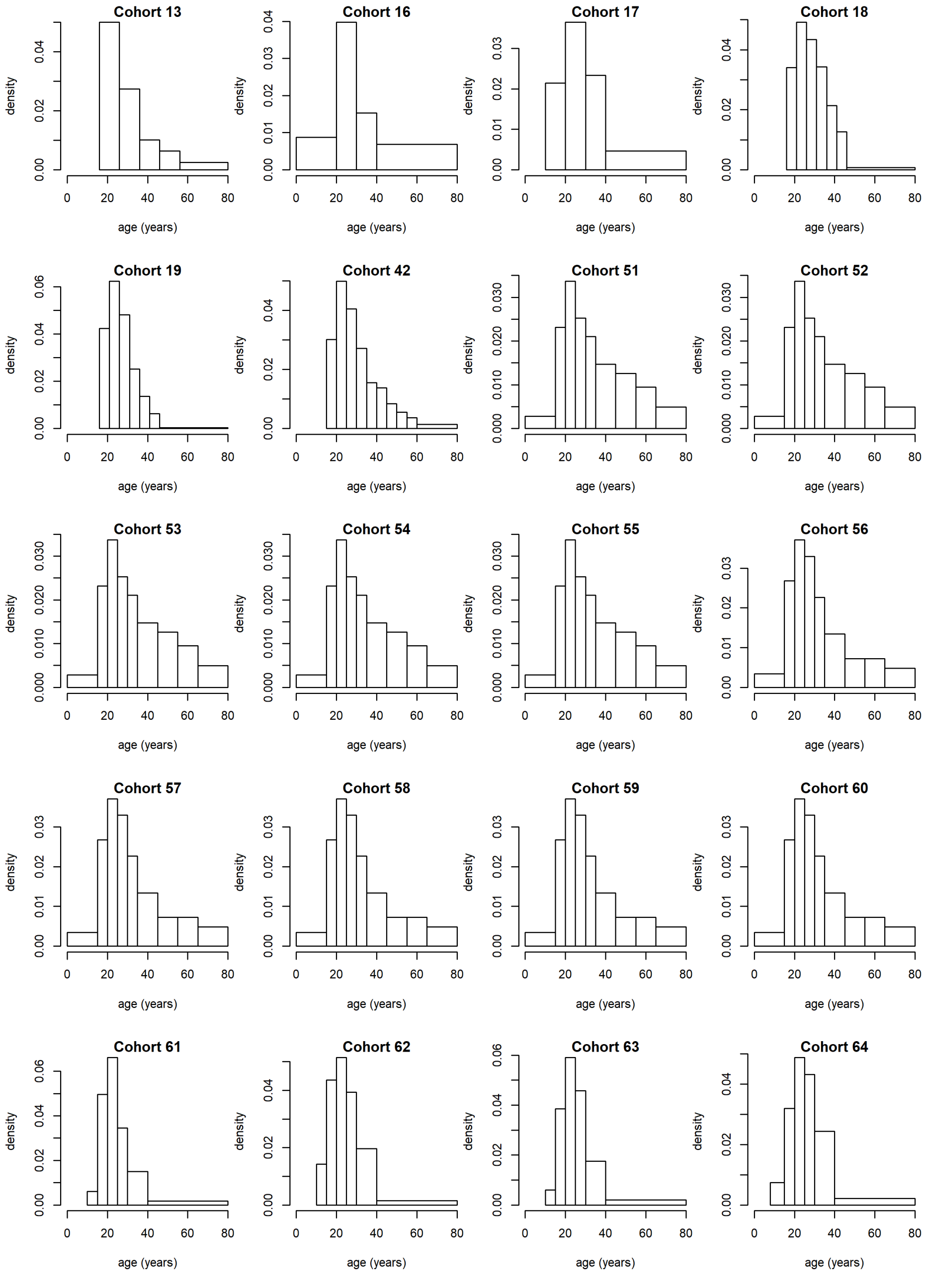

**Figure S1. Age distribution of TB patients**

### 2 Cohort sizes and times of patient diagnosis

The figure below presents the cohort sizes and the times of patient recruitment for the 60 cohorts included in the analysis.

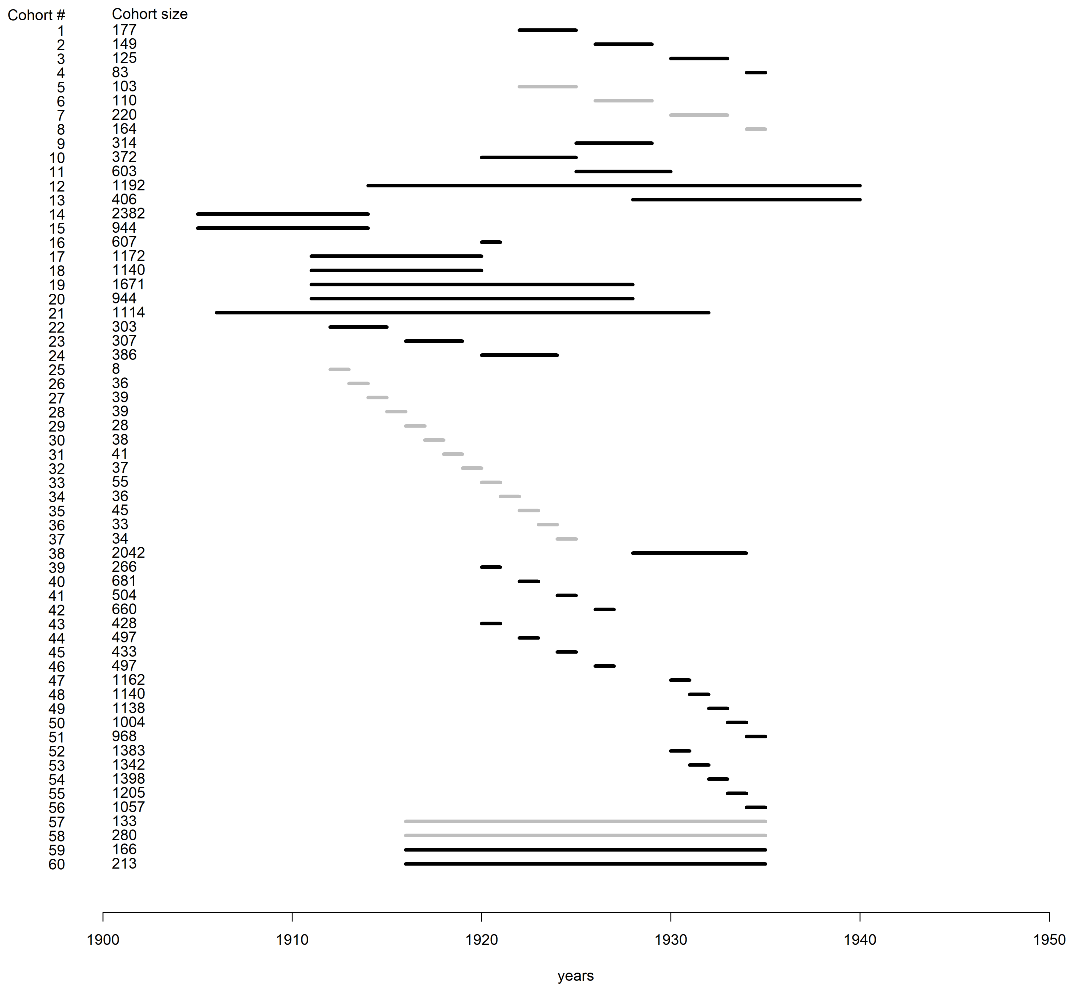


**Figure S2. Cohort sizes and years of patient diagnosis.**
Cohort of smear-positive TB patients are represented in black while cohorts of smear-negative TB patients are represented in grey.

### 3 Estimation of the natural mortality rate $\mu$

The natural mortality rates were estimated during the MCMC simulation as cohort-specific parameters, along with the other model parameters. The priors used for these rates were also cohort-specific and were gamma distributions parameterised according to the cohort demographic characteristics, as described below.

First, we determined the mean of the gamma distributions as follows: when an age-distribution was available for a cohort, we used the weighted average of the age-specific (and sex-specific when patients sex was known) mortality rates obtained from the England and Wales life tables provided by the UK Office for National Statistics. We consider the year that corresponds to the average recruitment year of the cohort. These data cover the period 1841-2016 and are accessible from **here**.

Figure S3 represents these data for years 1905-1940 when averaging male and female rates.

When the age-distribution was not provided for a cohort, we used 0.01 as mean for the gamma distribution, as this value is representative of the average mortality rate of the

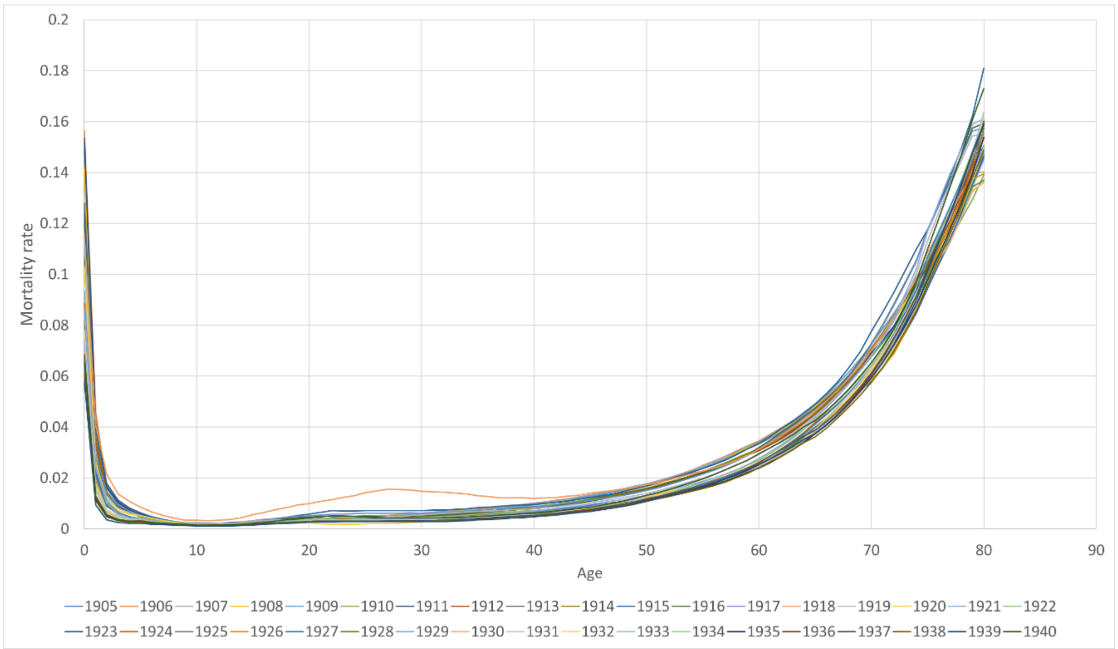


**Figure S3. Mortality rate by age for years 1905-1940.**

adult population during the first half of the 20$^{th}$ century (see Figure S3).

The standard deviations of the gamma distributions were set to 0.001 for cohorts with known age-distribution and 0.002 for cohorts with unknown age-distribution. For illustration, Figure S4 represents the density of the prior distributions used for the natural mortality rate of Cohort #1 (with unknown age-distribution) and Cohort #47 (known age-distribution).

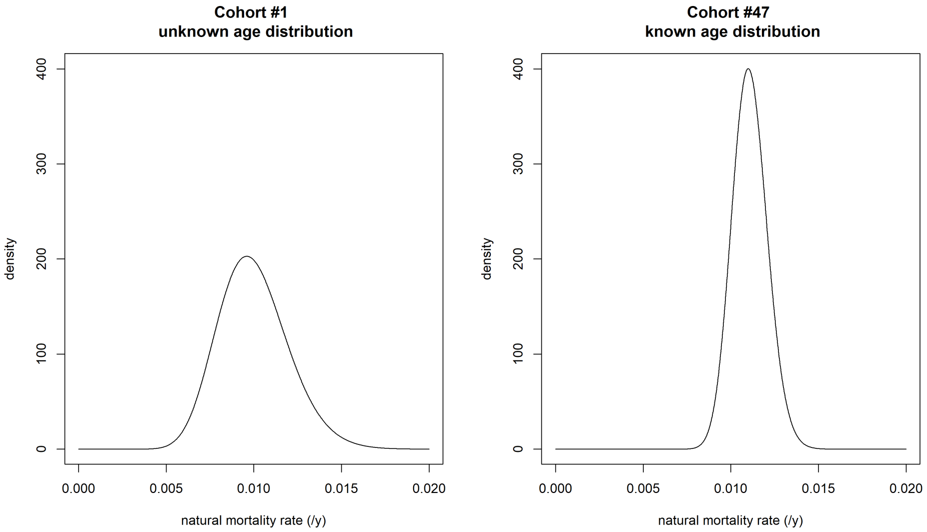


**Figure S4. Prior distributions used for the natural mortality rates of Cohort #1 and Cohort #47.**

### 4 Prior distributions

Table S1 lists the prior probability distributions associated with the different parameters. In our Bayesian hierarchical approach, the cohort-specific parameters $\mu_T^i$ and $\gamma^i$ are associated with priors that are not-cohort specific: $N(\lambda_{\mu_T}, \sigma_{\mu_T})$ and $N(\lambda_\gamma, \sigma_\gamma)$, respectively. This means that although we allow for these parameters to vary by cohort, we assume that the cohort-specific parameters are still related by a common distribution.

| parameter | definition | distribution | details |
|---|---|---|---|
| $\mu^i$ | natural mortality rate of cohort $i$ | $Gamma(\alpha^i, \beta^i)$ | see Section 3 for calculation of $\alpha^i$ and $\beta^i$ |
| $\mu_T^i$ | TB-induced mortality rate of cohort $i$ | $N(\lambda_{\mu_T}, \sigma_{\mu_T})$<br>$Gamma(\alpha_{\mu_T}, \beta_{\mu_T})$ | ← main analysis<br>← sensitivity analysis |
| $\gamma^i$ | self recovery rate of cohort $i$ | $N(\lambda_\gamma, \sigma_\gamma)$<br>$Gamma(\alpha_\gamma, \beta_\gamma)$ | ← main analysis<br>← sensitivity analysis |
| $\lambda_{\mu_T}, \sigma_{\mu_T}$, $\alpha_{\mu_T}, \beta_{\mu_T}$, $\lambda_\gamma, \sigma_\gamma$, $\alpha_\gamma, \beta_\gamma$ | hyperparameters | improper uniform | |

**Table S1. Prior distributions associated with the parameters.**

### 5 Likelihood calculation

As described in the main text, the likelihood of the observed data (numbers of deaths $d_k^i$ occurring during the time-interval $I_k^i$) is obtained by evaluating the density function of a binomial distribution $B(n_k^i, p_k^i)$, where $n_k^i$ is the number of people still alive at the beginning of each interval $I_k^i$, and $p_k^i$ is the individual probability of death within time-interval $I_k^i$, provided that the individual was alive at the beginning of $I_k^i$. Here, we describe how the probabilities $p_k^i$ are obtained analytically using the Markov process.

To calculate $p_k^i$, we need the expression of $p(t_1, t_2)$ for any $t_2 > t_1 \geq 0$, representing the probability of death by time $t_2$ provided that an individual was alive at time $t_1$. The transition matrix associated with the model represented in Figure 2 with state order ("Active TB", "Recovered", "Death") is:

$$Q = \begin{pmatrix} -(\gamma + \mu + \mu_T) & \gamma & \mu + \mu_T \\ 0 & -\mu & \mu \\ 0 & 0 & 0 \end{pmatrix}.$$

To simplify notations, the states ("Active TB", "Recovered", "Death") will now be referred to as ("T", "R", "D").

The marginal distribution of the Markov process after time $t$ is given by:

$$P(t) = P(0)e^{Qt},$$

where $P(t)$ is a row vector containing the state probabilities at time $t$. In our case, $P(0) = (1, 0, 0)$.

The eigenvalues of $Q$ are $(-(\gamma + \mu + \mu_T), -\mu, 0)$. Therefore, $Q$ is diagonalisable, as $\mu > 0$ and $\gamma + \mu_T > 0$ imply that the three eigenvalues are distinct. We can calculate $e^{Qt}$ using the diagonalised decomposition $Q = UDU^{-1}$ and therefore:

$$e^{Qt} = Ue^{Dt}U^{-1},$$

with

$$e^{Dt} = \begin{pmatrix} e^{-(\gamma+\mu+\mu_T)t} & 0 & 0 \\ 0 & e^{-\mu t} & 0 \\ 0 & 0 & 1 \end{pmatrix}.$$

After calculation, we obtain:

$$e^{Qt} = \begin{pmatrix} e^{-(\gamma+\mu+\mu_T)t} & \frac{\gamma}{\gamma+\mu_T}(e^{-\mu t} - e^{-(\gamma+\mu+\mu_T)t}) & (1-\frac{\gamma}{\gamma+\mu_T})e^{-\mu t} - \frac{\mu_T}{\gamma+\mu_T}e^{-(\gamma+\mu+\mu_T)t} \\ 0 & e^{-\mu t} & 1-e^{-\mu t} \\ 0 & 0 & 1 \end{pmatrix} \tag{1}$$

and therefore

$$P(t)^T = \begin{pmatrix} e^{-(\gamma+\mu+\mu_T)t} \\ \frac{\gamma}{\gamma+\mu_T}(e^{-\mu t} - e^{-(\gamma+\mu+\mu_T)t}) \\ (1-\frac{\gamma}{\gamma+\mu_T})e^{-\mu t} - \frac{\mu_T}{\gamma+\mu_T}e^{-(\gamma+\mu+\mu_T)t} \end{pmatrix}. \tag{2}$$

Note that the third component of $P(t)$ represents the case fatality ratio after $t$ years and was used to populate Table 1 in the main text.

For our observed data, as we know that the individual is alive at time $t_1$, the active Markov state at time $t_1$ could not be "D". Then, the probability of death before $t_2$ is given by:

$$p(t_1,t_2) = \mathbb{P}\big(\{S_2 = \text{“D”}\} | \{S_1 = \text{“T”}\} \cup \{S_1 = \text{“R”}\}\big),$$

where $S_i$ denotes the active state of the system at time $t_i$. Using Bayes' theorem, we obtain:

$$p(t_1,t_2) = \frac{\mathbb{P}\Big(\{S_2 = \text{“D”}\} \cap \{\{S_1 = \text{“T”}\} \cup \{S_1 = \text{“R”}\}\}\Big)}{\mathbb{P}\Big(\{S_1 = \text{“T”}\} \cup \{S_1 = \text{“R”}\}\Big)}.$$

As $\{S_1 = \text{“T”}\}$ and $\{S_1 = \text{“R”}\}$ are mutually exclusive, we can use the following decomposition:

$$p(t_1,t_2) = \frac{\mathbb{P}\big(\{S_1 = \text{“T”}\} \cap \{S_2 = \text{“D”}\}\big) + \mathbb{P}\big(\{S_1 = \text{“R”}\} \cap \{S_2 = \text{“D”}\}\big)}{\mathbb{P}\big(\{S_1 = \text{“T”}\}\big) + \mathbb{P}\big(\{S_1 = \text{“R”}\}\big)}.$$

Using Bayes' theorem again,

$$\mathbb{P}\big(\{S_1 = \text{“T”}\} \cap \{S_2 = \text{“D”}\}\big) = \mathbb{P}\big(\{S_1 = \text{“T”}\}\big) \times \mathbb{P}\big(\{S_2 = \text{“D”}\}|\{S_1 = \text{“T”}\}\big)$$

and

$$\mathbb{P}\big(\{S_1 = \text{“R”}\} \cap \{S_2 = \text{“D”}\}\big) = \mathbb{P}\big(\{S_1 = \text{“R”}\}\big) \times \mathbb{P}\big(\{S_2 = \text{“D”}\}|\{S_1 = \text{“R”}\}\big).$$

We can now come back to the matrix notations and note, for example, that

$$\mathbb{P}\big(\{S_1 = \text{“T”}\}\big) = P(t_1)[1],$$

where $X[i]$ denotes the $i^{\text{th}}$ component of vector $X$.

Similarly,

$$\mathbb{P}\big(\{S_2 = \text{“D”}\}|\{S_1 = \text{“T”}\}\big) = (1,0,0)e^{Q(t_2-t_1)}(0,0,1)^T.$$

Finally, the analytical expression of $p(t_1,t_2)$ using matrix notations is:

$$p(t_1,t_2)=\frac{P(t_1)[1]\times(1,0,0)e^{Q(t_2-t_1)}(0,0,1)^T+P(t_1)[2]\times(0,1,0)e^{Q(t_2-t_1)}(0,0,1)^T}{P(t_1)[1]+P(t_1)[2]}.$$

Using the expression of $e^{Qt}$ and $P(t)$ found in equations 1 and 2, we have:

$$p(t_1,t_2)=\frac{\alpha(1-\frac{\gamma}{\gamma+\mu_T}e^{-\mu(t_2-t_1)}-\frac{\mu_T}{\gamma+\mu_T}e^{-(\gamma+\mu+\mu_T)(t_2-t_1)})+\beta(1-e^{-\mu(t_2-t_1)})}{\alpha+\beta},$$

where

$$\alpha=e^{-(\gamma+\mu+\mu_T)t_1}$$

and

$$\beta=\frac{\gamma}{\gamma+\mu_T}(e^{-\mu t_1}-e^{-(\gamma+\mu+\mu_T)t_1}).$$

If $\Delta t_k^i$ denotes the length of the interval $I_k^i$ and $t_k^i$ denotes the starting time of $I_k^i$, we obtain:

$$p_k^i=p(t_k^i,t_k^i+\Delta t_k^i)=\frac{\alpha_k^i(1-\frac{\gamma}{\gamma+\mu_T}e^{-\mu\Delta t_k^i}-\frac{\mu_T}{\gamma+\mu_T}e^{-(\gamma+\mu+\mu_T)\Delta t_k^i})+\beta_k^i(1-e^{-\mu\Delta t_k^i})}{\alpha_k^i+\beta_k^i}, \quad (3)$$

where

$$\alpha_k^i=e^{-(\gamma+\mu+\mu_T)t_k^i}$$

and

$$\beta_k^i=\frac{\gamma}{\gamma+\mu_T}(e^{-\mu t_k^i}-e^{-(\gamma+\mu+\mu_T)t_k^i}).$$

To obtain the likelihood function, we first evaluate $p_k^i$ for each time interval of each cohort. We remind that the quantities $t_k^i$ and $\Delta t_k^i$ are observed from data, while the parameters $\gamma$, $\mu_T$ and $\mu$ are estimated. Let's denote $L_k^i$ the likelihood of drawing $d_k^i$ deaths from the binomial distribution $B(n_k^i,p_k^i)$ ($d_k^i$ and $n_k^i$ being observed from data). The overall likelihood of the observed data is finally obtained by multiplying all the quantities $L_k^i$ together:

$$L=\prod_{i,k}L_k^i.$$

Figure S5 presents an overview of the Bayesian approach used to estimate the natural history parameters.

### 6 Results of sensitivity analysis using a gamma distribution as prior

Figures S6 and S7 present the results of the sensitivity analysis considering gamma distributions instead of normal distributions for the priors associated with the estimated parameters $\gamma^i$ and $\mu_T^i$.

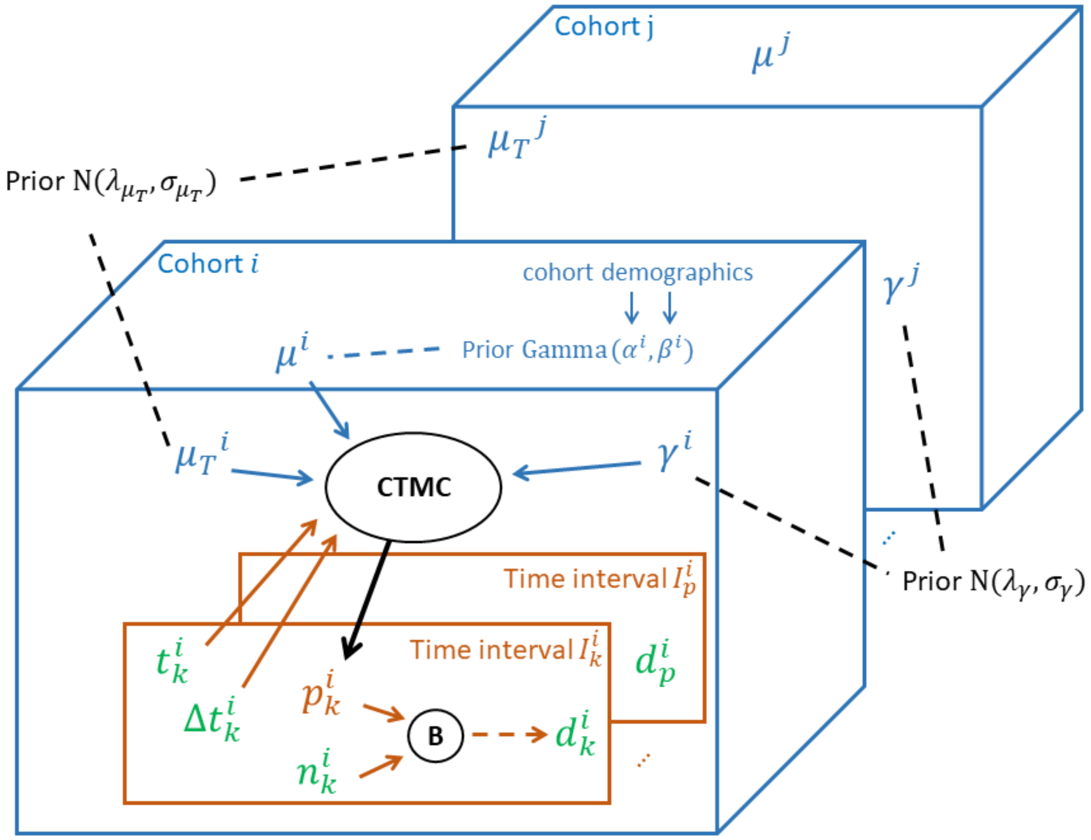


**Figure S5. Overview of the estimation framework.**
**CTMC**: Continuous-time Markov Chain. **B**: Binomial process. The blue boxes represent the different cohorts. The orange rectangles represent the different time-intervals for which data are available for a cohort. The variables written in green are those observed from data. Solid arrows represent deterministic dependencies while dashed arrows represent stochastic dependencies.

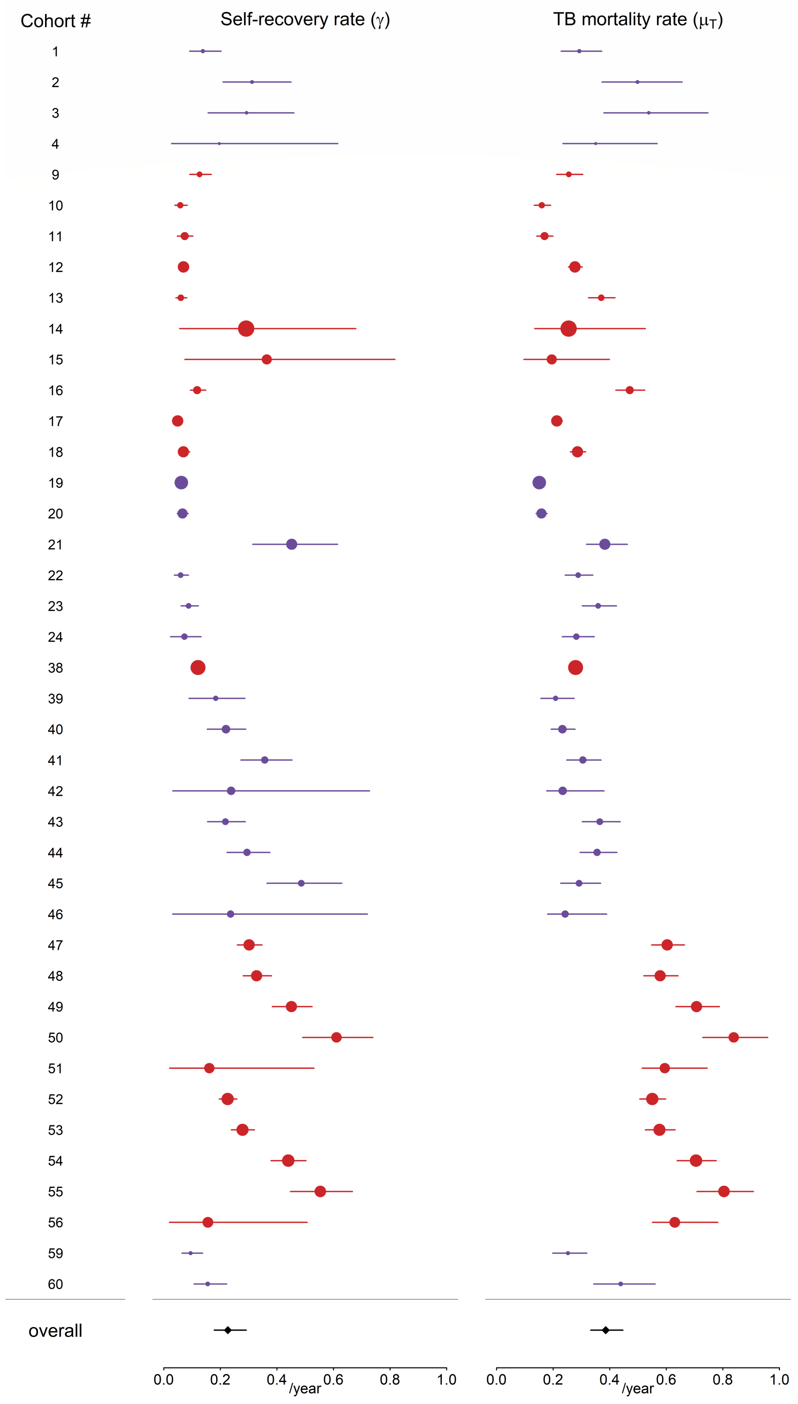


**Figure S6. Parameter estimates for individuals with smear-positive TB when using gamma priors.**
Dots represent the median estimates and horizontal bars represent the 95% credible intervals. The symbol areas are proportional to the cohort sizes. Colours represent sanatorium attendance status (purple: Yes, red: No).

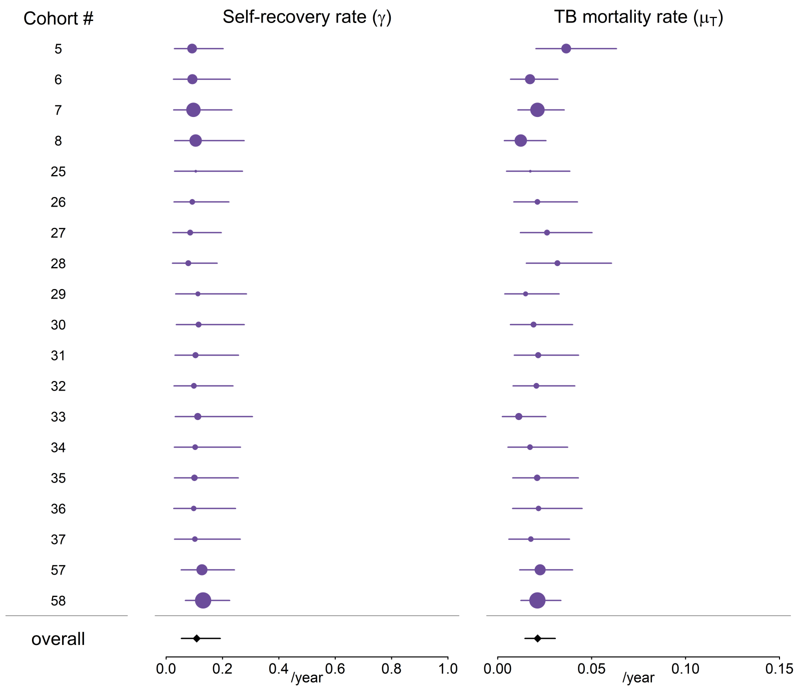


**Figure S7. Parameter estimates for individuals with smear-negative TB when using gamma priors.**
Dots represent the median estimates and horizontal bars represent the 95% credible intervals. The symbol areas are proportional to the cohort sizes. Colours represent sanatorium attendance status (purple: Yes, red: No).
