## Supplementary material for "Revisiting the Natural History of Pulmonary Tuberculosis: a Bayesian Estimation of Natural Recovery and Mortality rates": Cohort profiles smear-negative

#### Cohort profiles of smear-negative TB patients

The cohort profiles below present the main characteristics of the cohorts used in our analysis (top panel), the data points with posterior predictive curves (central panel) and the posterior distributions of the cohort-specific parameters $\gamma$ and $\mu_T$ (bottom panels). In the central panel, the data are represented with red diamonds and we show model realisations associated with 100 randomly selected posterior samples obtained from the MCMC simulation (blue lines).

### Cohort 5

Author: Baart De La Faille

Publication year: 1939

Patient recruitment years: 1922-1925

Cohort size: 103

Location: The Netherlands

Type of TB (smear-status): negative

Details: TB cases hospitalized in the Sanatorium "Berg en Bosch".

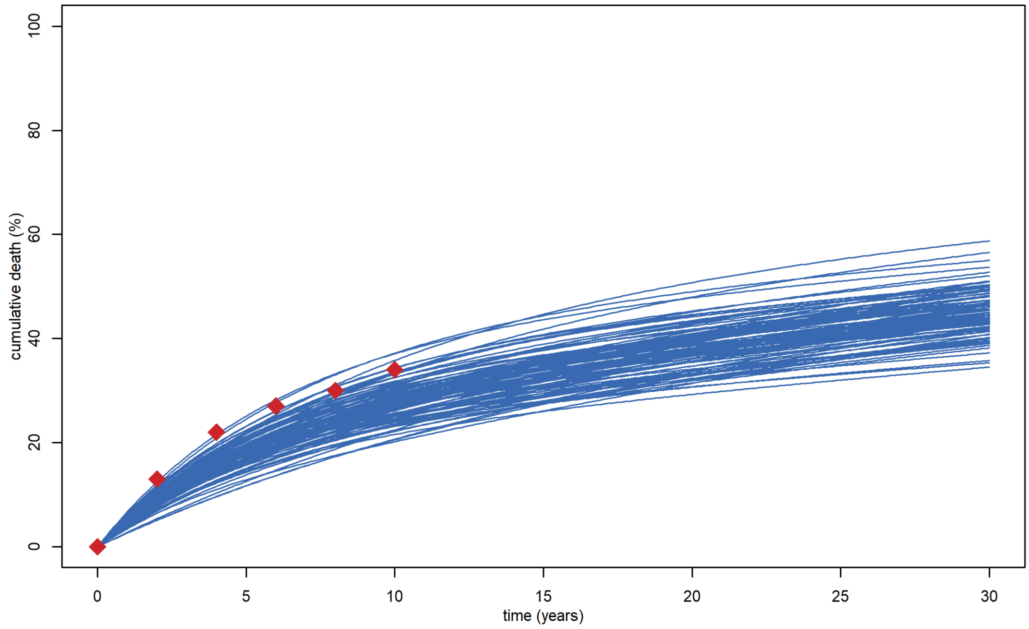


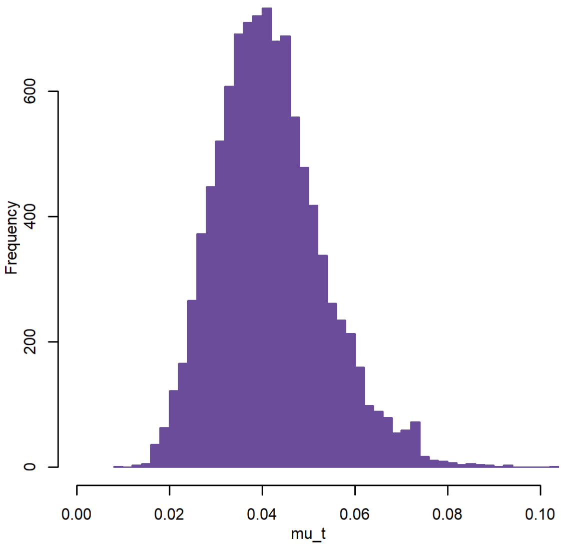


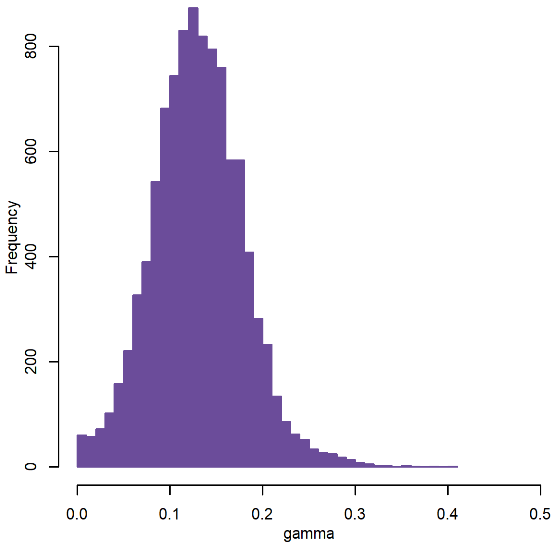

### Cohort 6

Author: Baart De La Faille
Publication year: 1939
Patient recruitment years: 1926-1929
Cohort size: 110

Location: The Netherlands
Type of TB (smear-status): negative
Details: TB cases hospitalized in the Sanatorium "Berg en Bosch".

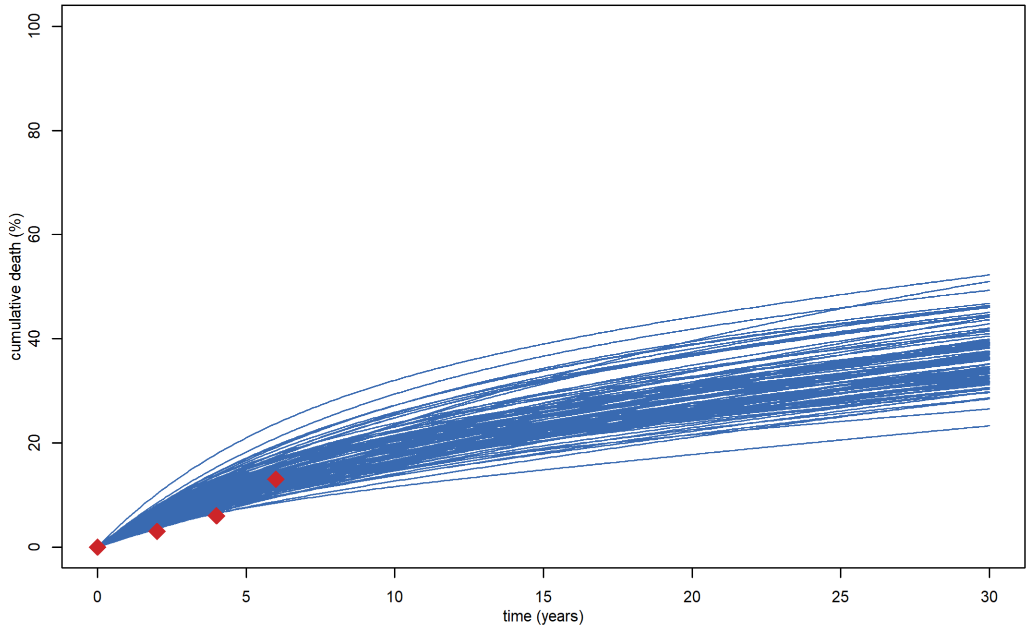


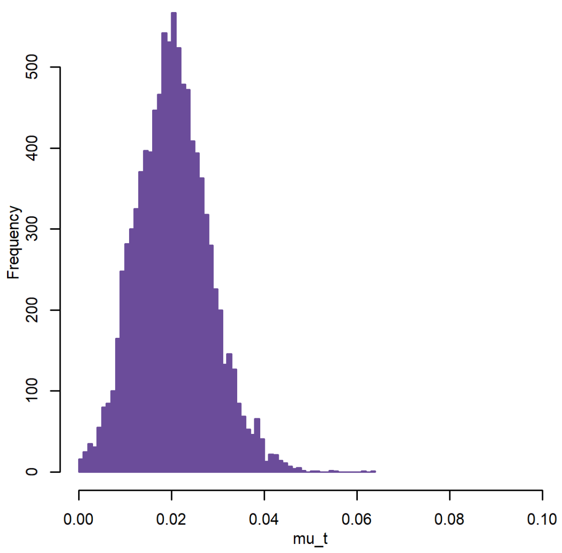


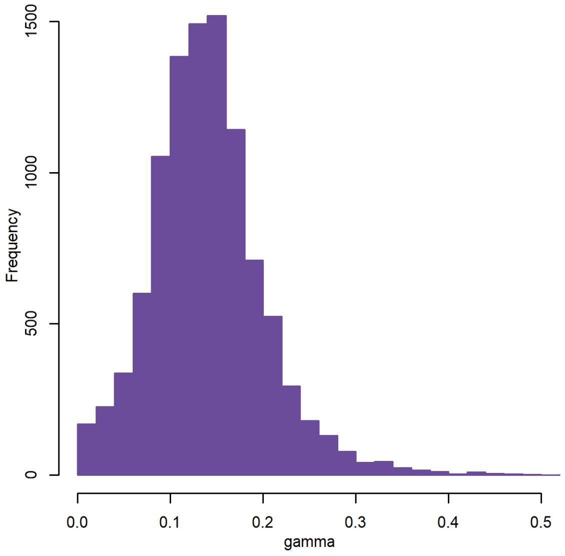

### Cohort 7

Author: Baart De La Faille

Publication year: 1939

Patient recruitment years: 1930-1933

Cohort size: 220

Location: The Netherlands

Type of TB (smear-status): negative

Details: TB cases hospitalized in the Sanatorium "Berg en Bosch".

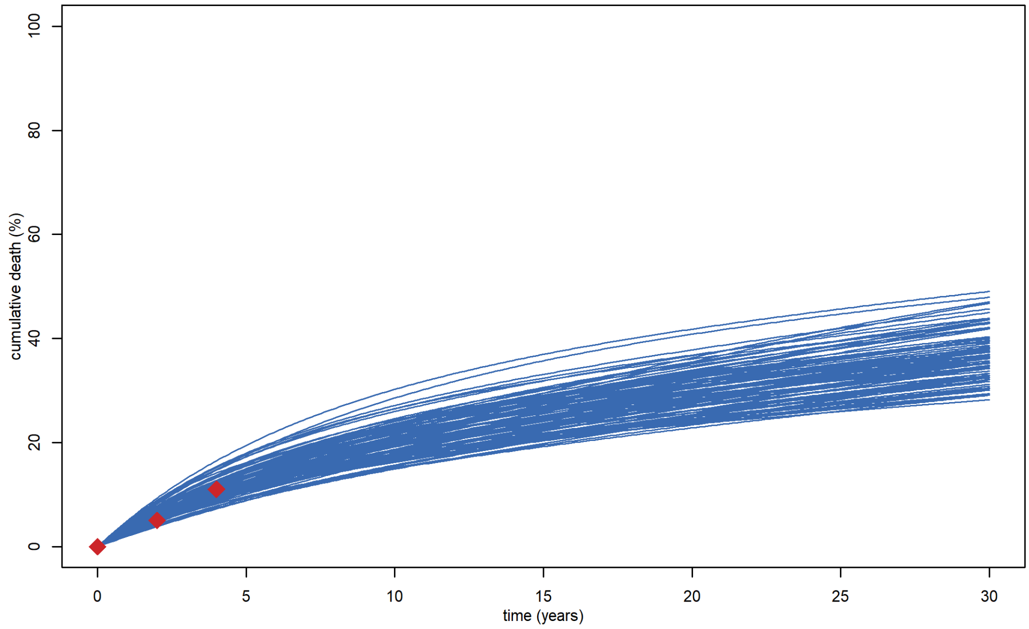


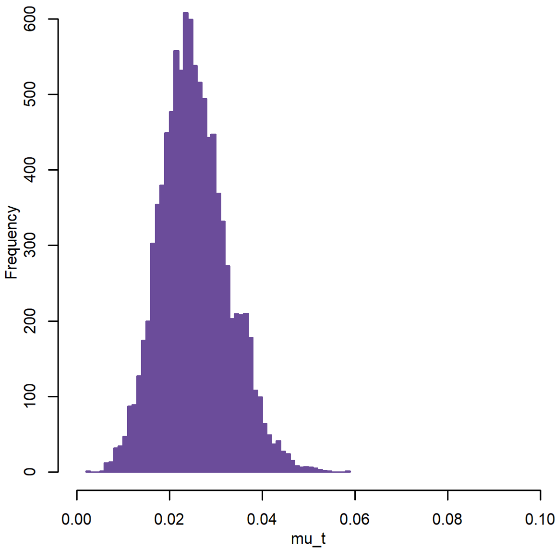


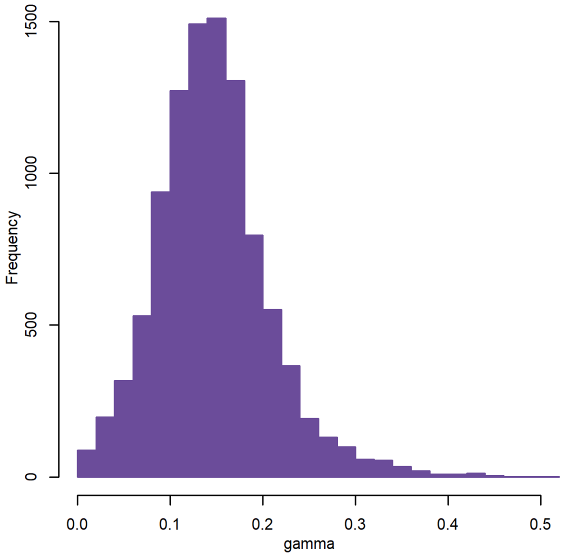

### Cohort 8

Author: Baart De La Faille

Publication year: 1939

Patient recruitment years: 1934-1935

Cohort size: 164

Location: The Netherlands

Type of TB (smear-status): negative

Details: TB cases hospitalized in the Sanatorium "Berg en Bosch".

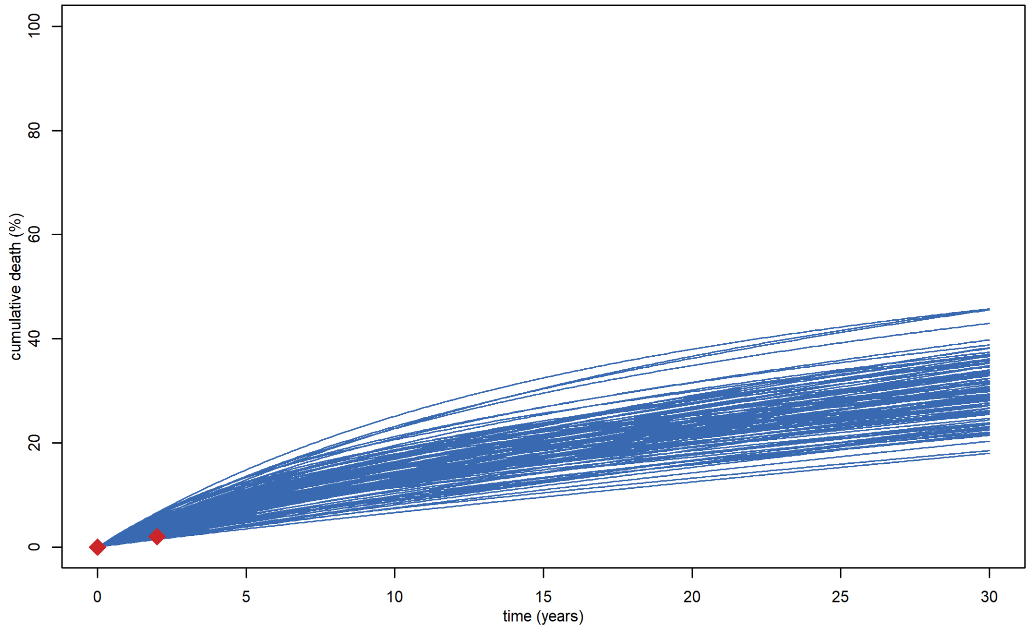


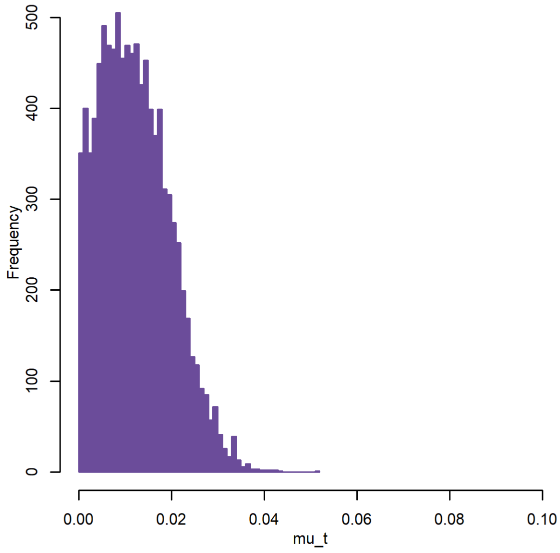


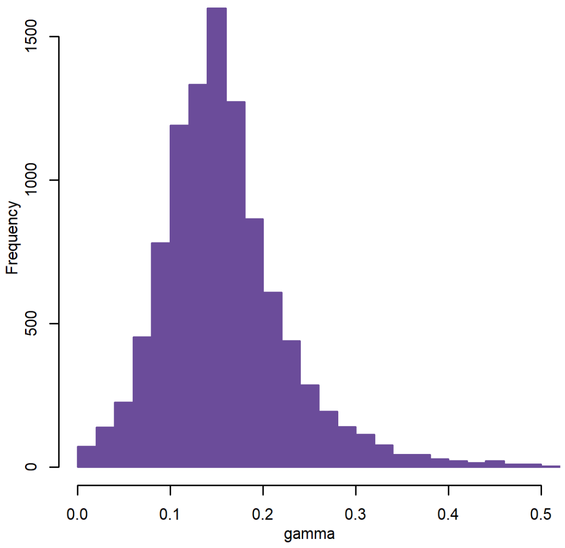

### Cohort 25

Author: Furth

Publication year: 1930

Patient recruitment years: 1912-1913

Cohort size: 8

Location: Switzerland

Type of TB (smear-status): negative

Details: Pulmonary TB patients
from Barmelweid sanatorium.

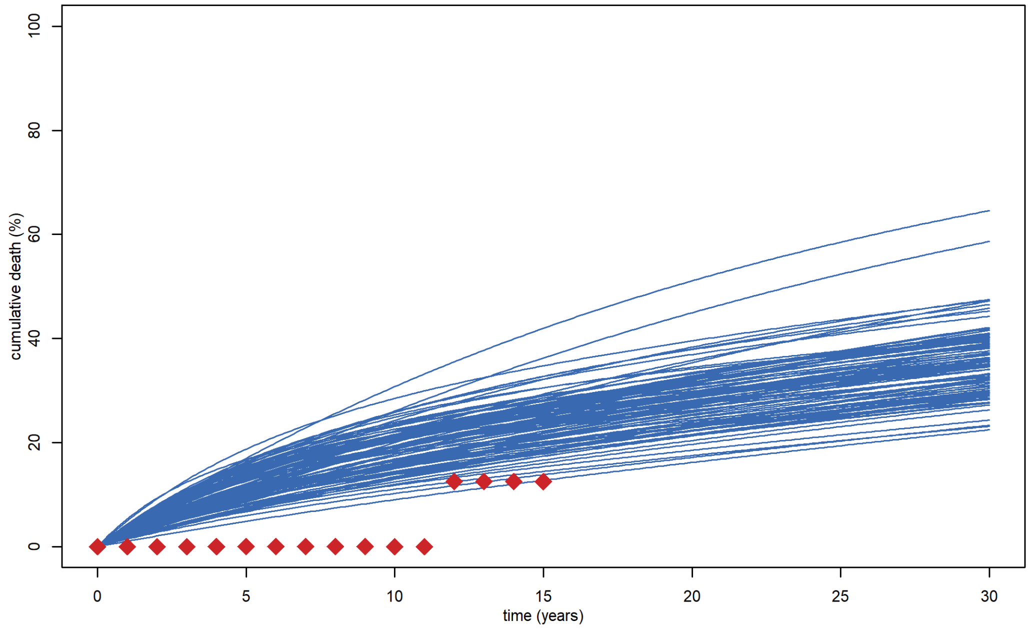


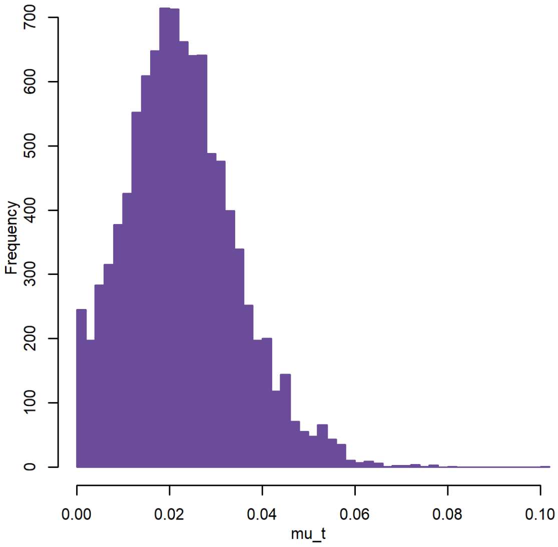


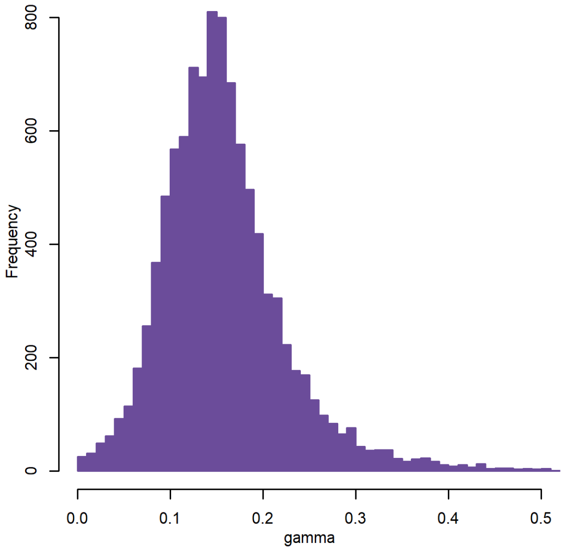

### Cohort 26

Author: Furth

Publication year: 1930

Patient recruitment years: 1913-1914

Cohort size: 36

Location: Switzerland

Type of TB (smear-status): negative

Details: Pulmonary TB patients
from Barmelweid sanatorium.

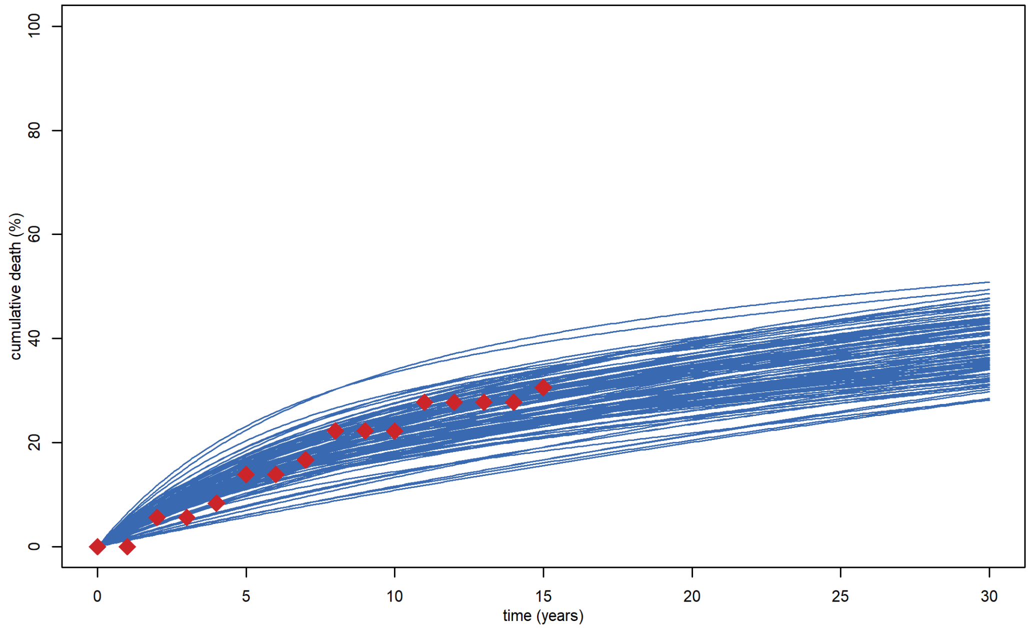


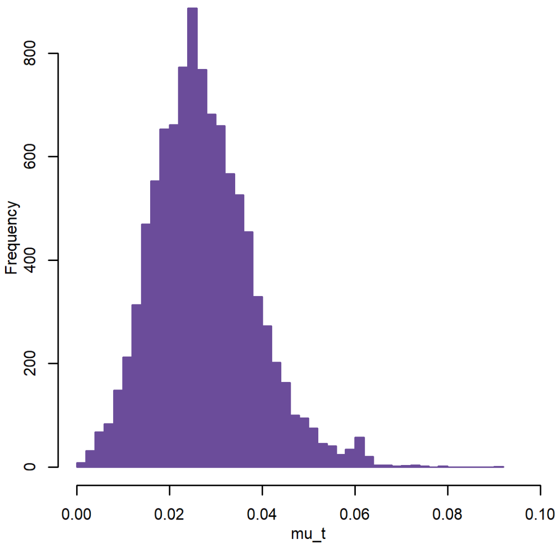


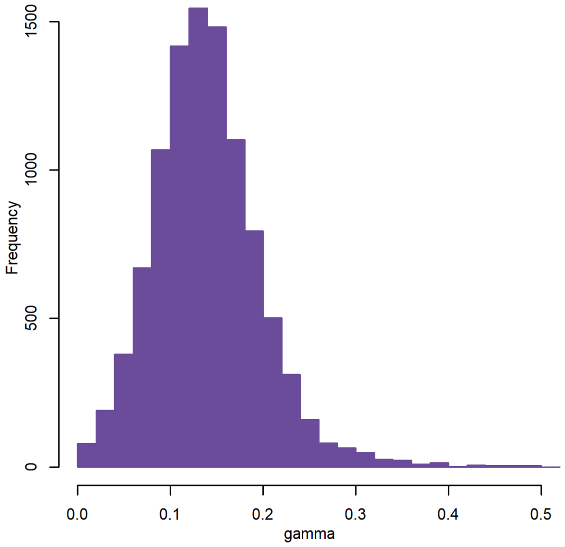

### Cohort 27

Author: Furth

Publication year: 1930

Patient recruitment years: 1914-1915

Cohort size: 39

Location: Switzerland

Type of TB (smear-status): negative

Details: Pulmonary TB patients from Barmelweid sanatorium.

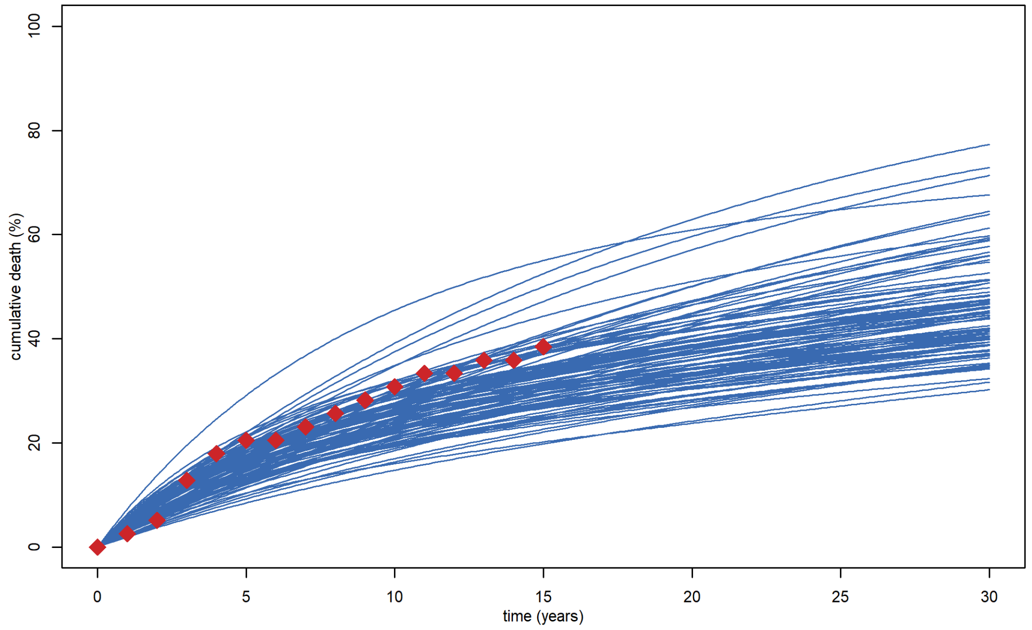


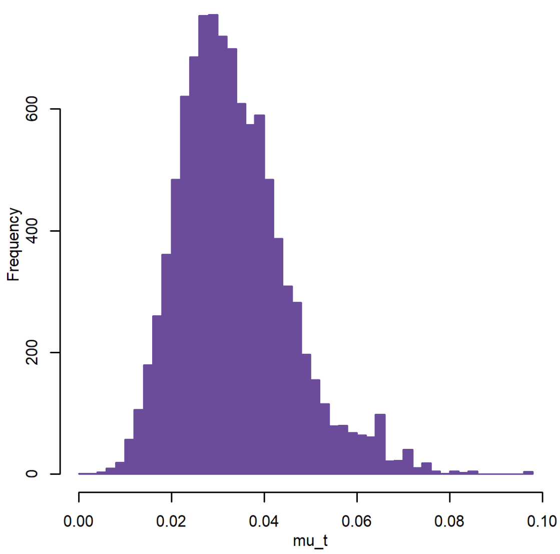


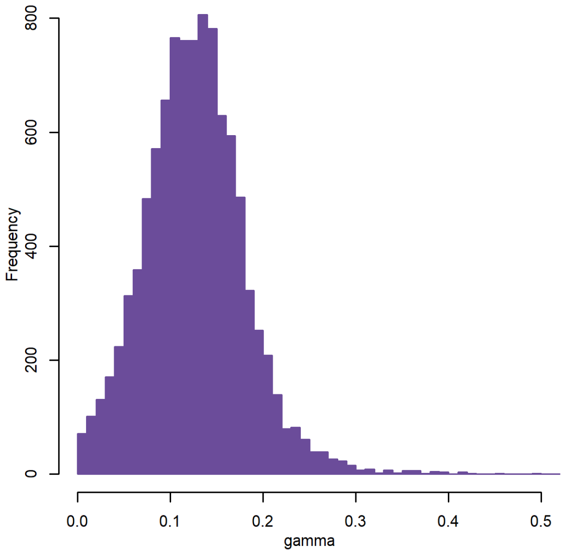

### Cohort 28

Author: Furth

Publication year: 1930

Patient recruitment years: 1915-1916

Cohort size: 39

Location: Switzerland

Type of TB (smear-status): negative

Details: Pulmonary TB patients
from Barmelweid sanatorium.

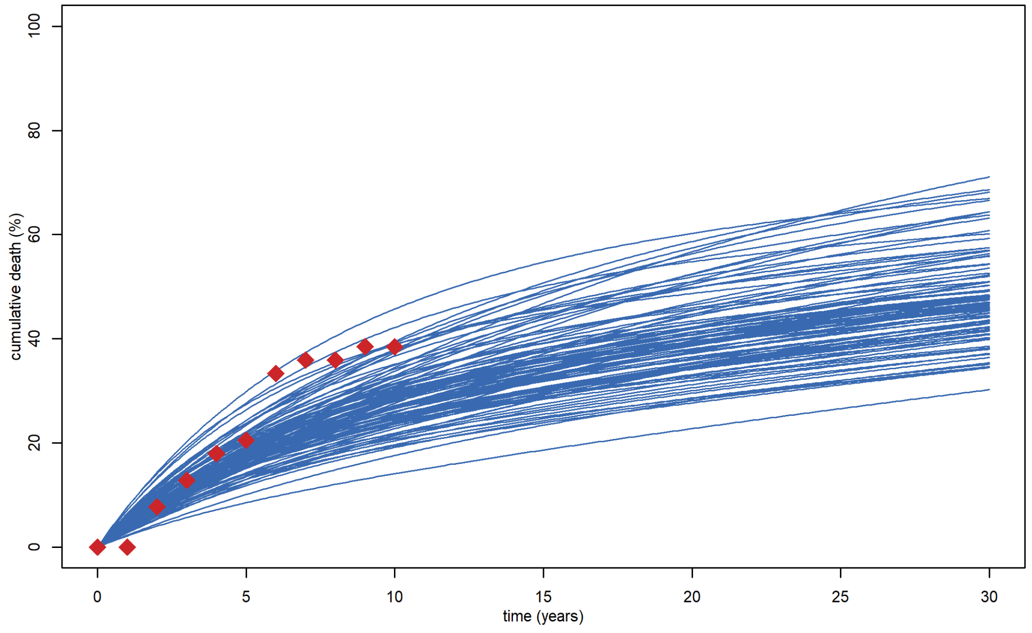


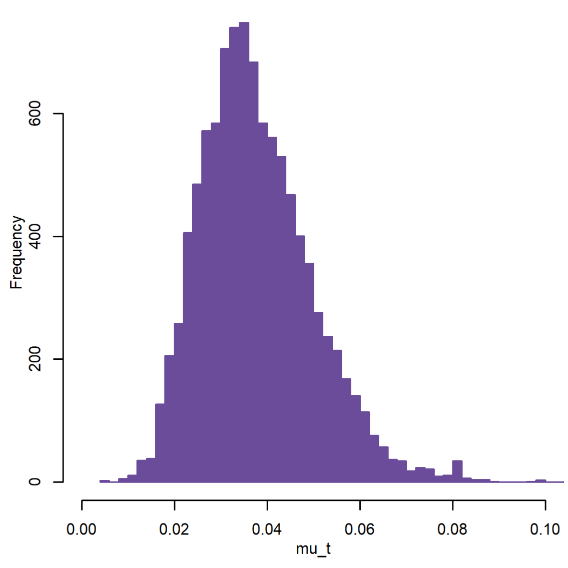


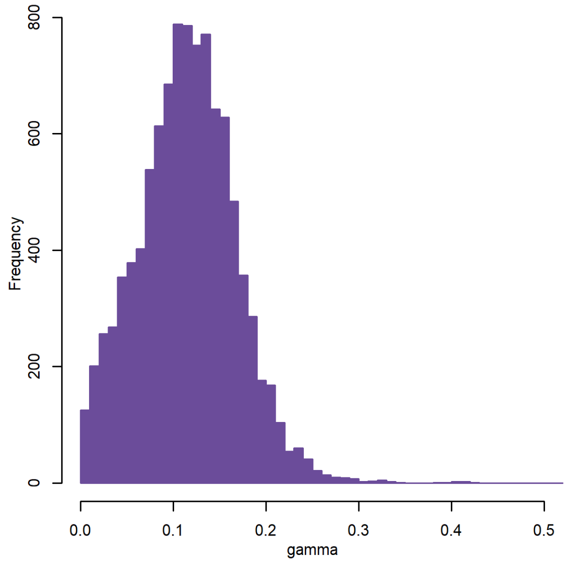

### Cohort 29

Author: Furth

Publication year: 1930

Patient recruitment years: 1916-1917

Cohort size: 28

Location: Switzerland

Type of TB (smear-status): negative

Details: Pulmonary TB patients
from Barmelweid sanatorium.

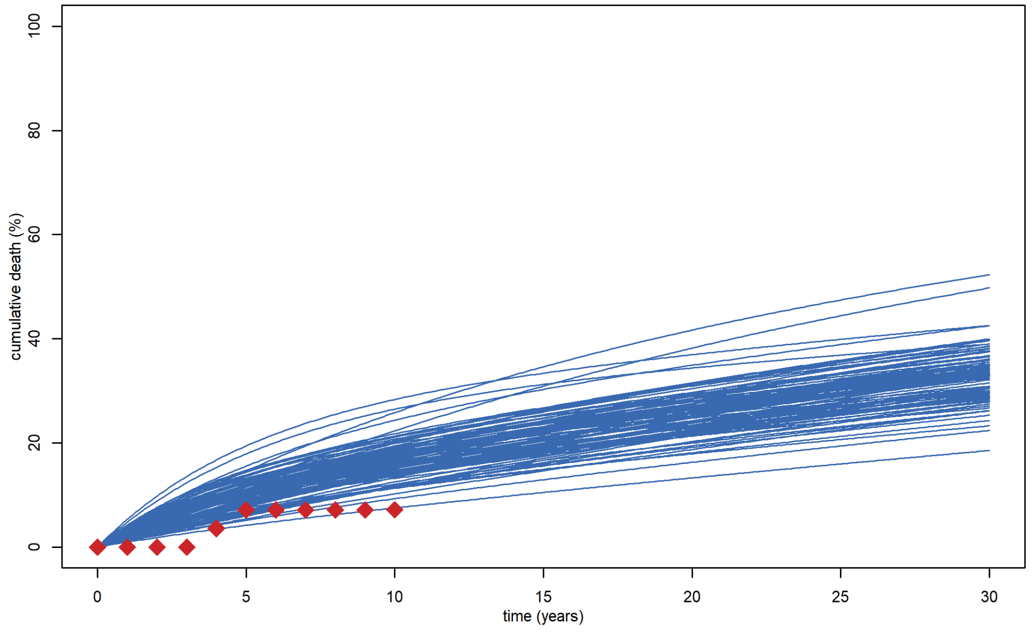


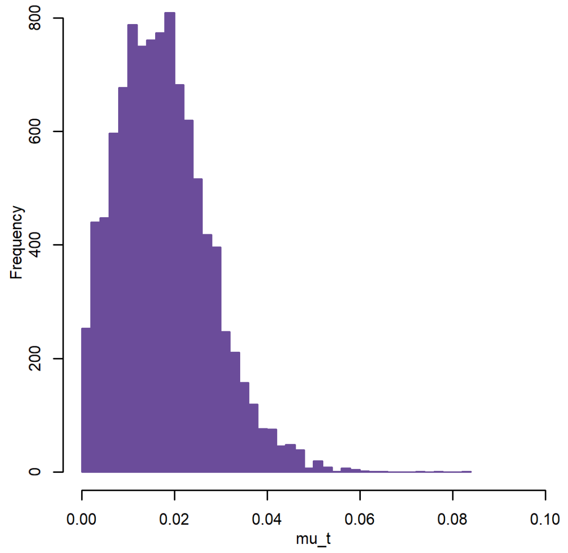


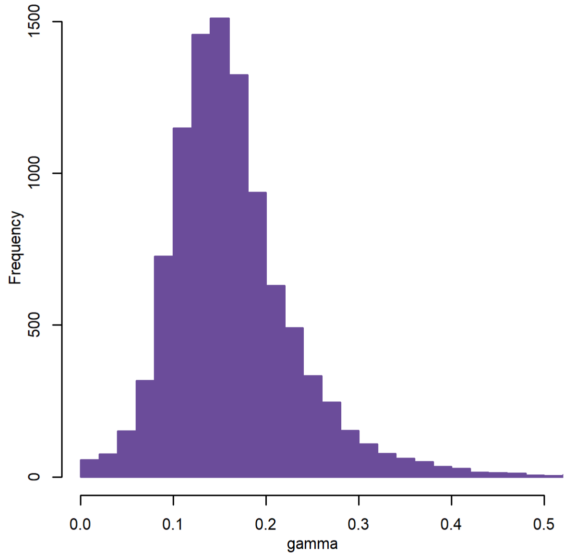

### Cohort 30

Author: Furth

Publication year: 1930

Patient recruitment years: 1917-1918

Cohort size: 38

Location: Switzerland

Type of TB (smear-status): negative

Details: Pulmonary TB patients from Barmelweid sanatorium.

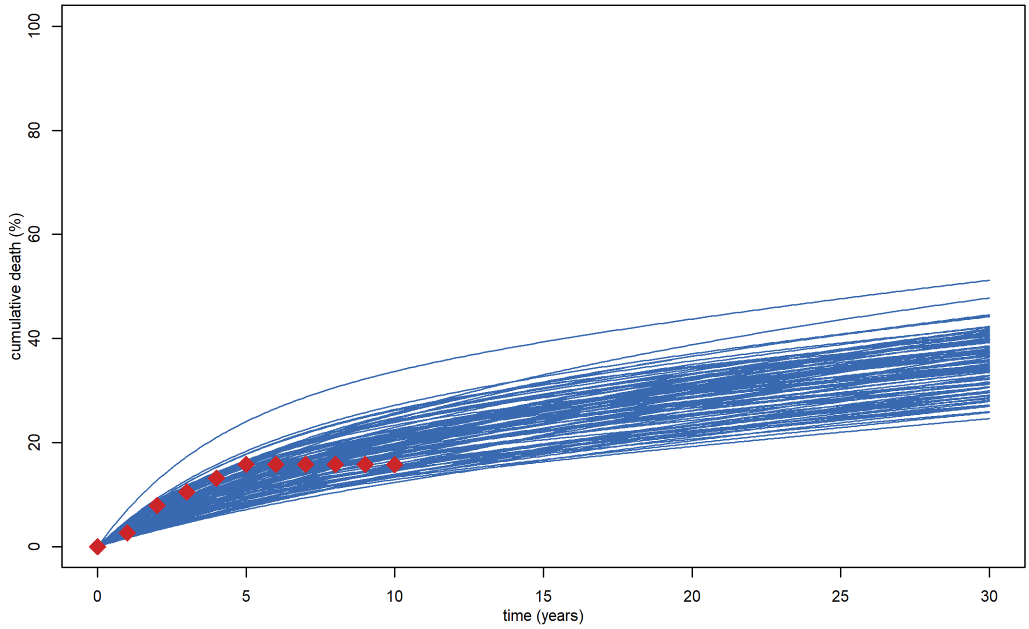


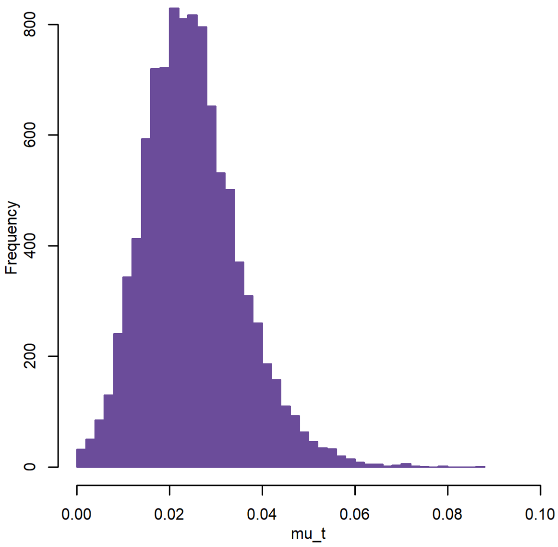


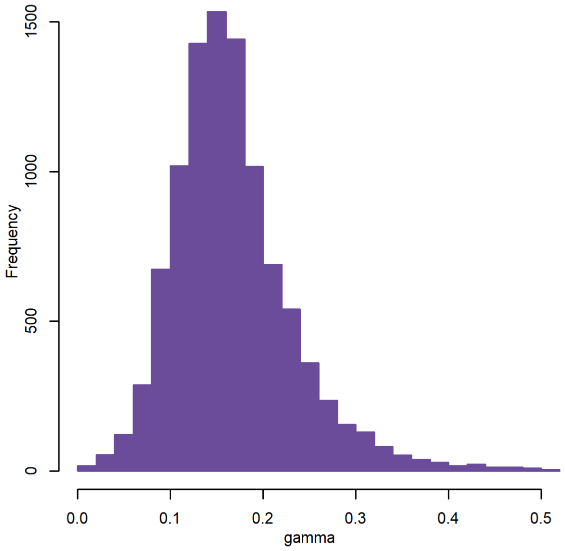

### Cohort 31

Author: Furth

Publication year: 1930

Patient recruitment years: 1918-1919

Cohort size: 41

Location: Switzerland

Type of TB (smear-status): negative

Details: Pulmonary TB patients from Barmelweid sanatorium.

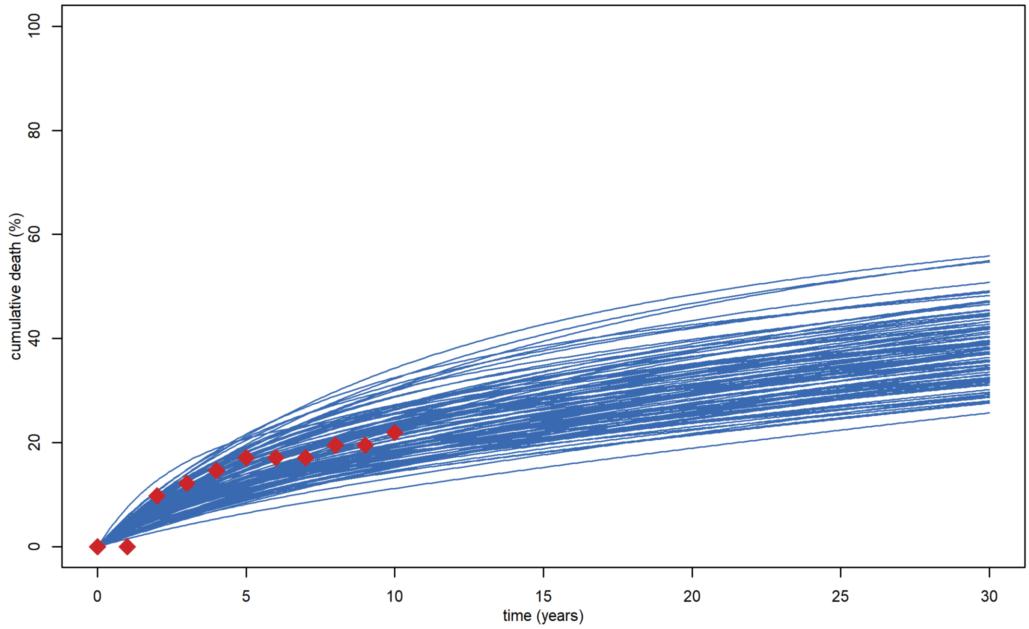


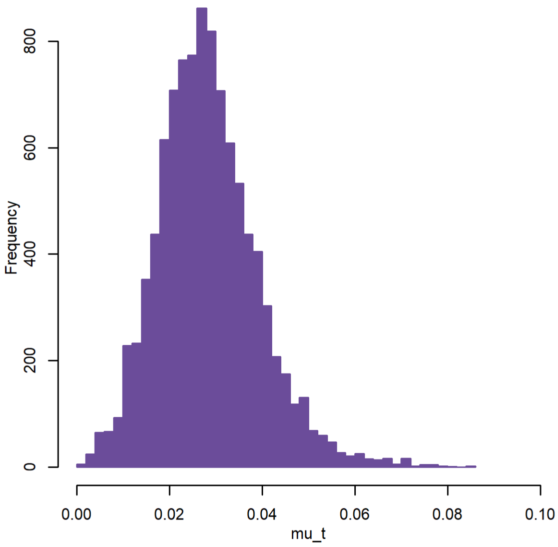


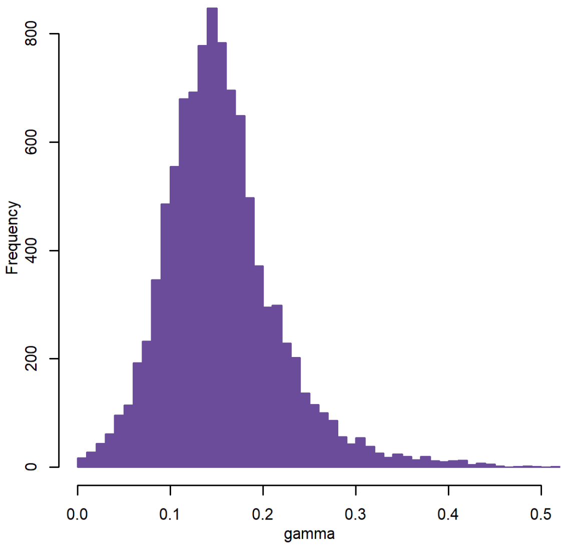

### Cohort 32

Author: Furth
Publication year: 1930
Patient recruitment years: 1919-1920
Cohort size: 37

Location: Switzerland
Type of TB (smear-status): negative
Details: Pulmonary TB patients
from Barmelweid sanatorium.

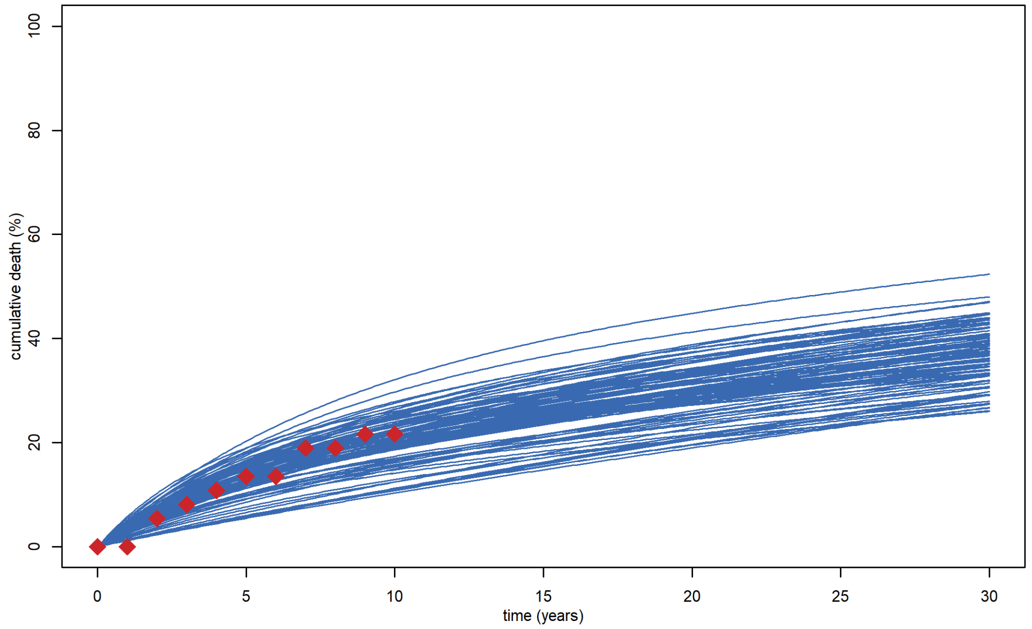


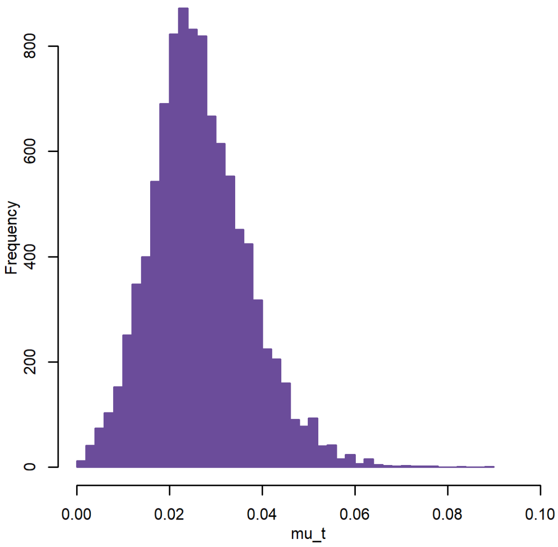


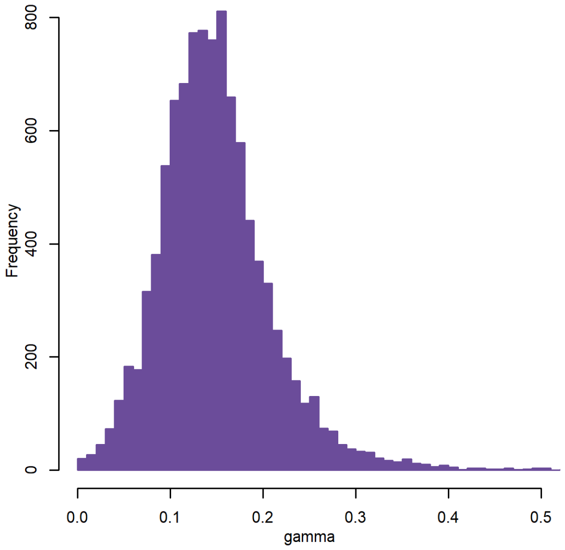

### Cohort 33

Author: Furth

Publication year: 1930

Patient recruitment years: 1920-1921

Cohort size: 55

Location: Switzerland

Type of TB (smear-status): negative

Details: Pulmonary TB patients from Barmelweid sanatorium.

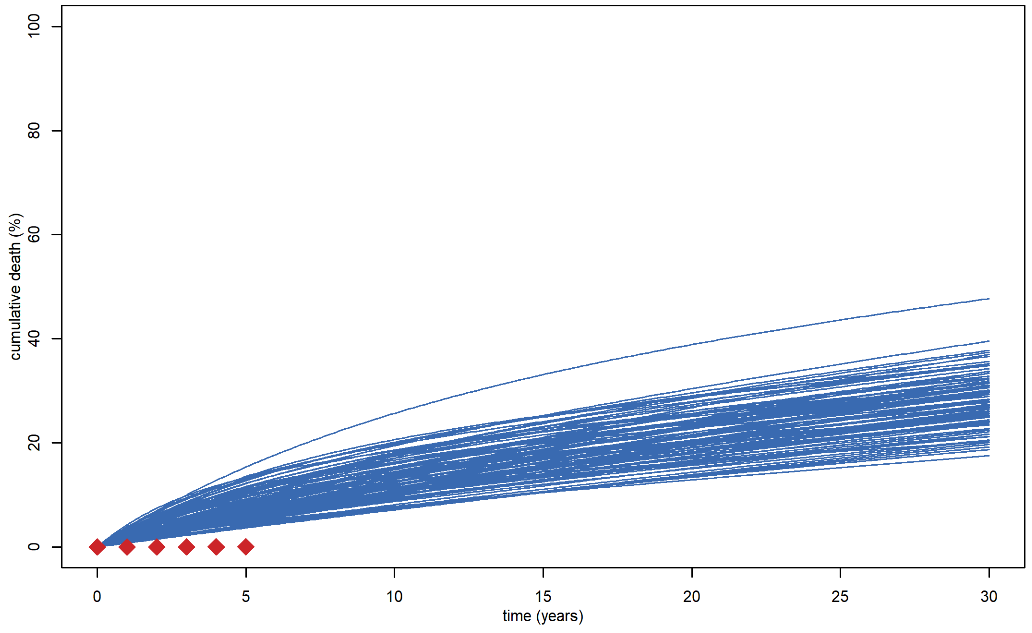


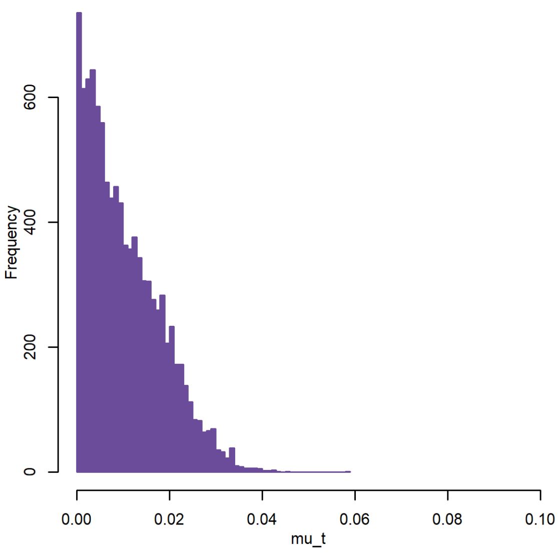


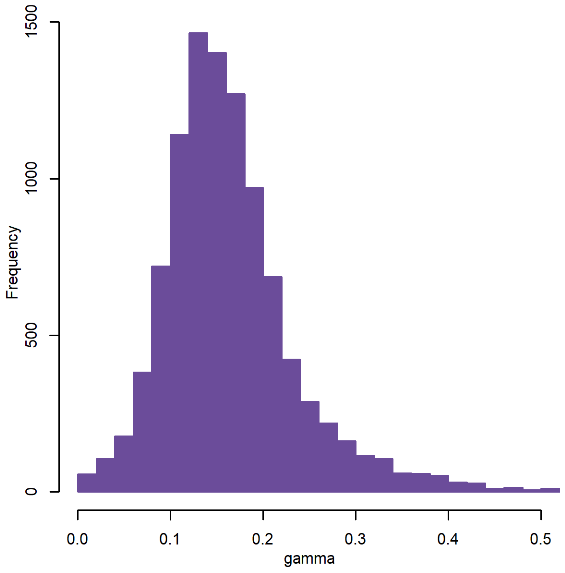

### Cohort 34

Author: Furth

Publication year: 1930

Patient recruitment years: 1921-1922

Cohort size: 36

Location: Switzerland

Type of TB (smear-status): negative

Details: Pulmonary TB patients
from Barmelweid sanatorium.

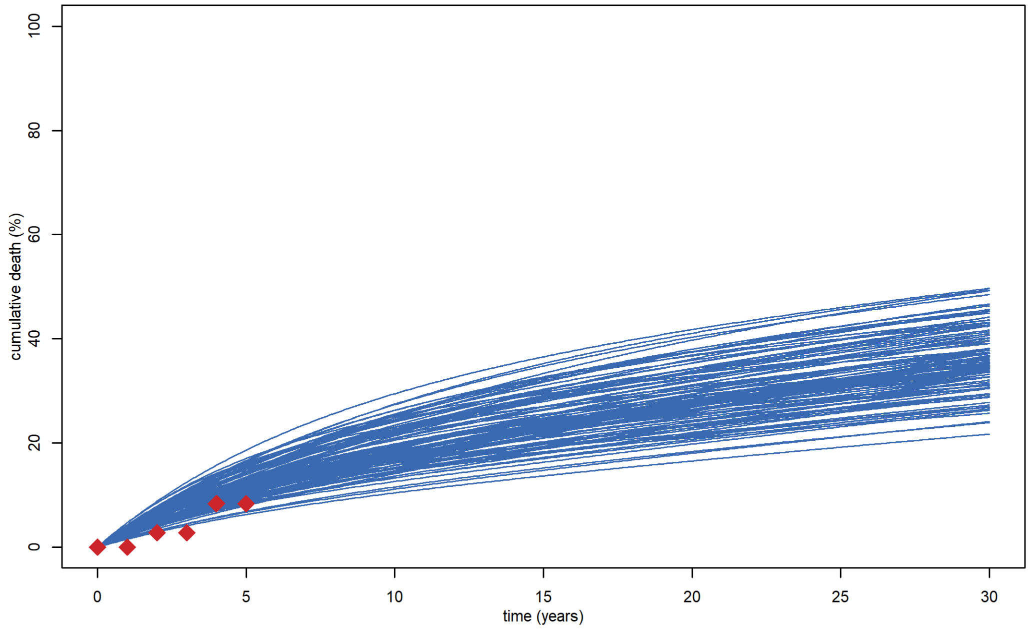


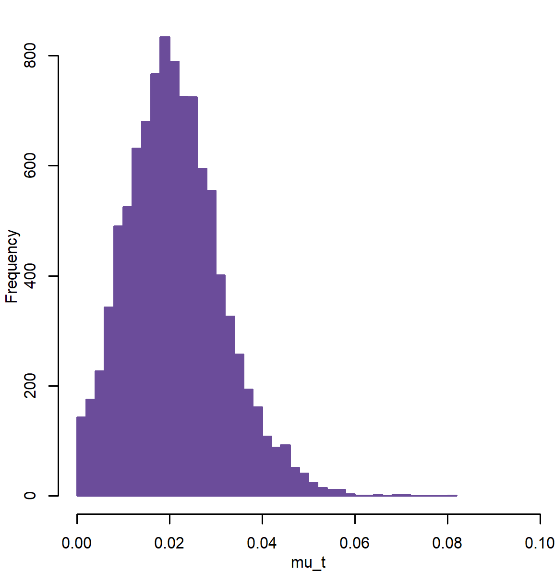


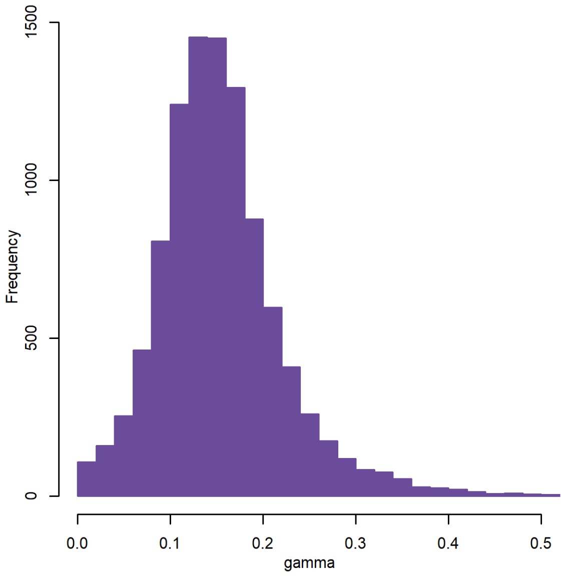

### Cohort 35

Author: Furth

Publication year: 1930

Patient recruitment years: 1922-1923

Cohort size: 45

Location: Switzerland

Type of TB (smear-status): negative

Details: Pulmonary TB patients
from Barmelweid sanatorium.

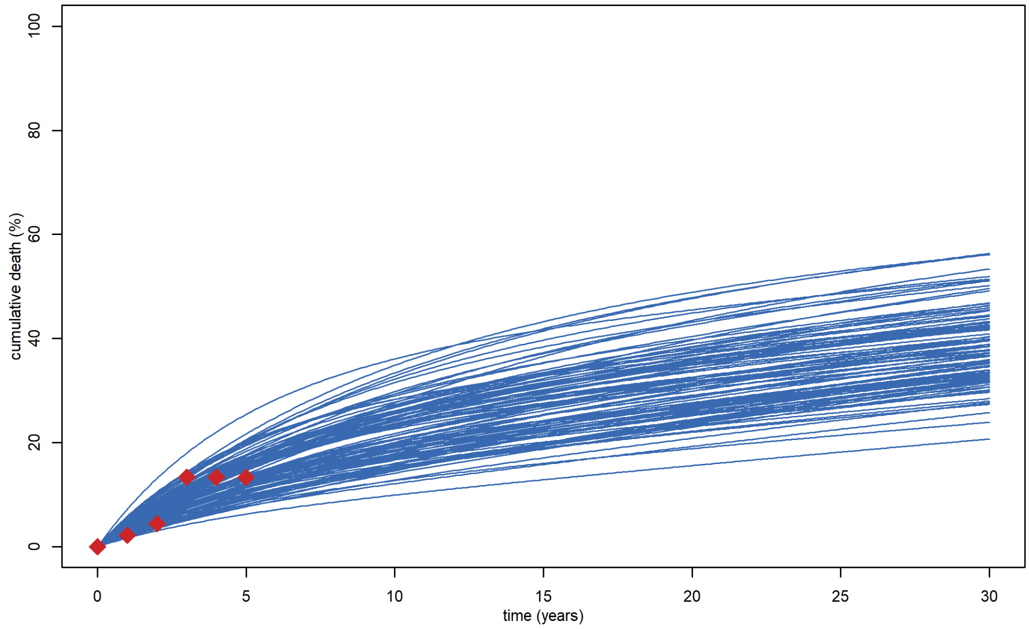


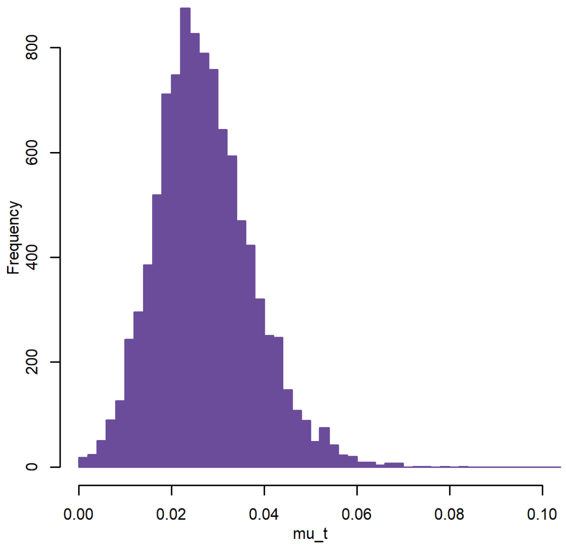


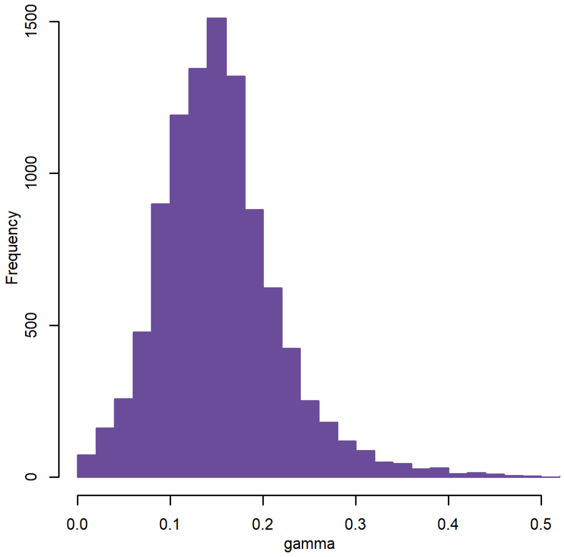

### Cohort 36

Author: Furth

Publication year: 1930

Patient recruitment years: 1923-1924

Cohort size: 33

Location: Switzerland

Type of TB (smear-status): negative

Details: Pulmonary TB patients from Barmelweid sanatorium.

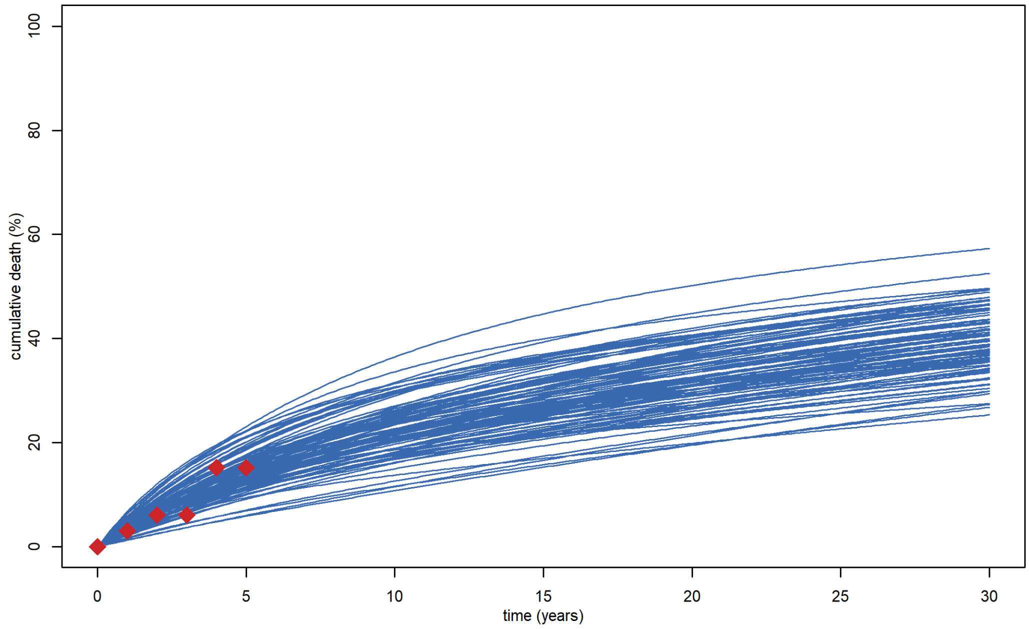


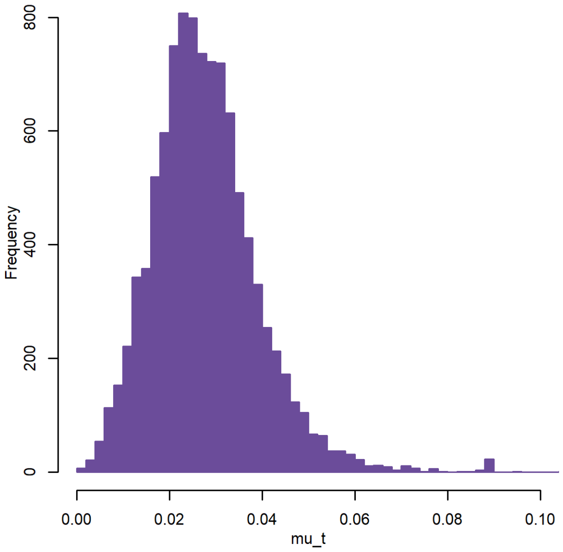


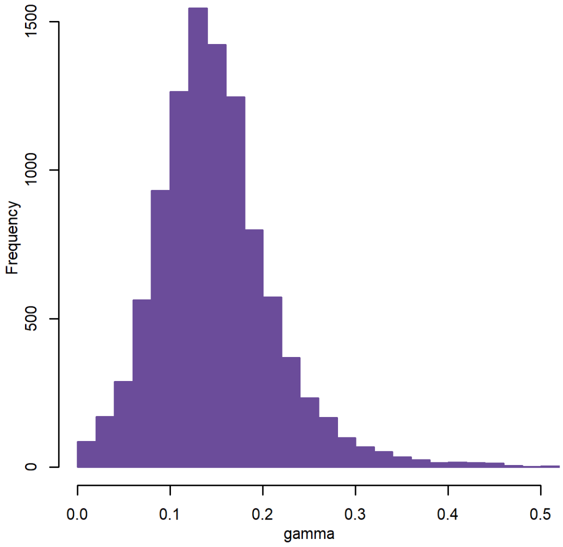

### Cohort 37

Author: Furth

Publication year: 1930

Patient recruitment years: 1924-1925

Cohort size: 34

Location: Switzerland

Type of TB (smear-status): negative

Details: Pulmonary TB patients
from Barmelweid sanatorium.

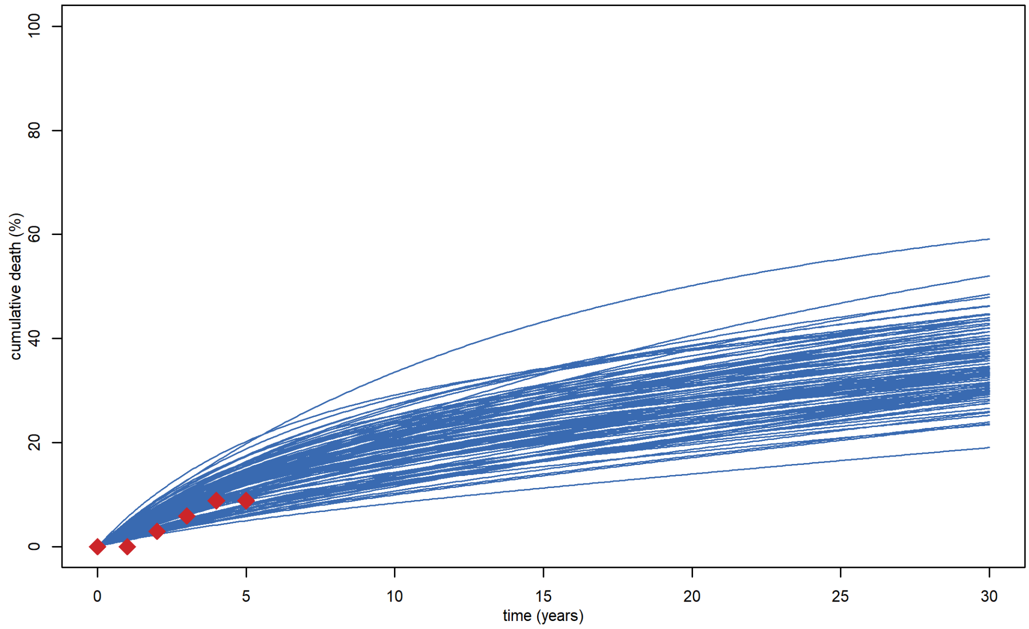


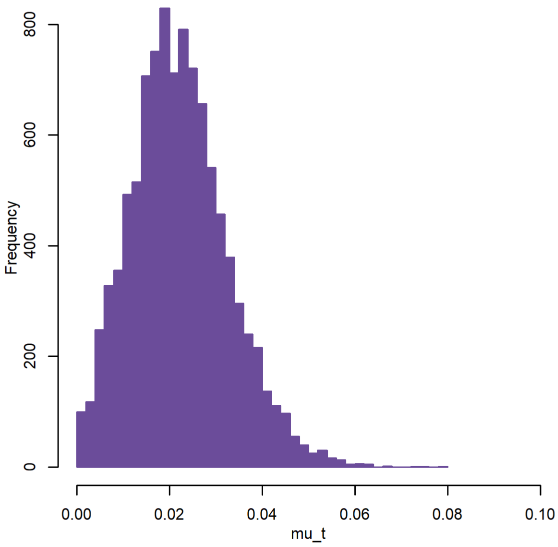


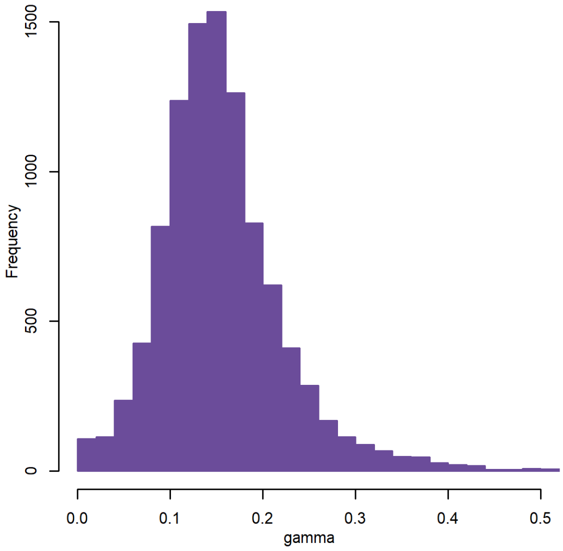

### Cohort 57

Author: Magnusson

Publication year: 1938

Patient recruitment years: 1916-1935

Cohort size: 133

Location: Iceland

Type of TB (smear-status): negative

Details: Cases admitted for treatment at the Vifillsstadir Sanatorium in Reykjavik. Male patients only.

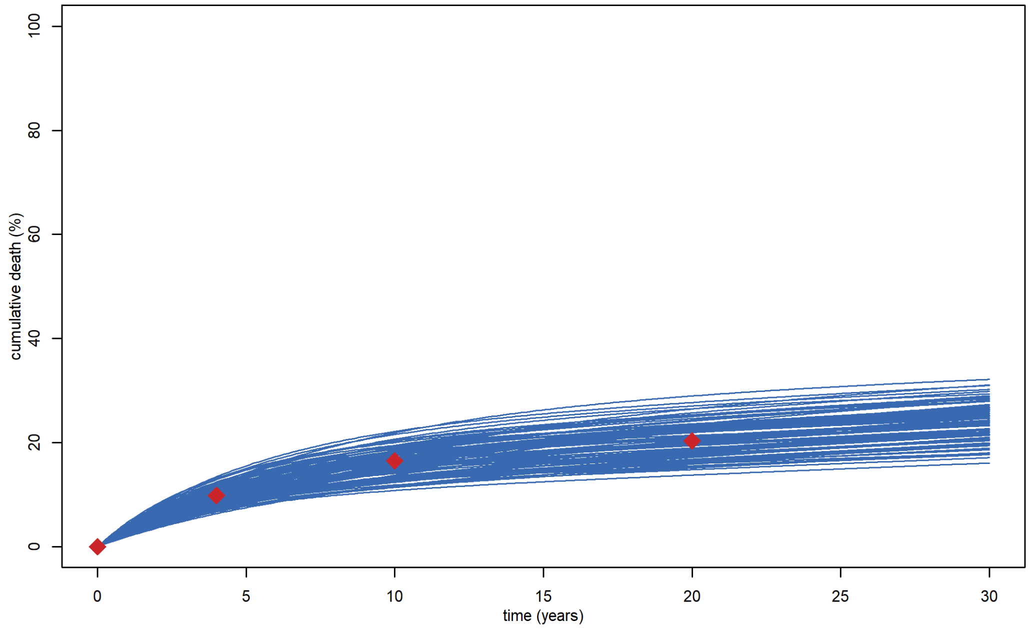


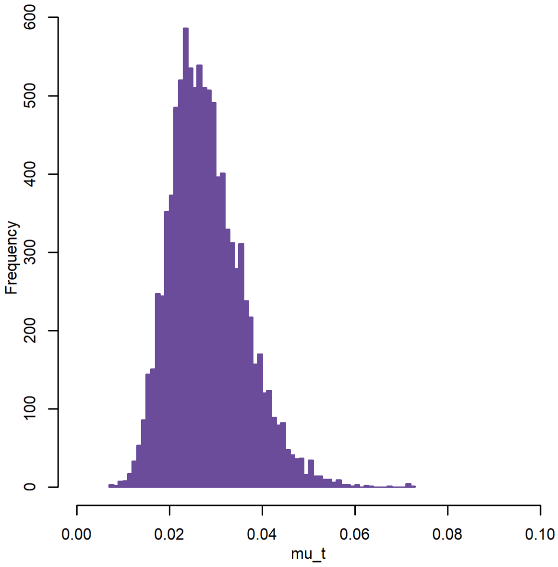


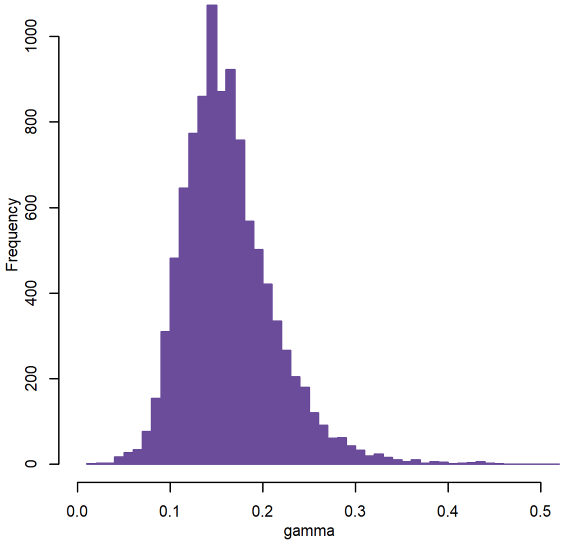

### Cohort 58

Author: Magnusson

Publication year: 1938

Patient recruitment years: 1916-1935

Cohort size: 280

Location: Iceland

Type of TB (smear-status): negative

Details: Cases admitted for treatment at the Vifillsstadir Sanatorium in Reykjavik. Female patients only.

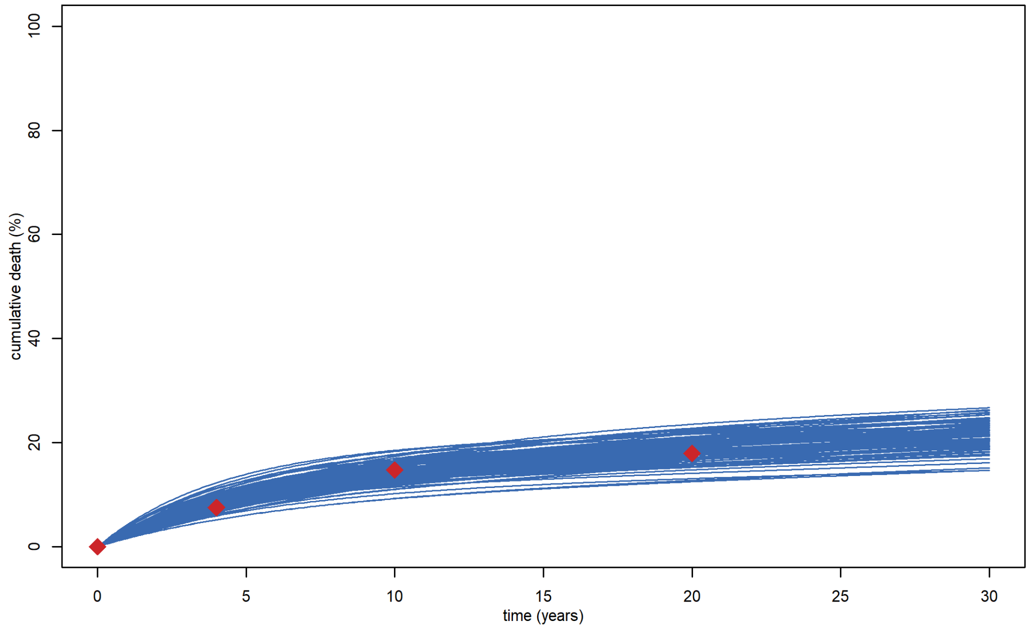


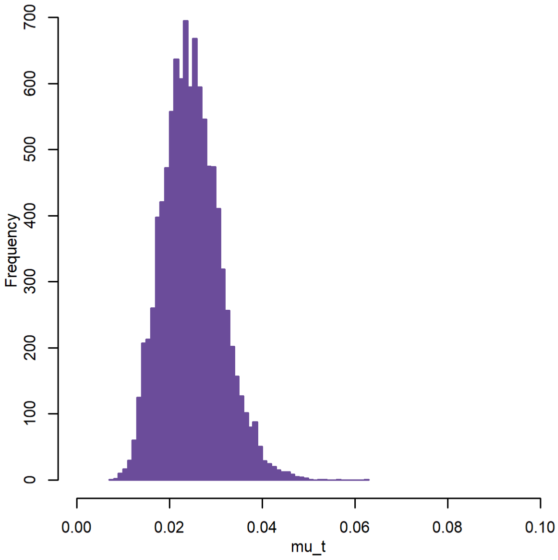


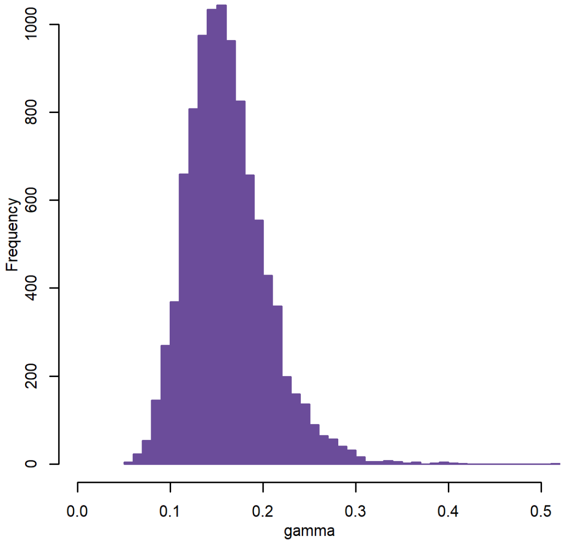
