## Supplementary material for "Revisiting the Natural History of Pulmonary Tuberculosis: a Bayesian Estimation of Natural Recovery and Mortality rates": Cohort profiles smear-positive

#### Cohort profiles of smear-positive TB patients

The cohort profiles below present the main characteristics of the cohorts used in our analysis (top panel), the data points with posterior predictive curves (central panel) and the posterior distributions of the cohort-specific parameters $\gamma$ and $\mu_T$ (bottom panels). In the central panel, the data are represented with red diamonds and we show model realisations associated with 100 randomly selected posterior samples obtained from the MCMC simulation (blue lines).

### Cohort 1

Author: Baart De La Faille

Publication year: 1939

Patient recruitment years: 1922-1925

Cohort size: 177

Location: The Netherlands

Type of TB (smear-status): positive

Details: TB cases hospitalized in the Sanatorium "Berg en Bosch".

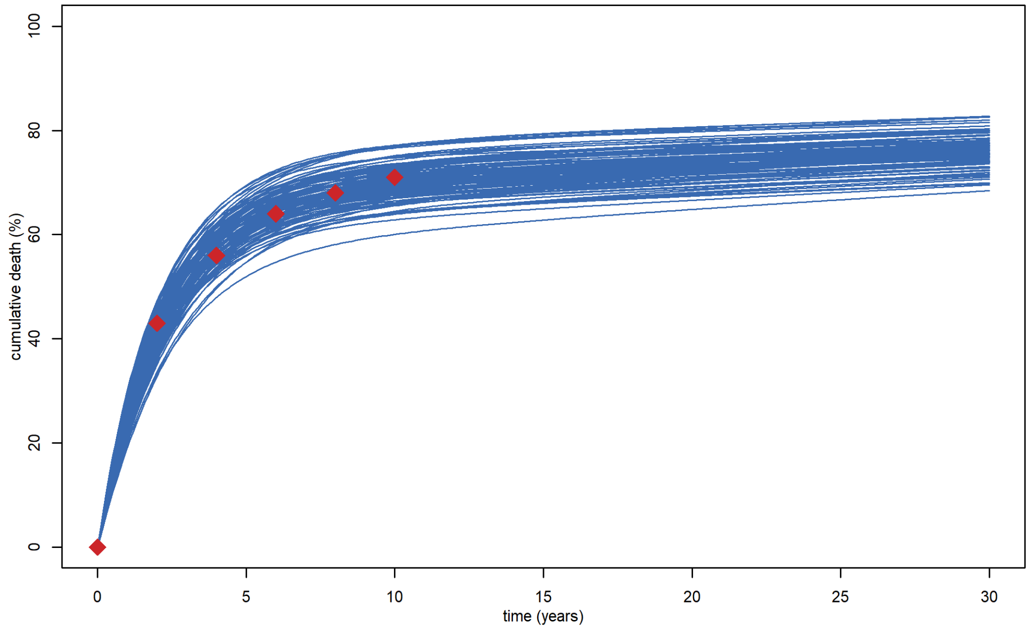


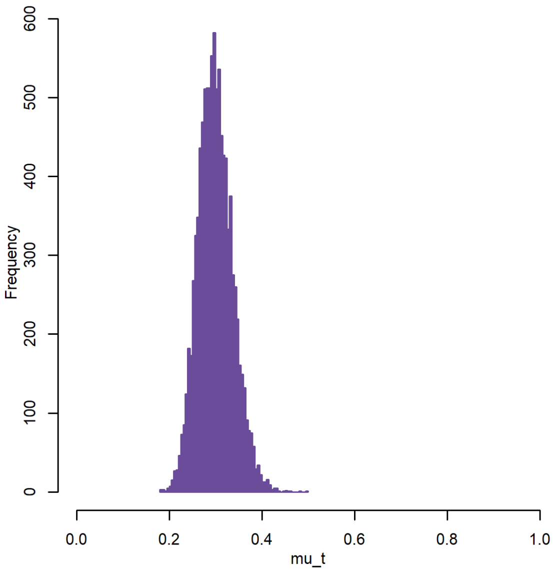


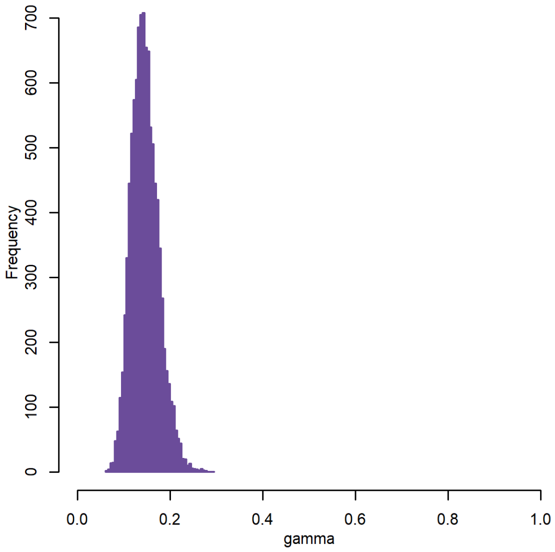

### Cohort 2

Author: Baart De La Faille

Publication year: 1939

Patient recruitment years: 1926-1929

Cohort size: 149

Location: The Netherlands

Type of TB (smear-status): positive

Details: TB cases hospitalized in the Sanatorium "Berg en Bosch".

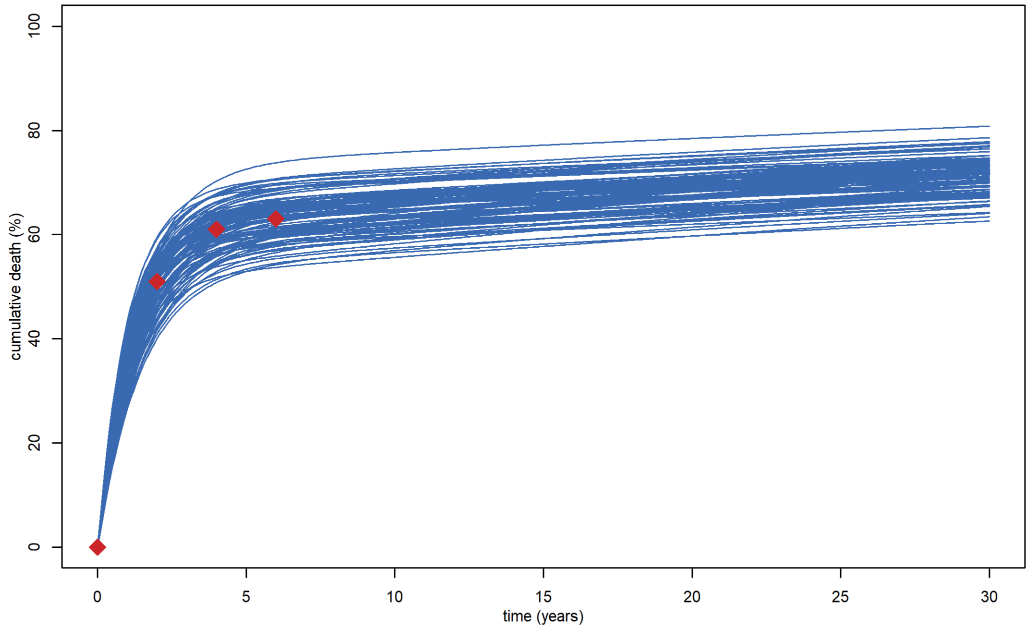


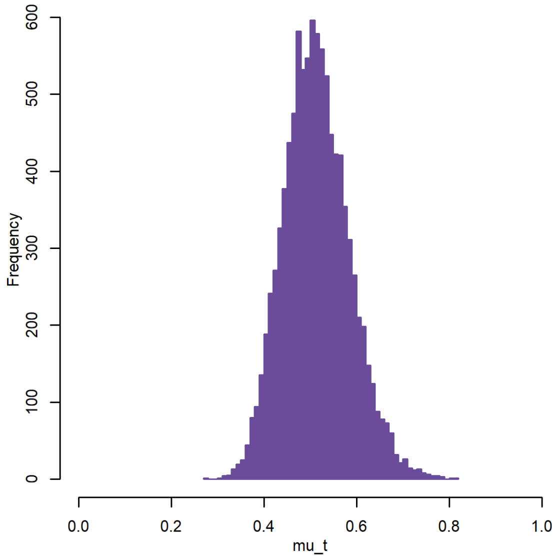


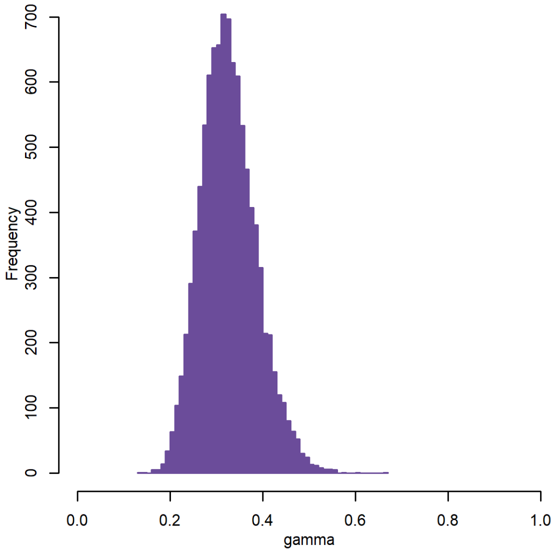

### Cohort 3

Author: Baart De La Faille

Publication year: 1939

Patient recruitment years: 1930-1933

Cohort size: 125

Location: The Netherlands

Type of TB (smear-status): positive

Details: TB cases hospitalized in the Sanatorium "Berg en Bosch".

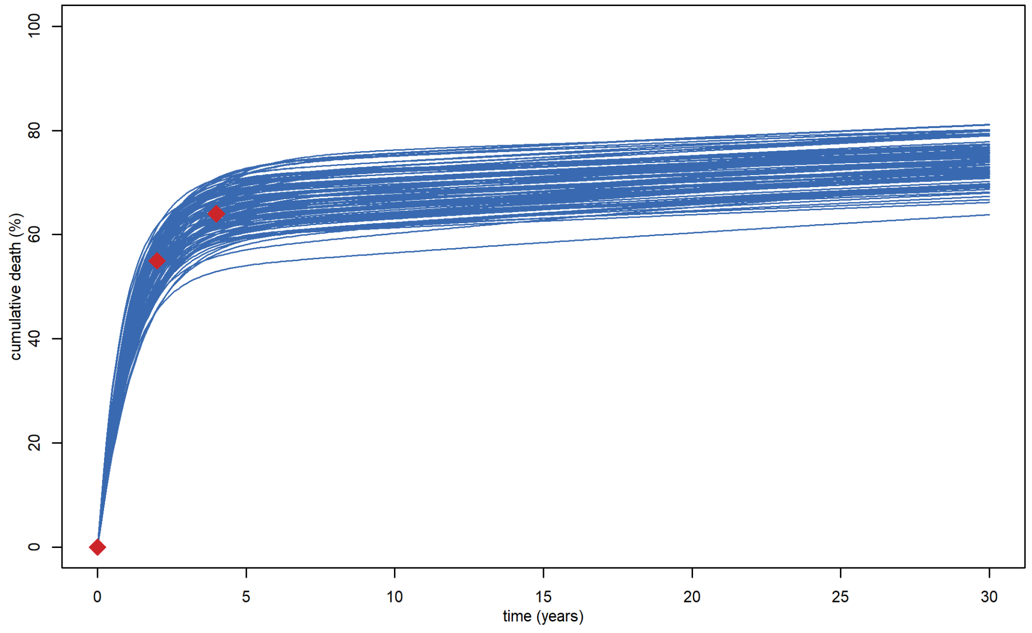


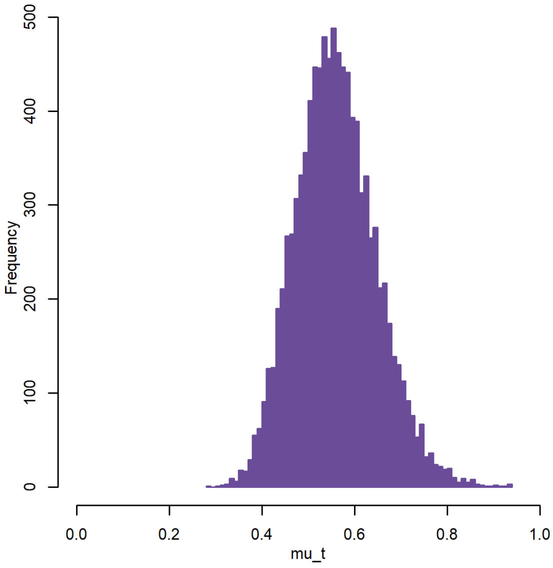


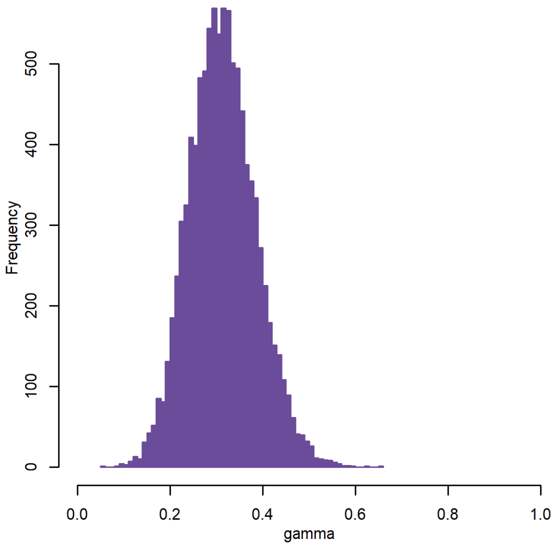

### Cohort 4

Author: Baart De La Faille

Publication year: 1939

Patient recruitment years: 1934-1935

Cohort size: 83

Location: The Netherlands

Type of TB (smear-status): positive

Details: TB cases hospitalized in the Sanatorium "Berg en Bosch".

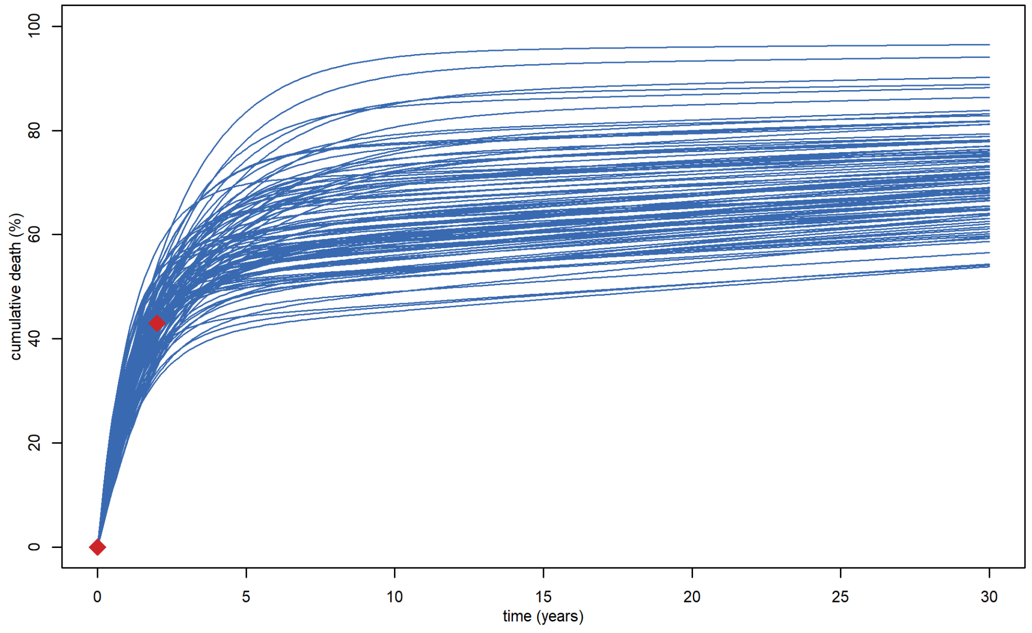


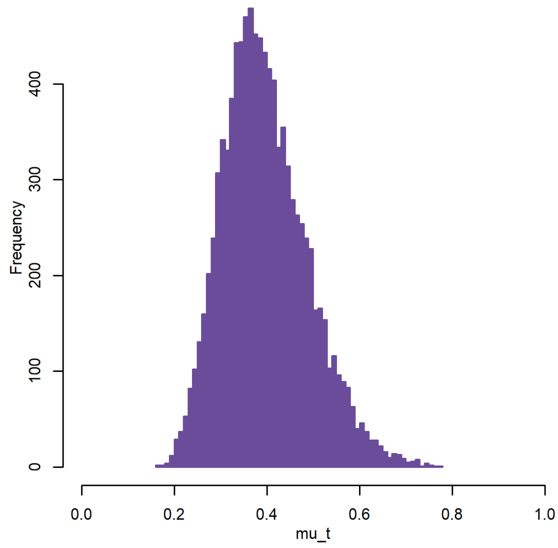


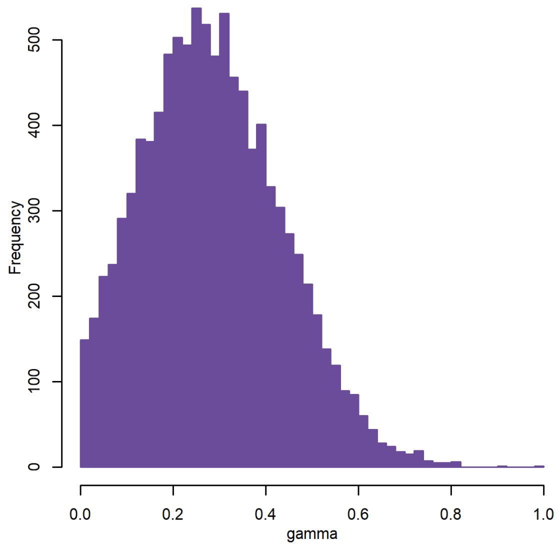

### Cohort 9

Author: Buhl

Publication year: 1967

Patient recruitment years: 1925-1929

Cohort size: 314

Location: Denmark

Type of TB (smear-status): positive

Details: Danish TB patients
diagnosed between 1925 and 1954.

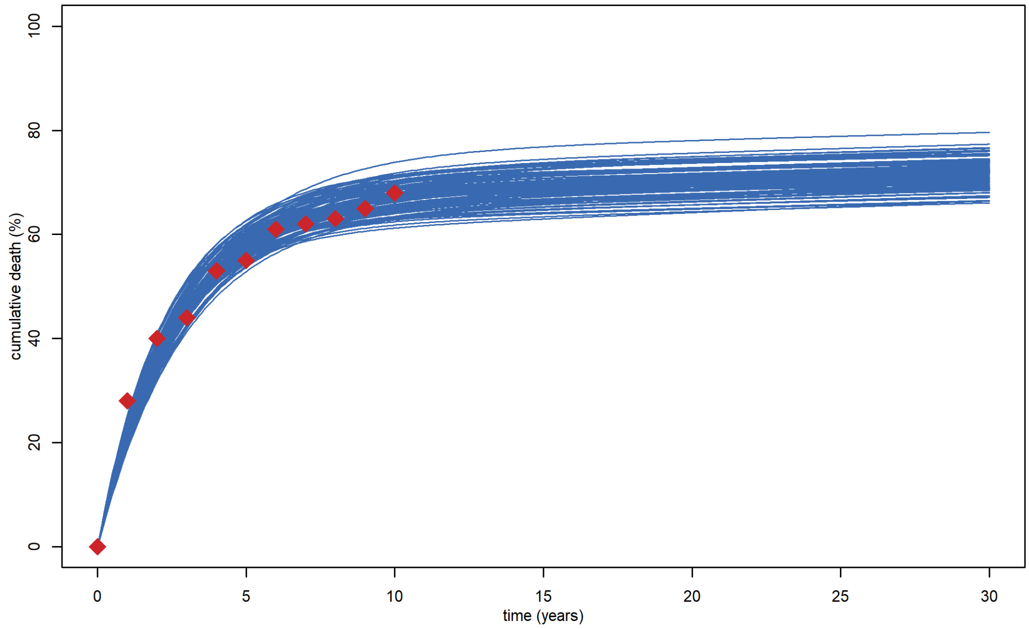


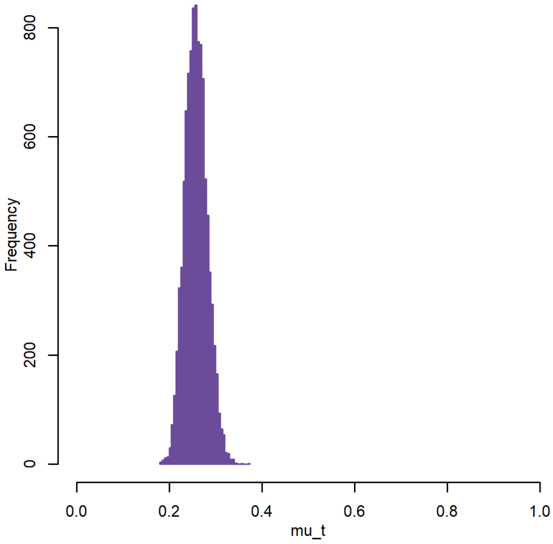


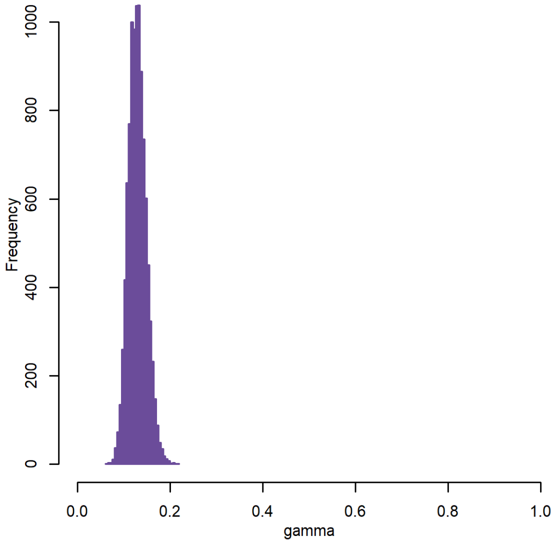

### Cohort 10

Author: Griep

Publication year: 1939

Patient recruitment years: 1920-1925

Cohort size: 372

Location: The Netherlands

Type of TB (smear-status): positive

Details: All notified cases of "open" pulmonary TB occurring in The Hague between 1920 and 1937.

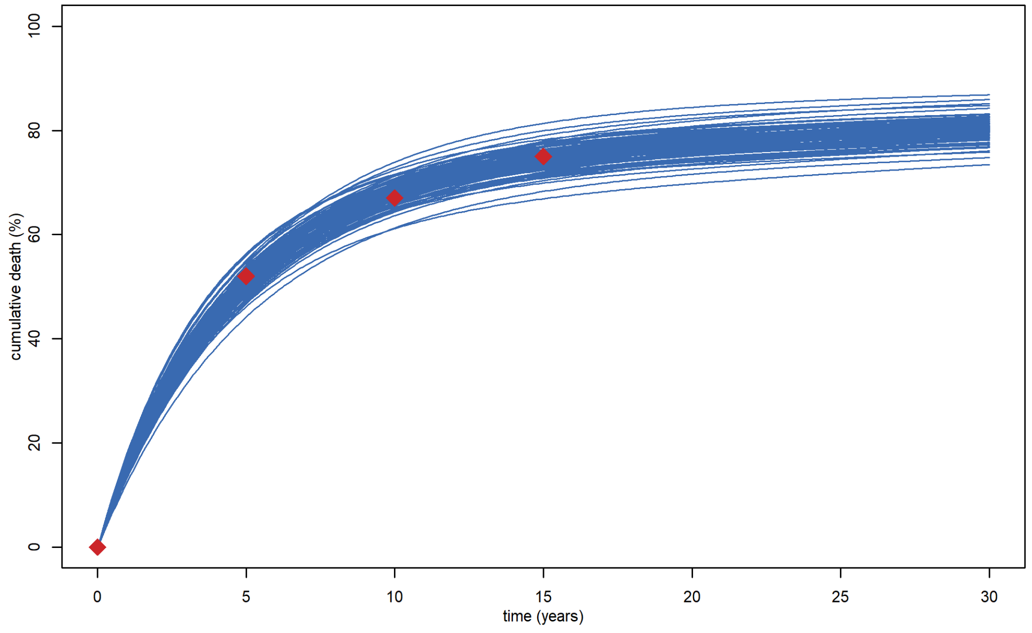


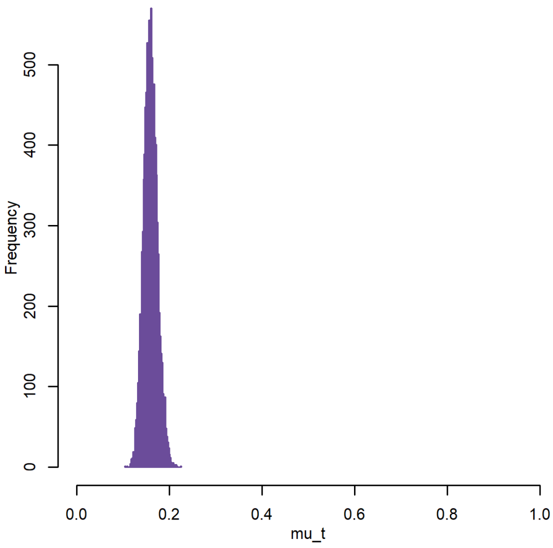


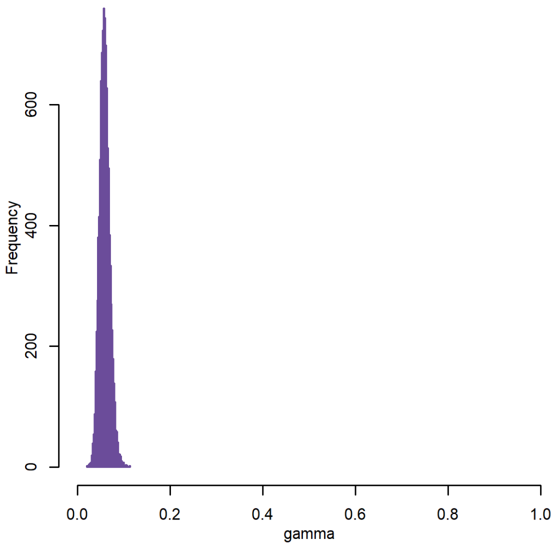

### Cohort 11

Author: Griep

Publication year: 1939

Patient recruitment years: 1925-1930

Cohort size: 603

Location: The Netherlands

Type of TB (smear-status): positive

Details: All notified cases of "open" pulmonary TB occurring in The Hague between 1920 and 1937.

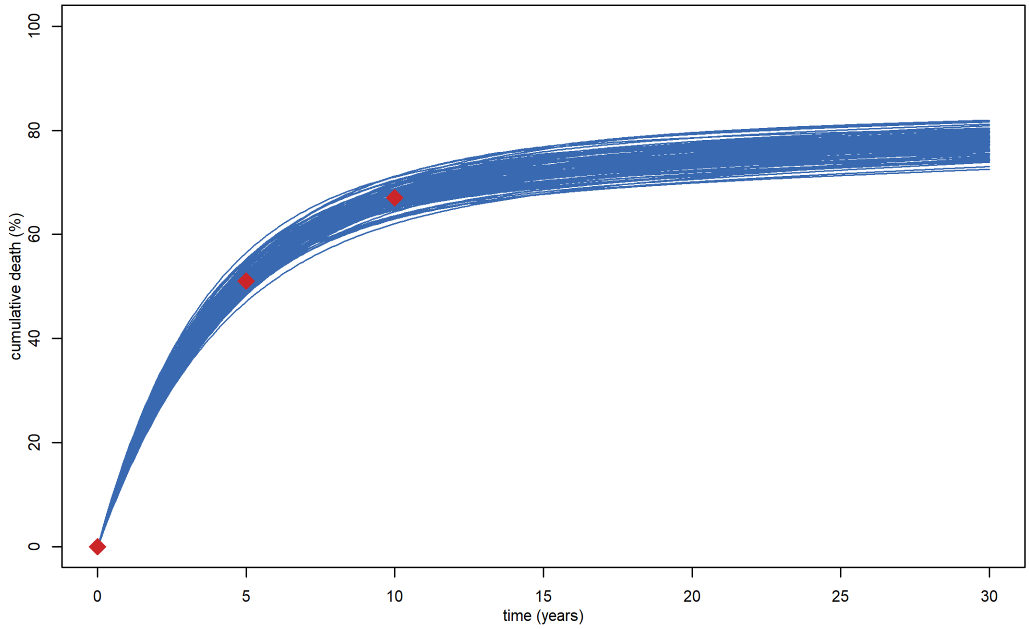


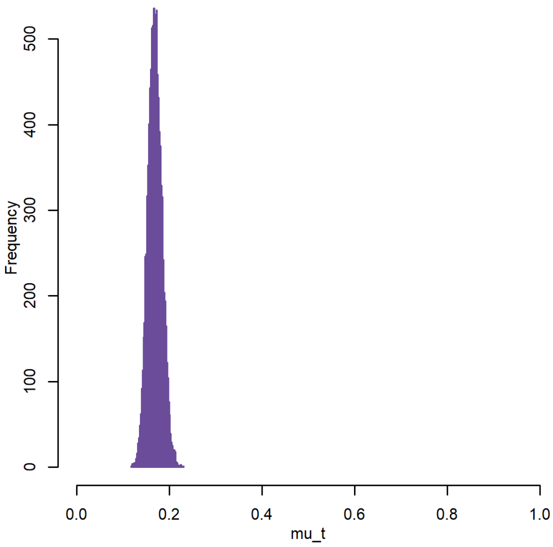


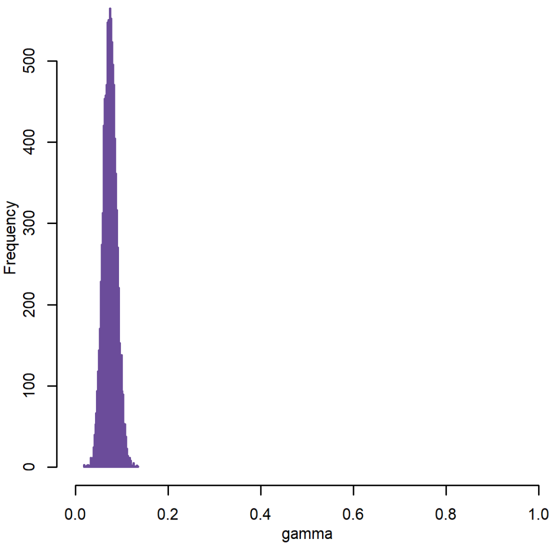

### Cohort 12

Author: Tattersall

Publication year: 1947

Patient recruitment years: 1914-1940

Cohort size: 1192

Location: England

Type of TB (smear-status): positive

Details: Sputum-positive cases attending Reading (UK) dispensary between 1914 and 1940.

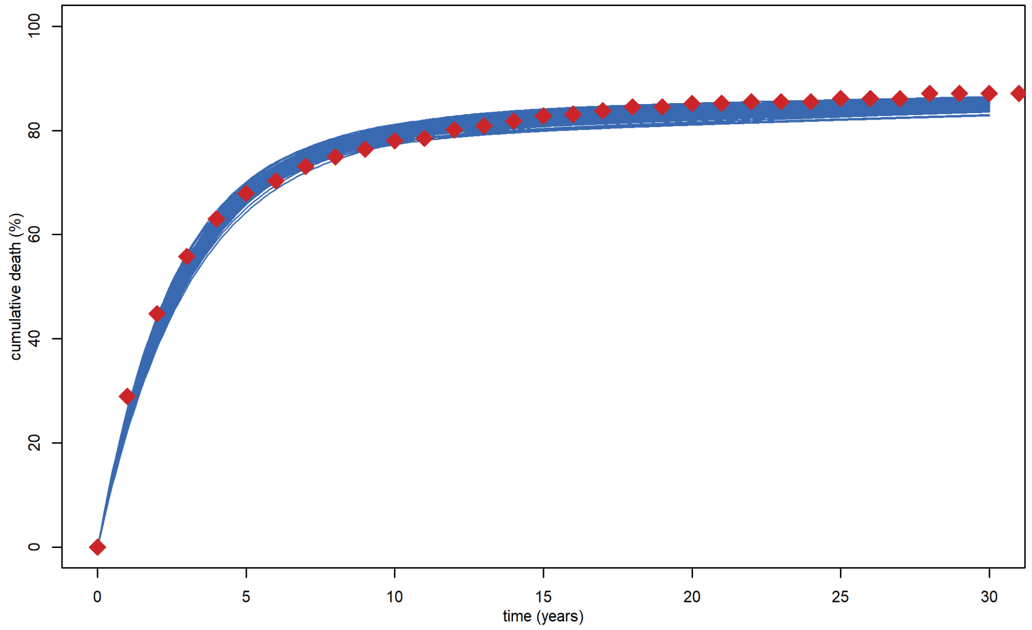


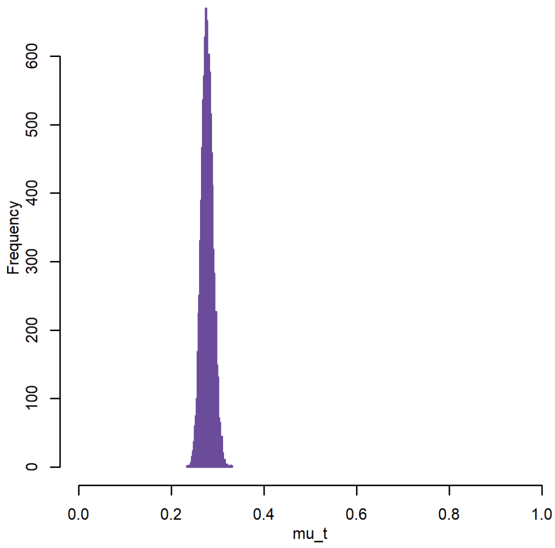


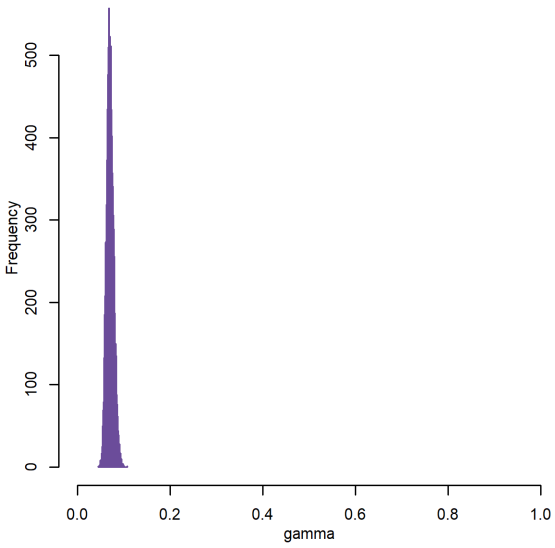

### Cohort 13

Author: Thompson

Publication year: 1943

Patient recruitment years: 1928-1940

Cohort size: 406

Location: England

Type of TB (smear-status): positive

Details: All sputum-positive TB patients occurring in a compact industrial area in Middlesex County.

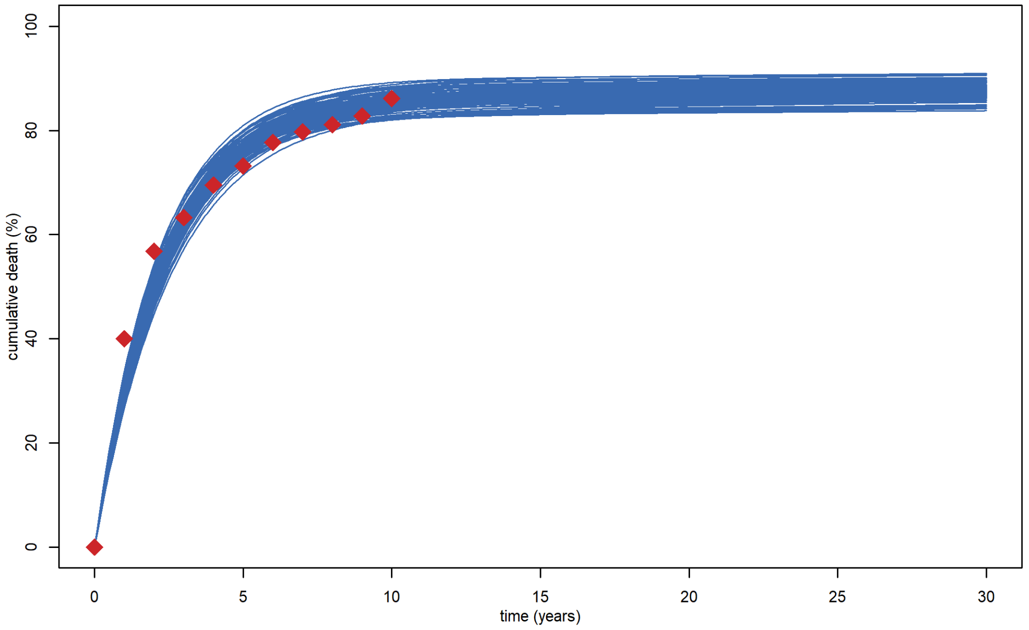


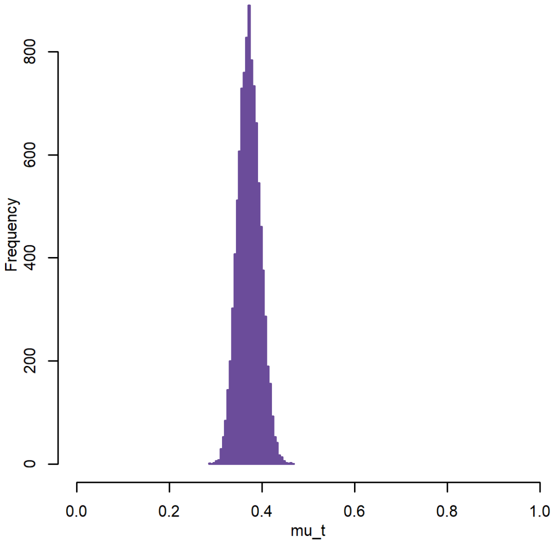


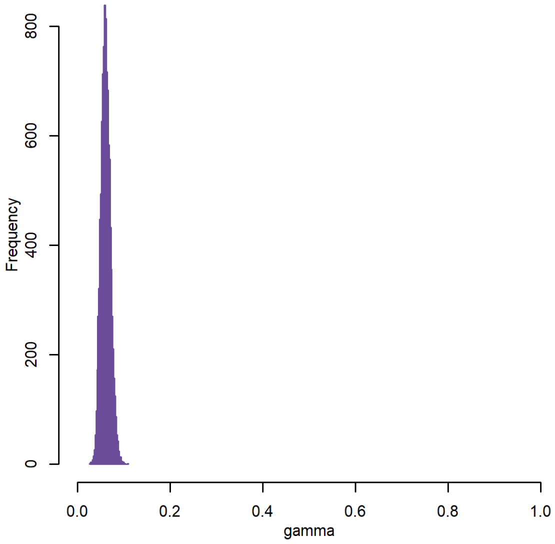

### Cohort 14

Author: Hartley

Publication year: 1935

Patient recruitment years: 1905-1914

Cohort size: 2382

Location: England

Type of TB (smear-status): positive

Details: Retrospective cohort study of cases treated for TB at Brompton Hospital.

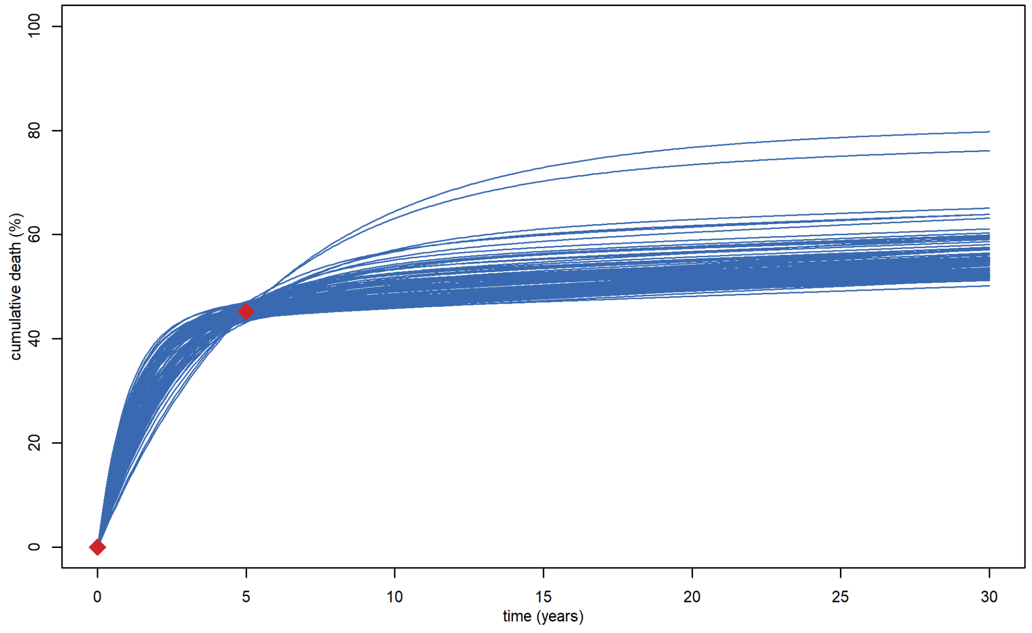


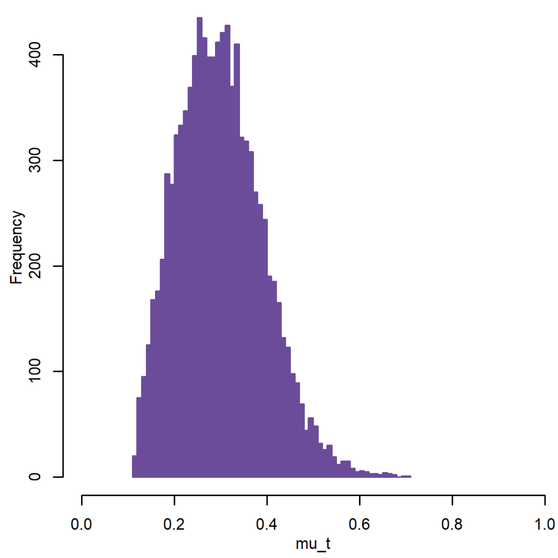


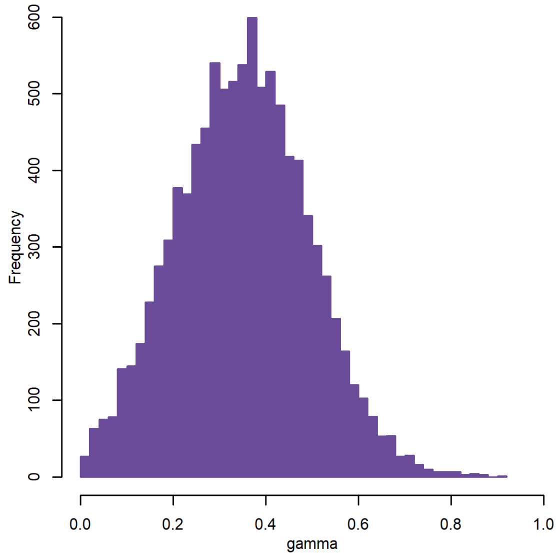

### Cohort 15

Author: Hartley

Publication year: 1935

Patient recruitment years: 1905-1914

Cohort size: 944

Location: England

Type of TB (smear-status): positive

Details: Retrospective cohort study of cases treated for TB at Brompton Hospital. Male patients only.

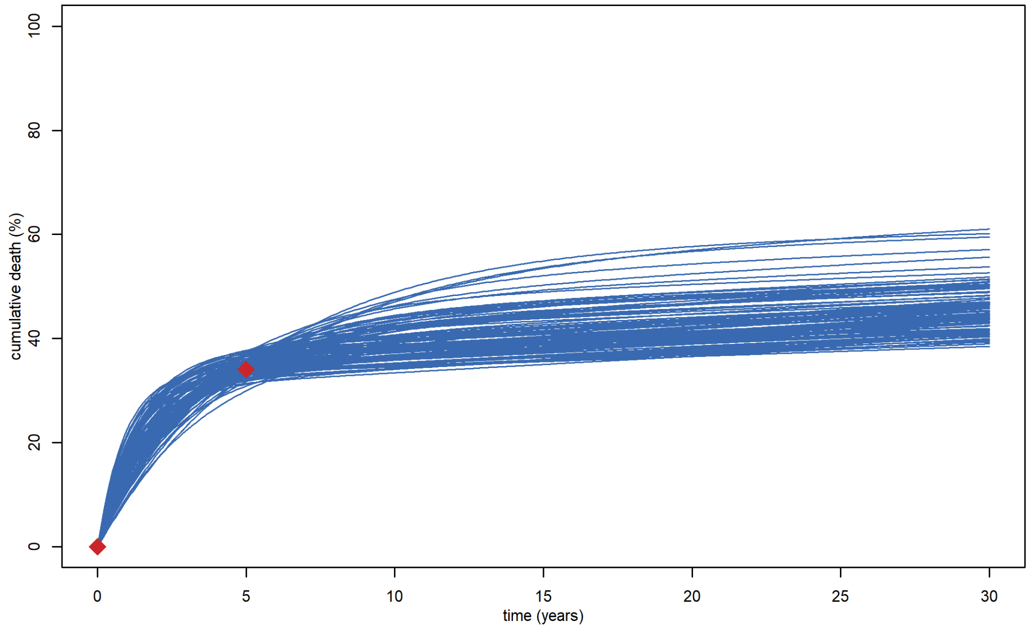


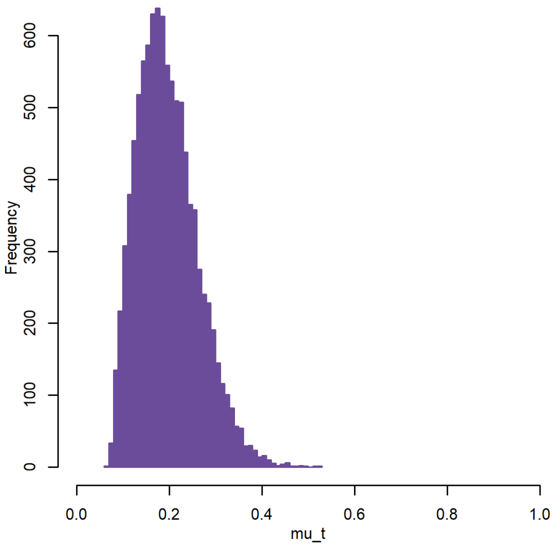


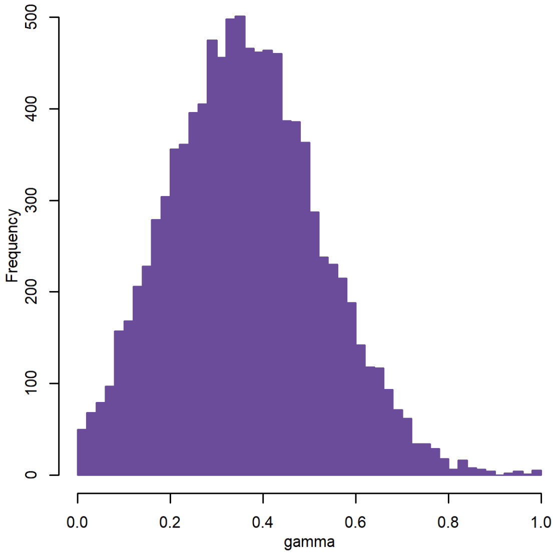

### Cohort 16

Author: Braeuning

Publication year: 1936

Patient recruitment years: 1920-1921

Cohort size: 607

Location: Poland

Type of TB (smear-status): positive

Details: TB dispensary patients from Szczecin, Poland (then known as Stettin, Germany).

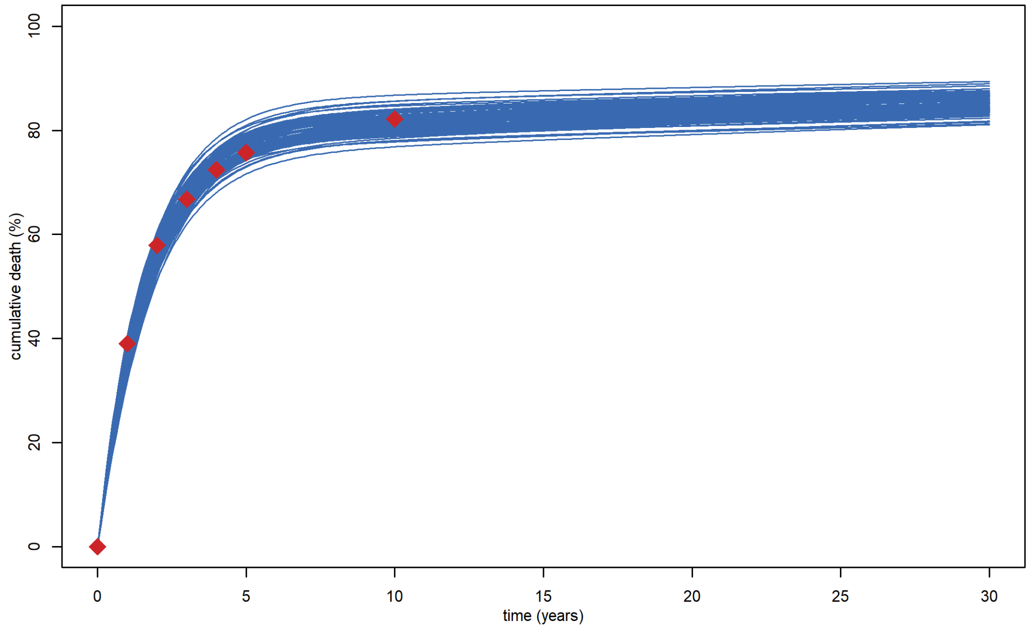


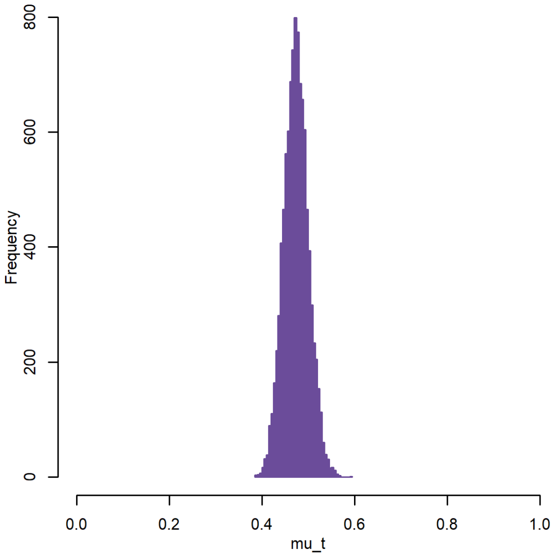


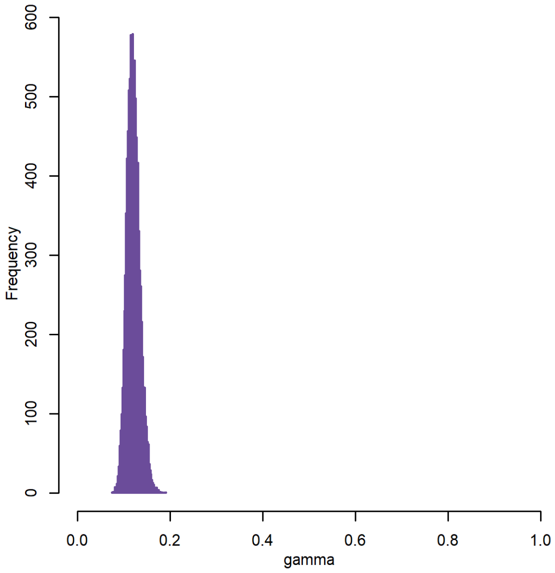

### Cohort 17

Author: Backer

Publication year: 1937

Patient recruitment years: 1911-1920

Cohort size: 1172

Location: Norway

Type of TB (smear-status): positive

Details: Patients notified to the Board of Health in Oslo.

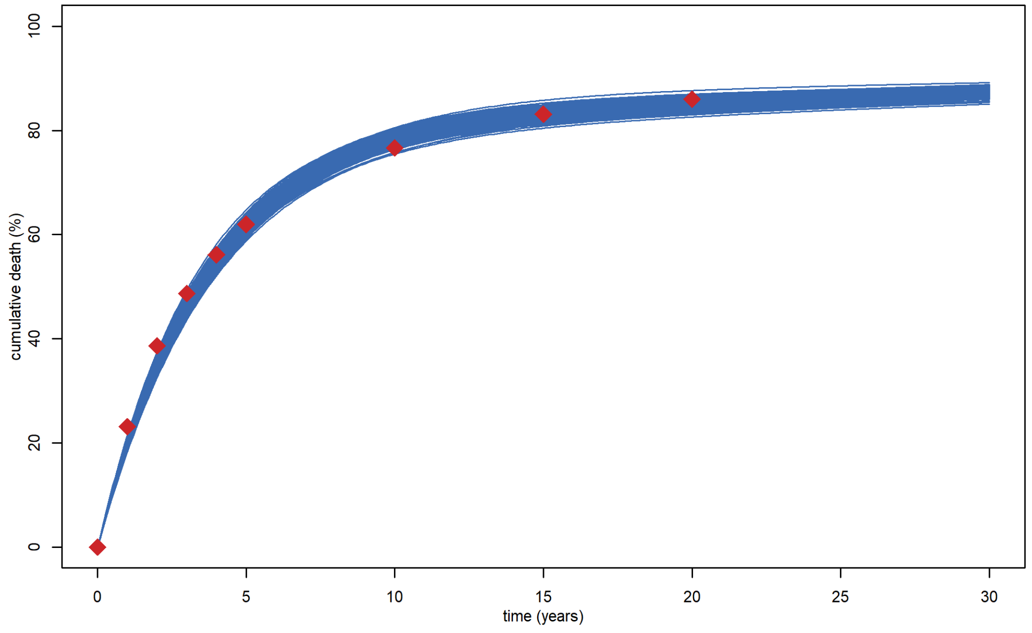


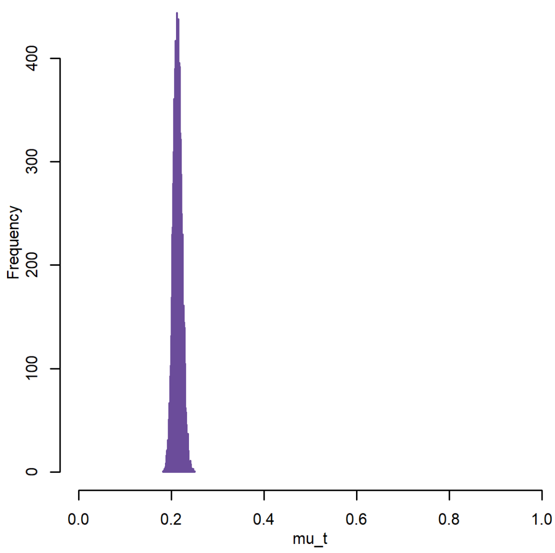


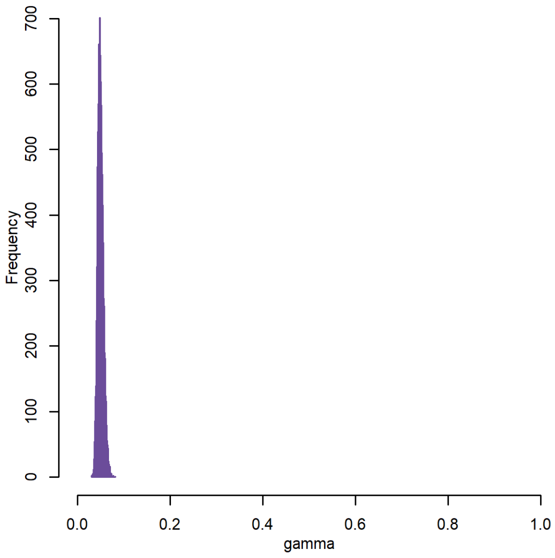

### Cohort 18

Author: Backer

Publication year: 1937

Patient recruitment years: 1911-1920

Cohort size: 1140

Location: Norway

Type of TB (smear-status): positive

Details: Patients notified to the Board of Health in Oslo. Male patients only.

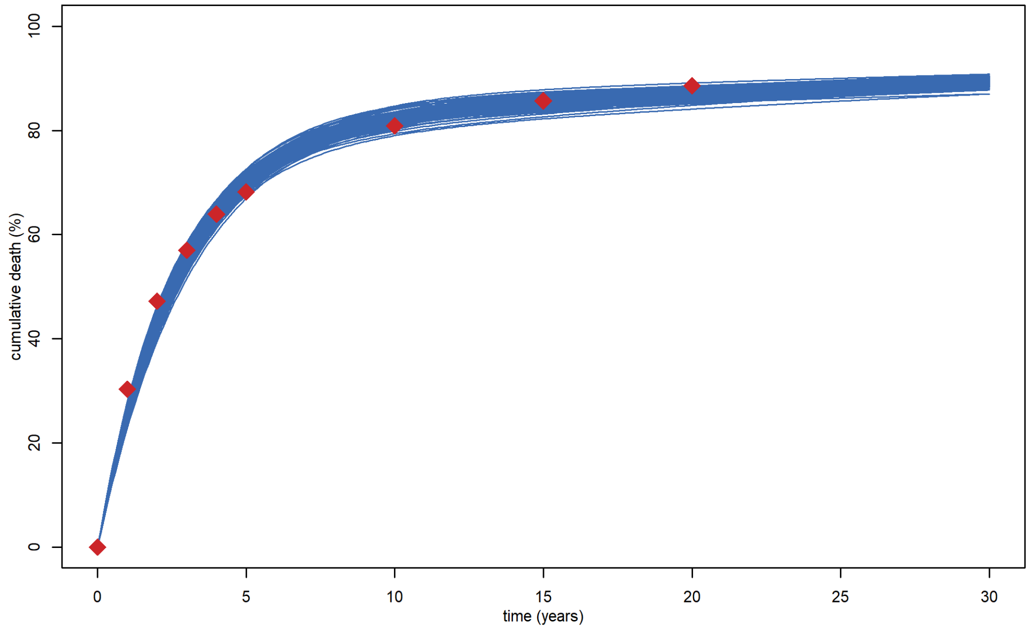


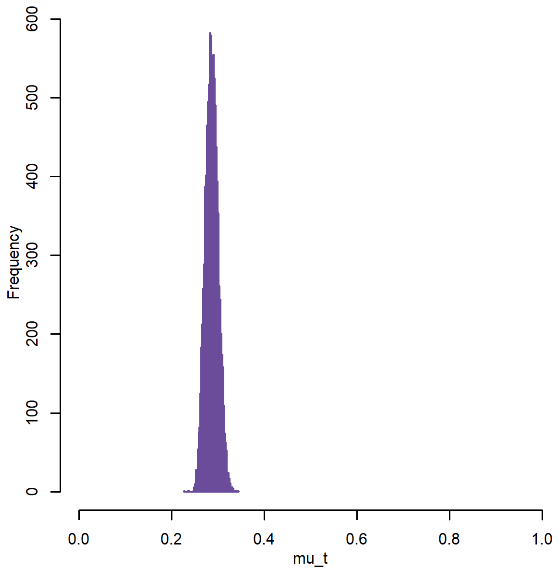


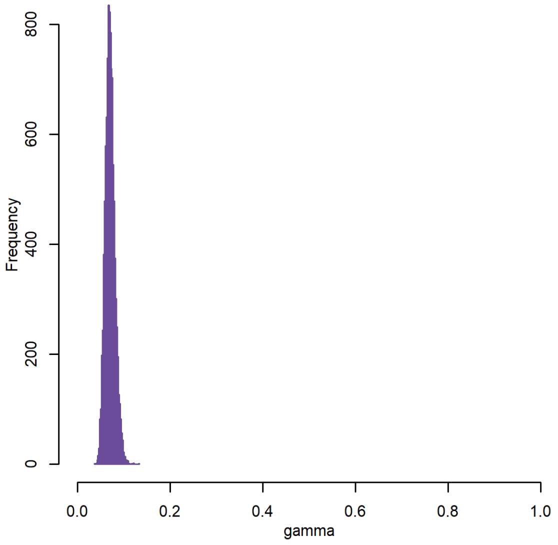

### Cohort 19

Author: Trail

Publication year: 1931

Patient recruitment years: 1911-1928

Cohort size: 1671

Location: England

Type of TB (smear-status): positive

Details: Cohort study among patients of the King Edward VII sanatorium in Midhurst.

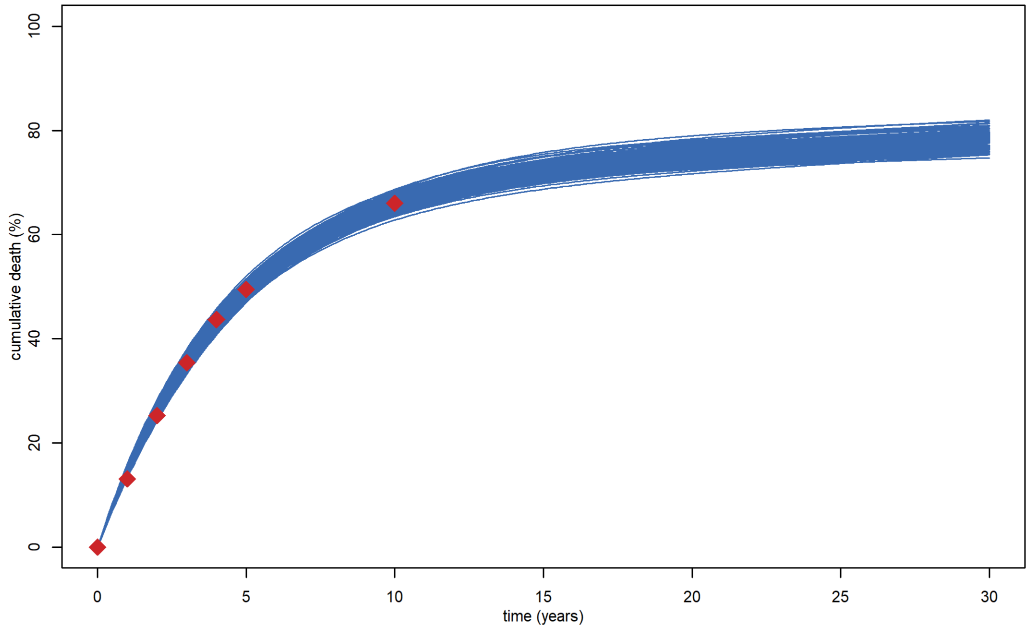


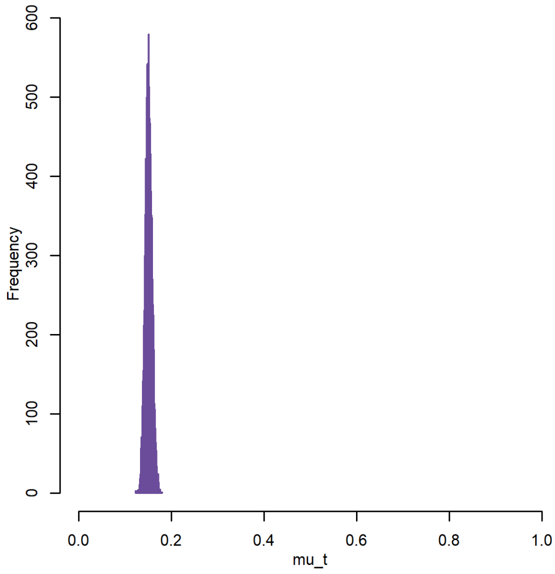


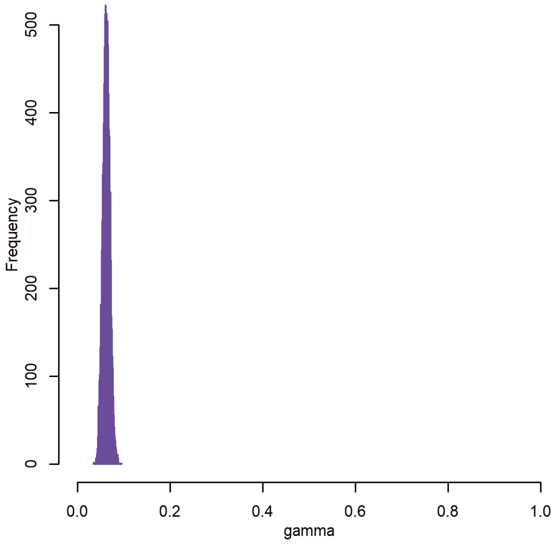

### Cohort 20

Author: Trail

Publication year: 1931

Patient recruitment years: 1911-1928

Cohort size: 944

Location: England

Type of TB (smear-status): positive

Details: Cohort study among patients of the King Edward VII sanatorium in Midhurst. Male patients only.

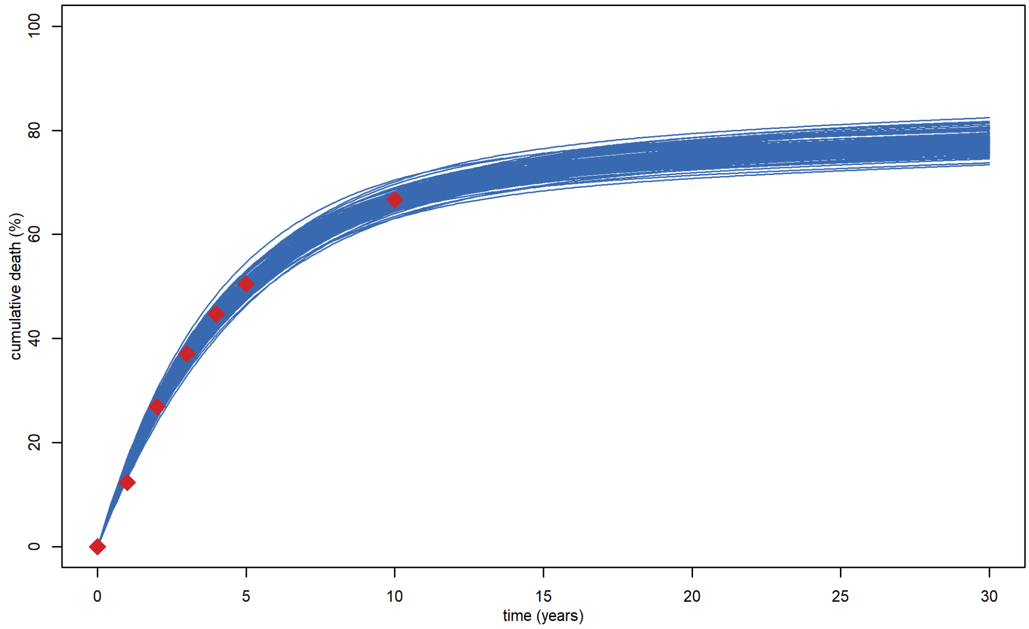


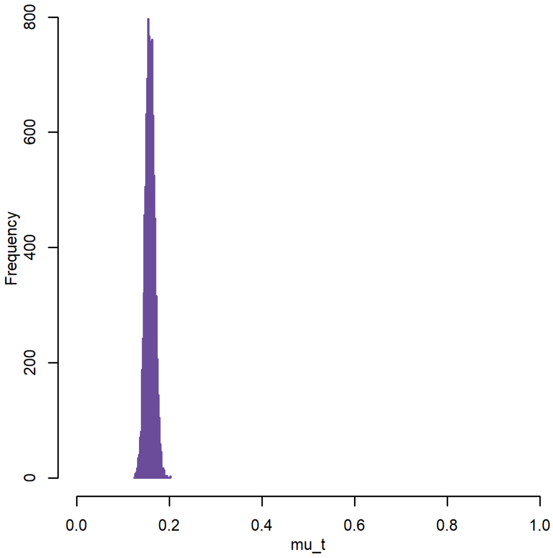


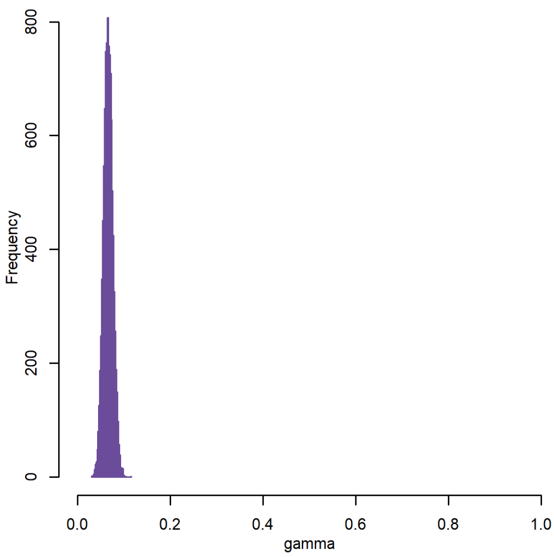

### Cohort 21

Author: Sinding-Larsen

Publication year: 1937

Patient recruitment years: 1906-1932

Cohort size: 1114

Location: Denmark

Type of TB (smear-status): positive

Details: Cohort study in Denmark among sanatorium patients.

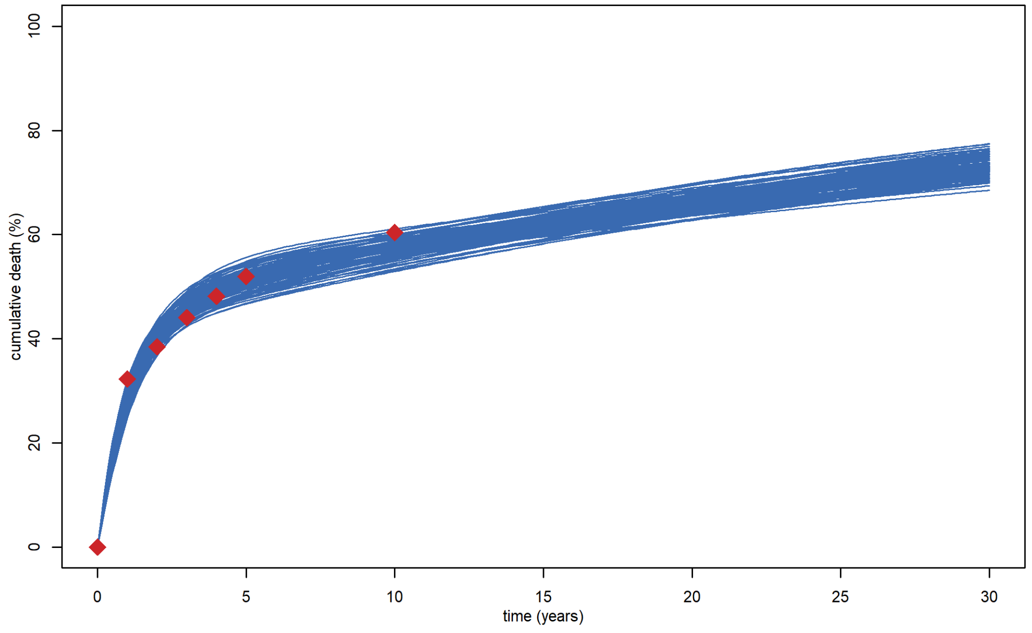


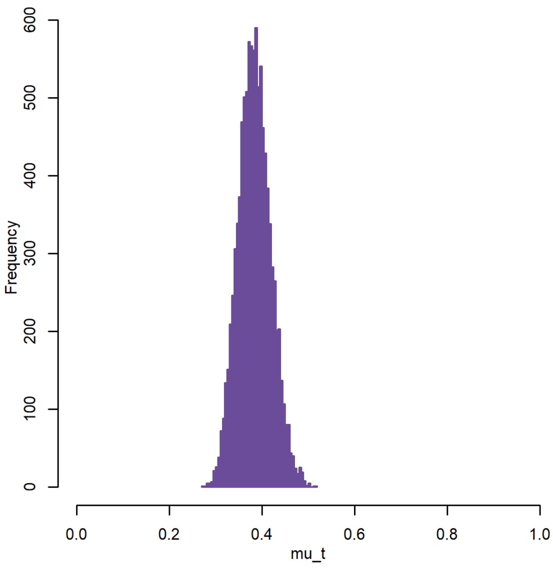


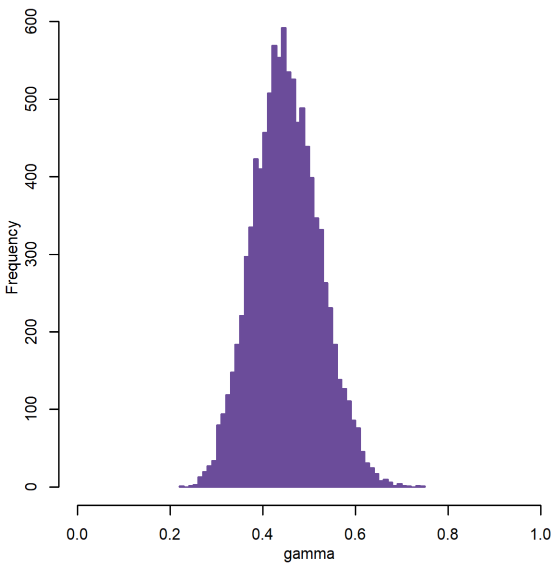

### Cohort 22

Author: Furth

Publication year: 1930

Patient recruitment years: 1912-1915

Cohort size: 303

Location: Switzerland

Type of TB (smear-status): positive

Details: Pulmonary TB patients
from Barmelweid sanatorium.

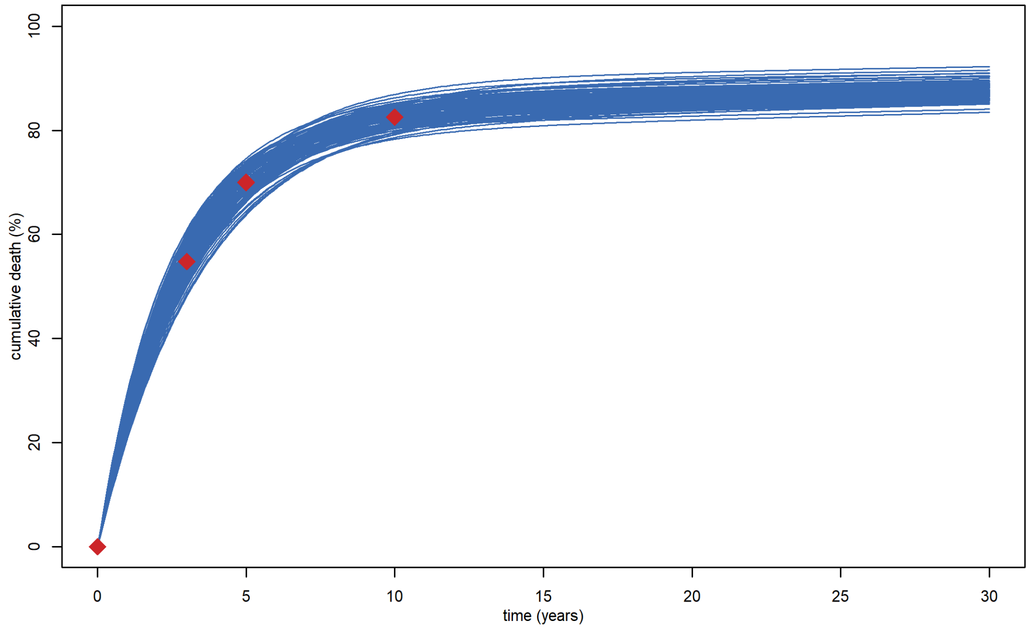


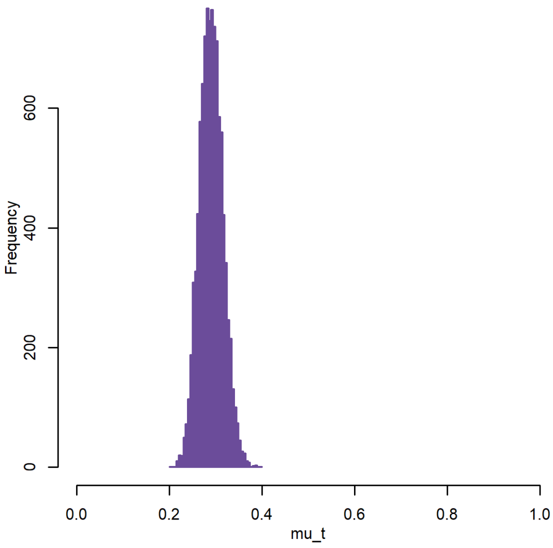


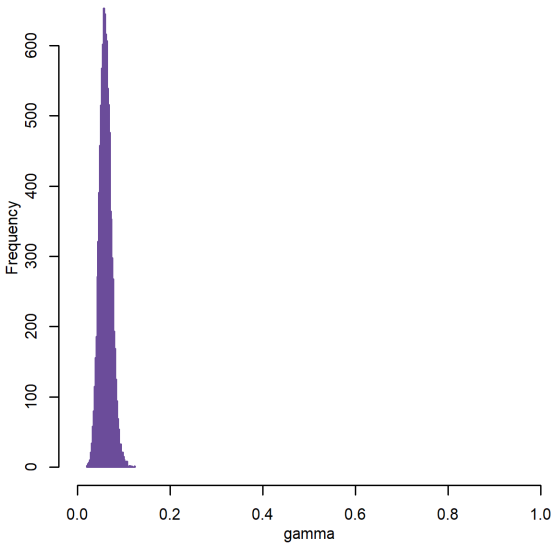

### Cohort 23

Author: Furth

Publication year: 1930

Patient recruitment years: 1916-1919

Cohort size: 307

Location: Switzerland

Type of TB (smear-status): positive

Details: Pulmonary TB patients
from Barmelweid sanatorium.

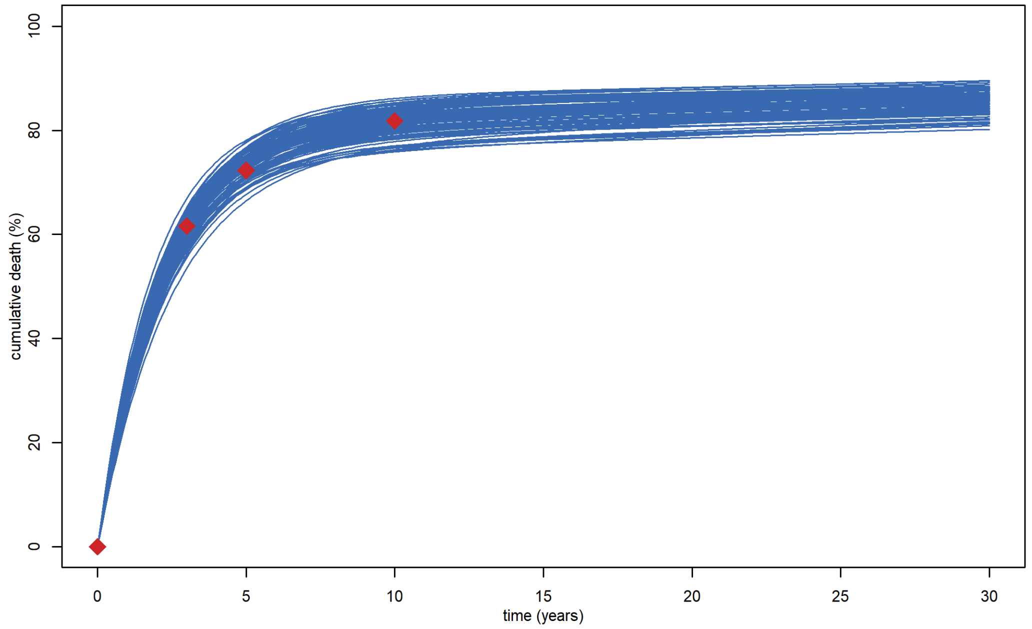


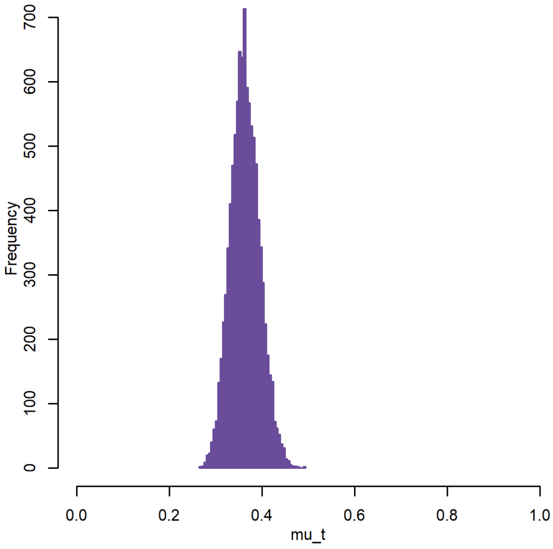


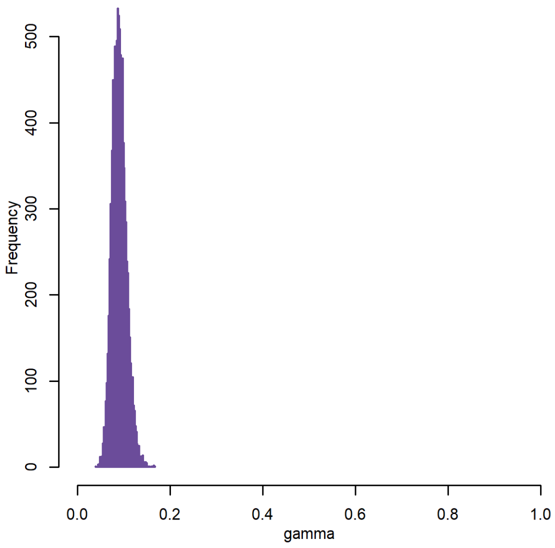

### Cohort 24

Author: Furth

Publication year: 1930

Patient recruitment years: 1920-1924

Cohort size: 386

Location: Switzerland

Type of TB (smear-status): positive

Details: Pulmonary TB patients
from Barmelweid sanatorium.

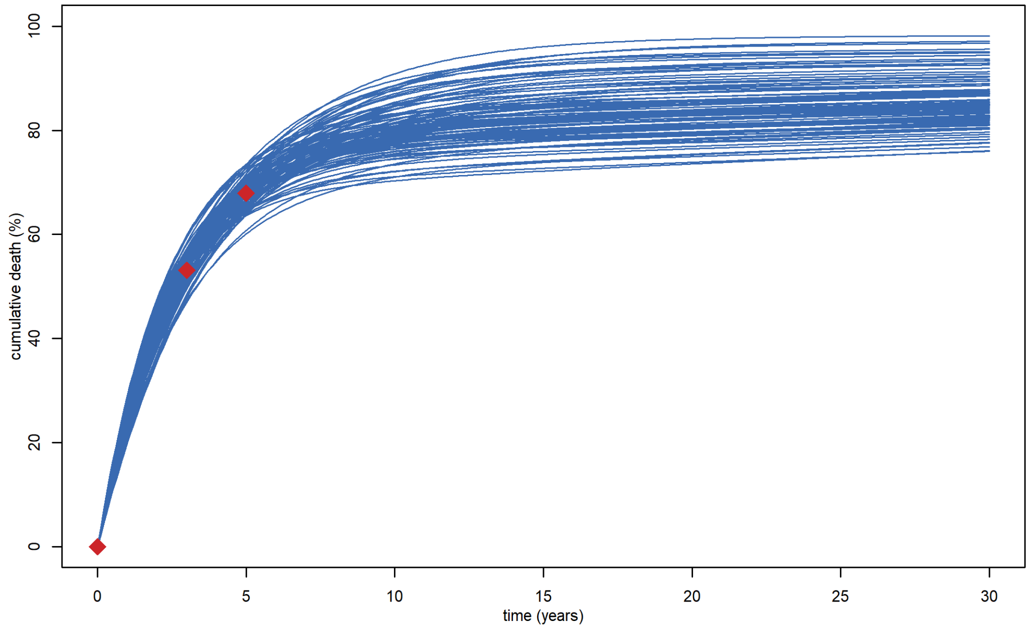


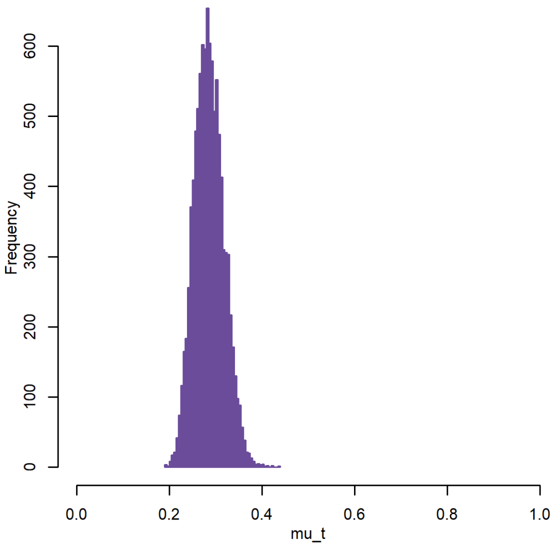


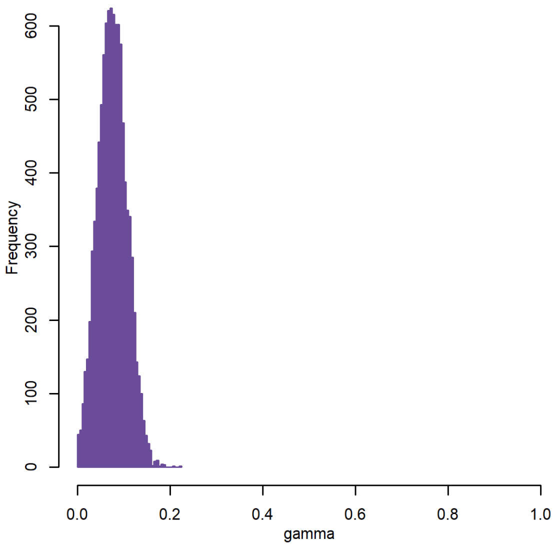

### Cohort 38

Author: Berg

Publication year: 1939

Patient recruitment years: 1928-1934

Cohort size: 2042

Location: Sweden

Type of TB (smear-status): positive

Details: All patients with "open" TB from Gothenburg diagnosed between 1928 and 1934.

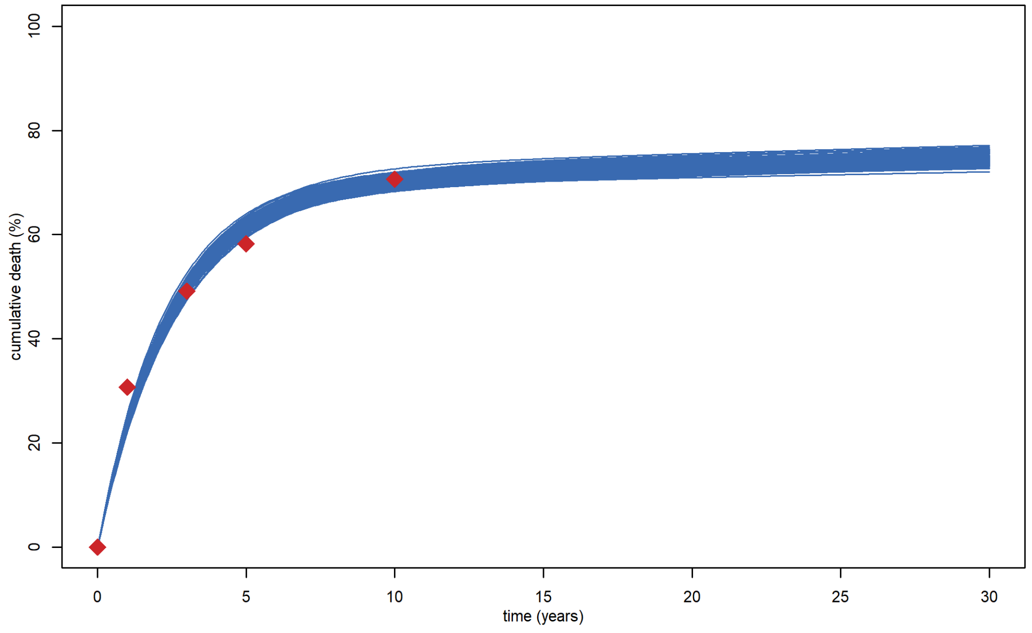


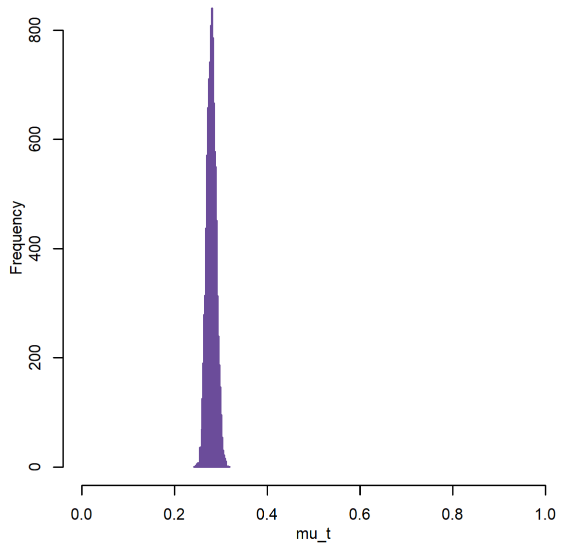


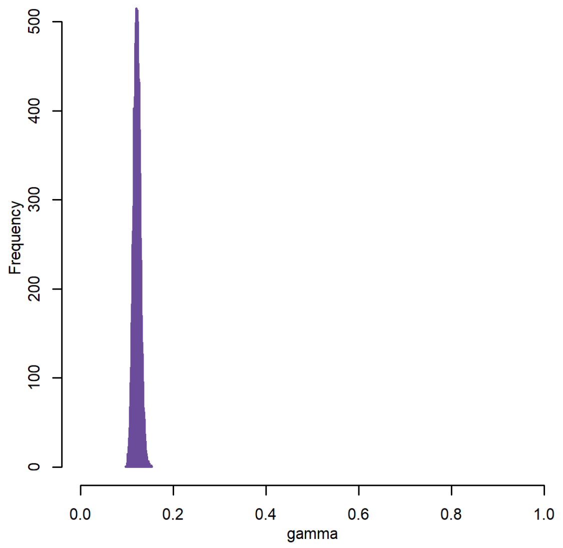

### Cohort 39

Author: Munchbach

Publication year: 1939 (from Berg et al.)

Patient recruitment years: 1920-1921

Cohort size: 266

Location: Germany

Type of TB (smear-status): positive

Details: Sanatorium patients with "open" bacillary TB. Male patients only.

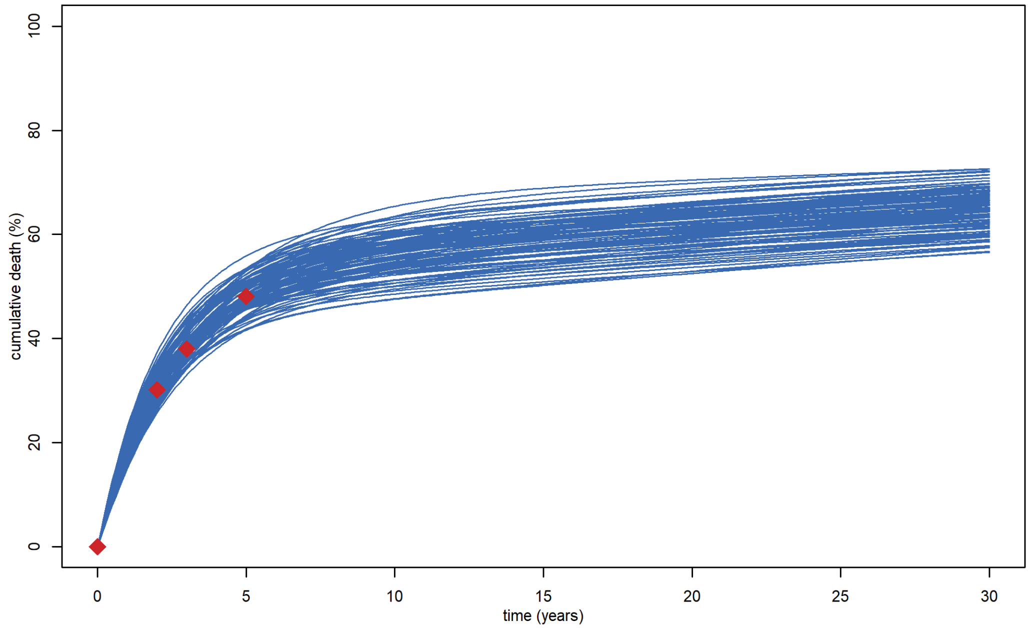


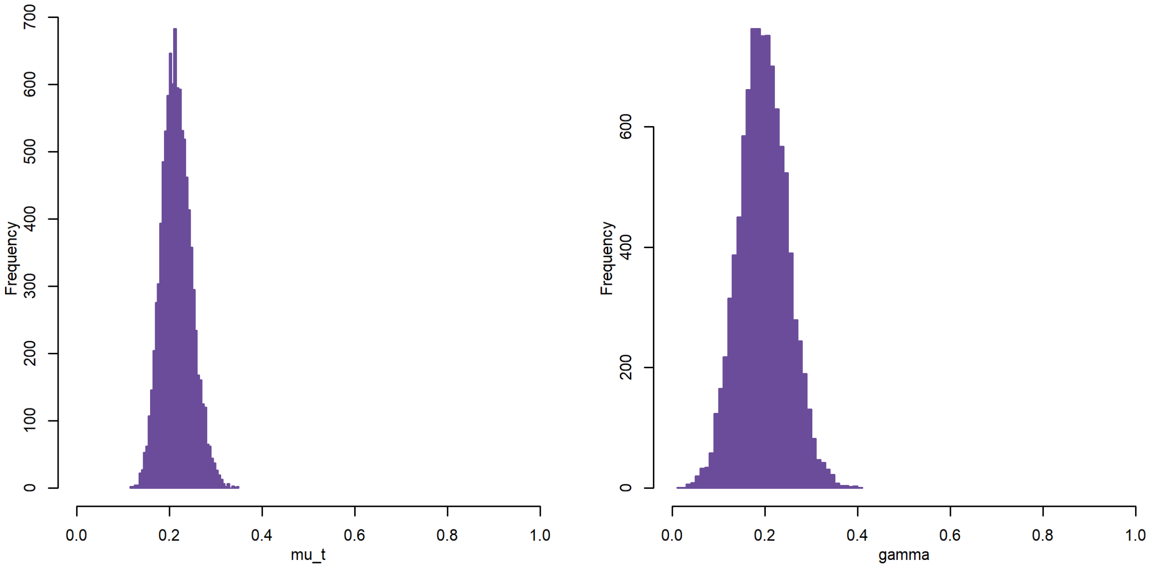

### Cohort 40

Author: Munchbach

Publication year: 1939 (from Berg et al.)

Patient recruitment years: 1922-1923

Cohort size: 681

Location: Germany

Type of TB (smear-status): positive

Details: Sanatorium patients with "open" bacillary TB. Male patients only.

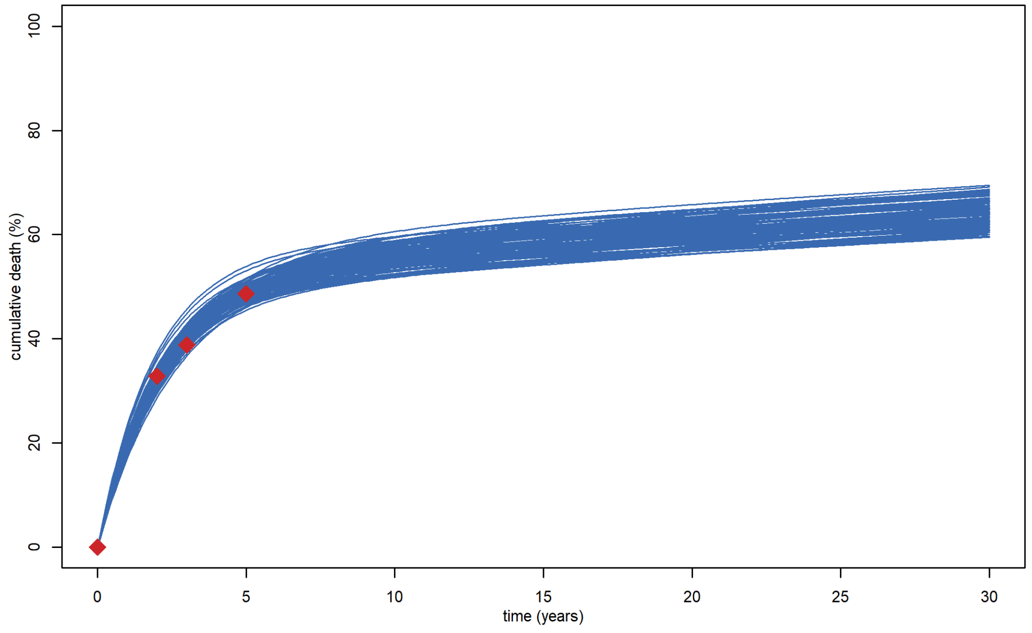


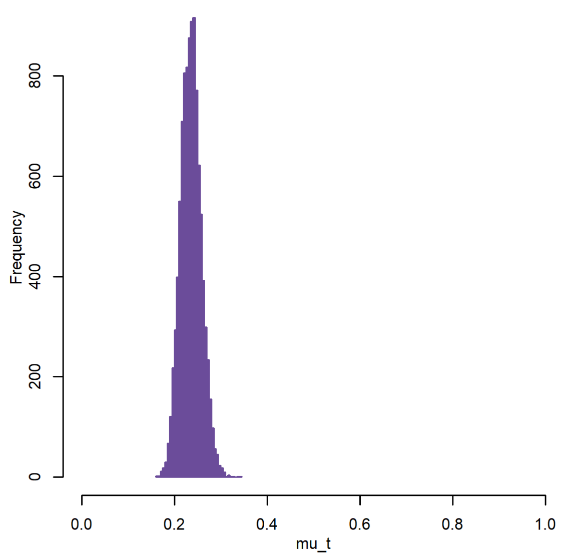


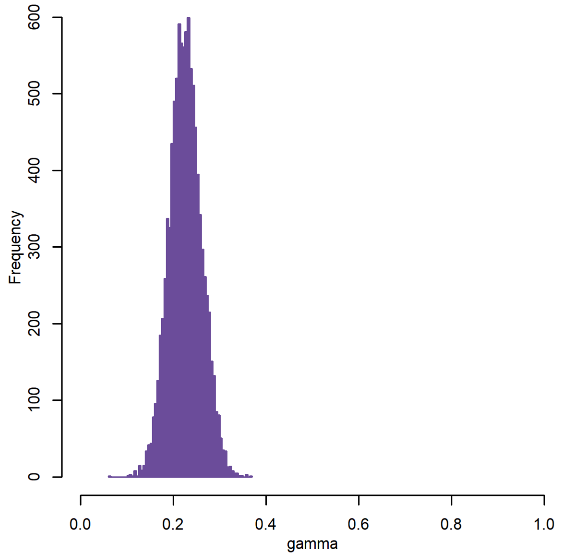

### Cohort 41

Author: Munchbach

Publication year: 1939 (from Berg et al.)

Patient recruitment years: 1924-1925

Cohort size: 504

Location: Germany

Type of TB (smear-status): positive

Details: Sanatorium patients with "open" bacillary TB. Male patients only.

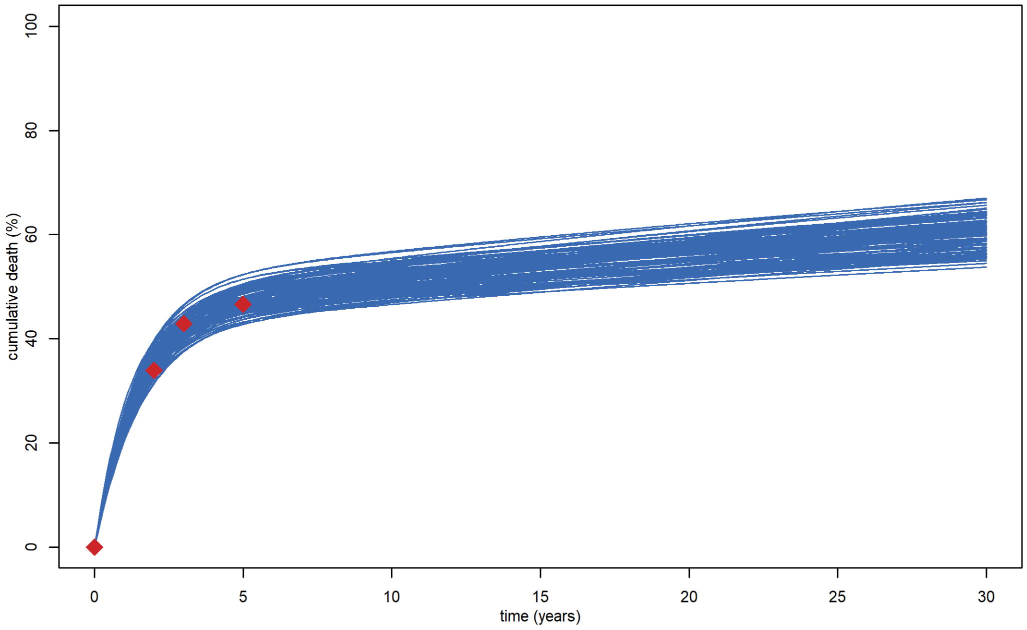


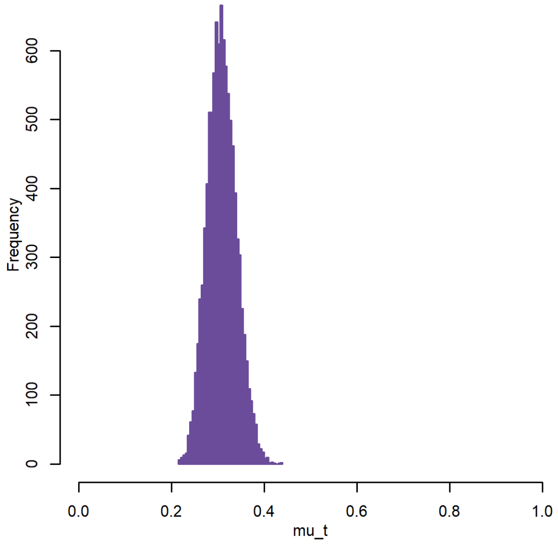


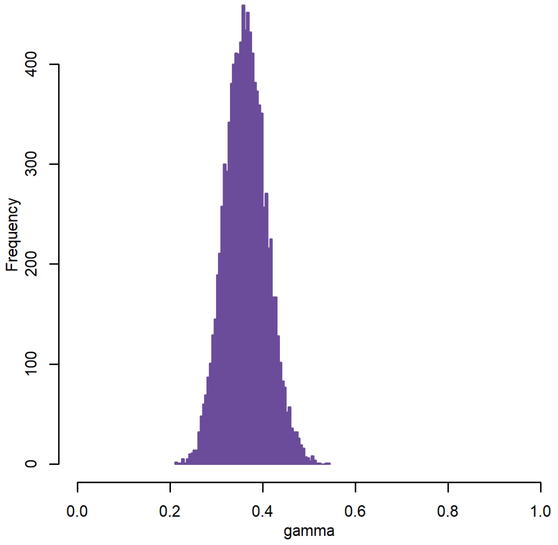

### Cohort 42

Author: Munchbach

Publication year: 1939 (from Berg et al.)

Patient recruitment years: 1926-1927

Cohort size: 660

Location: Germany

Type of TB (smear-status): positive

Details: Sanatorium patients with "open" bacillary TB. Male patients only.

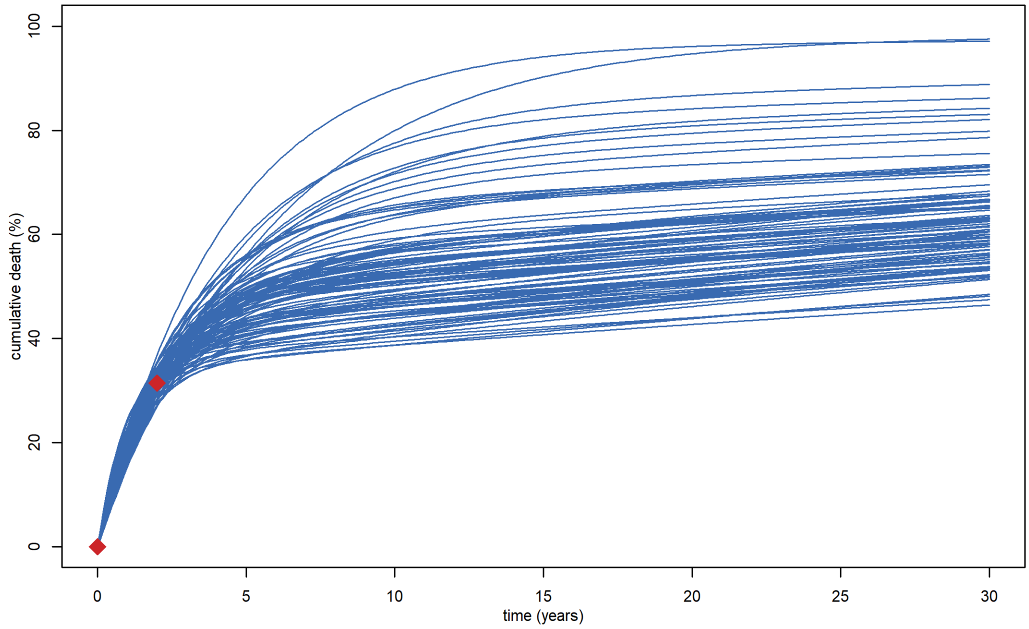


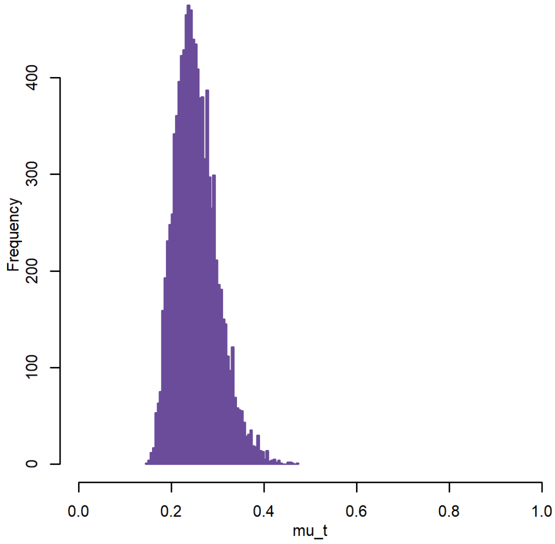


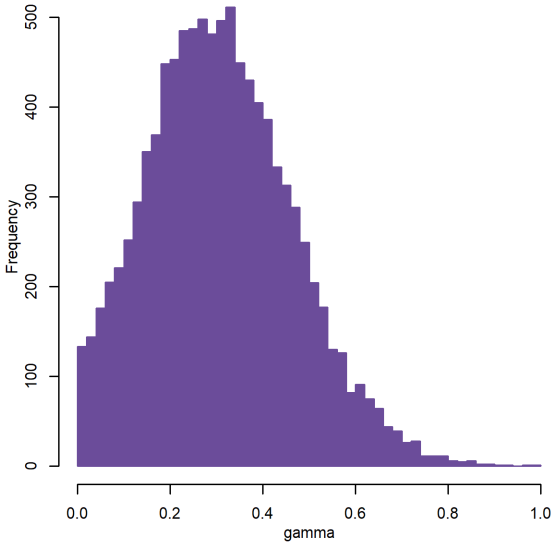

### Cohort 43

Author: Munchbach

Publication year: 1939 (from Berg et al.)

Patient recruitment years: 1920-1921

Cohort size: 428

Location: Germany

Type of TB (smear-status): positive

Details: Sanatorium patients with "open" bacillary TB. Female patients only.

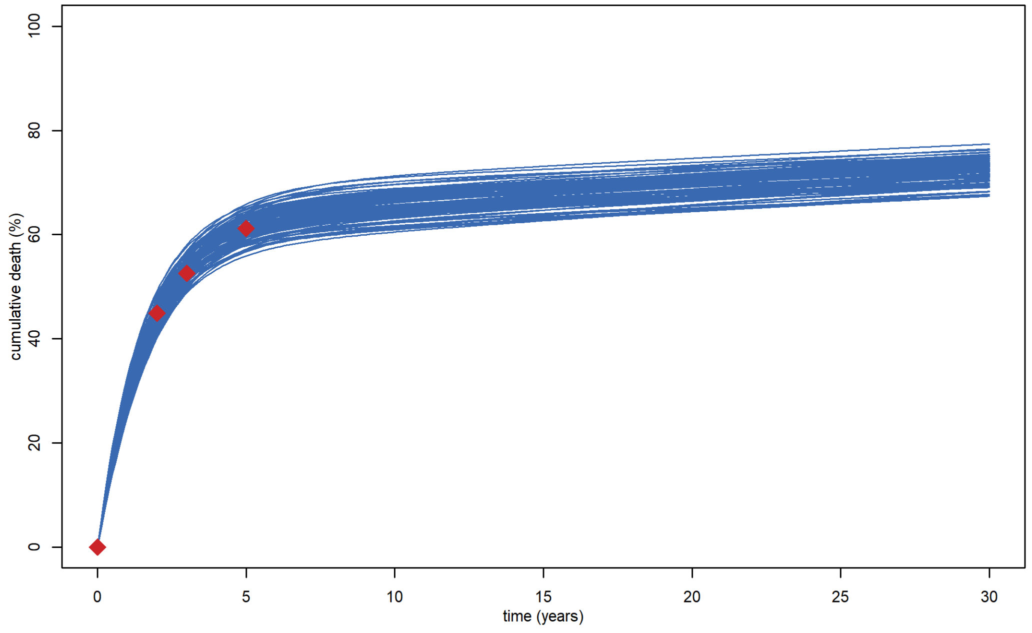


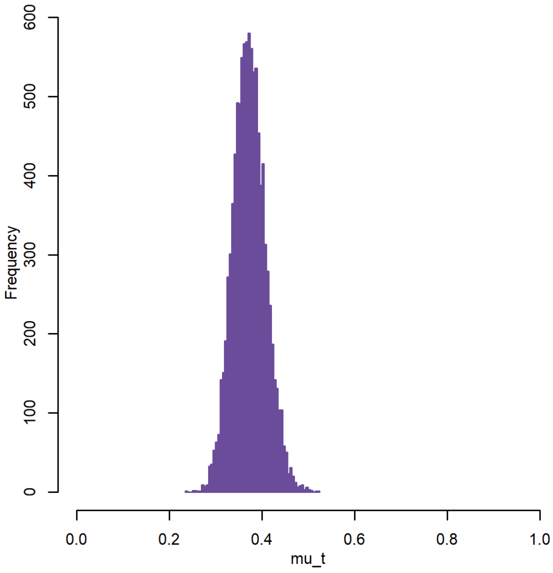


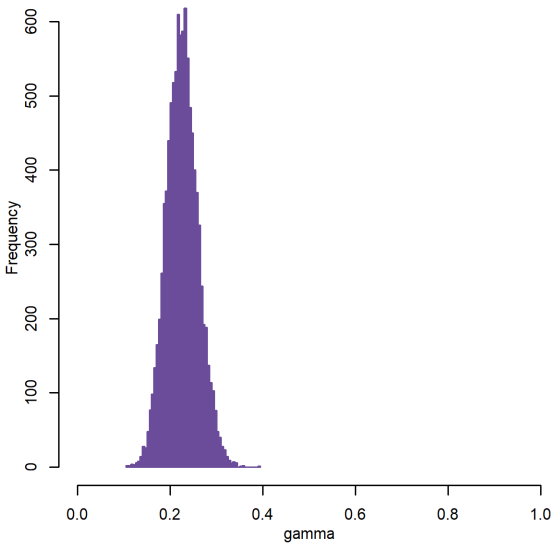

### Cohort 44

Author: Munchbach

Publication year: 1939 (from Berg et al.)

Patient recruitment years: 1922-1923

Cohort size: 497

Location: Germany

Type of TB (smear-status): positive

Details: Sanatorium patients with "open" bacillary TB. Female patients only.

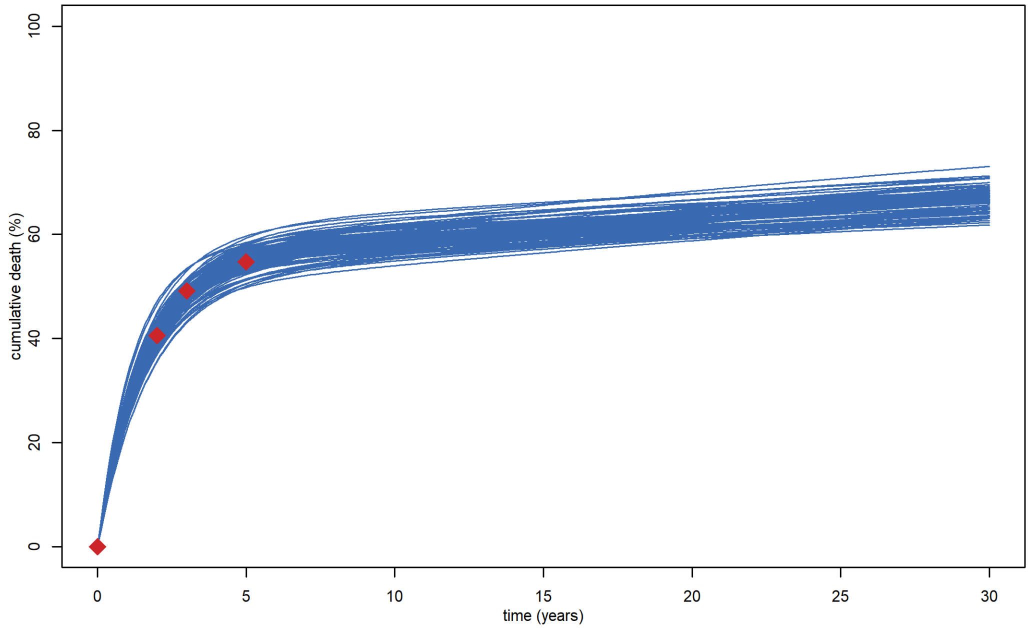


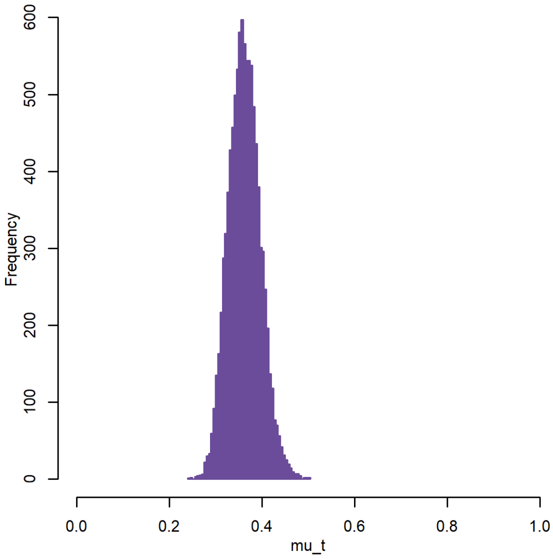


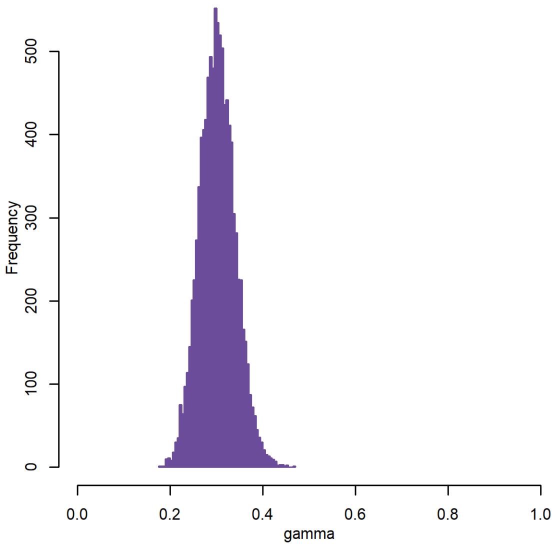

### Cohort 45

Author: Munchbach

Publication year: 1939 (from Berg et al.)

Patient recruitment years: 1924-1925

Cohort size: 433

Location: Germany

Type of TB (smear-status): positive

Details: Sanatorium patients with "open" bacillary TB. Female patients only.

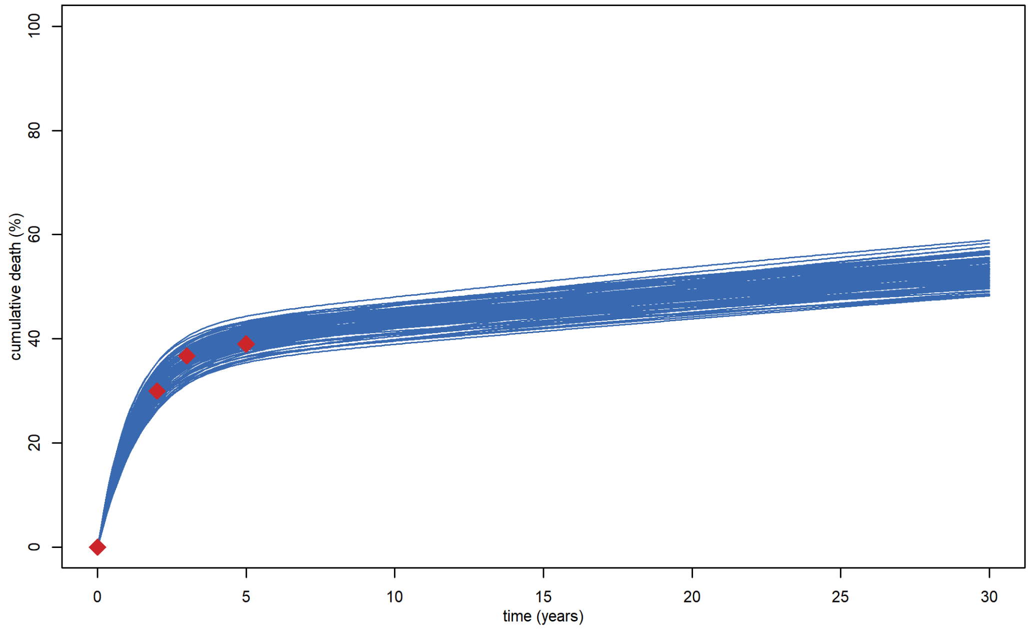


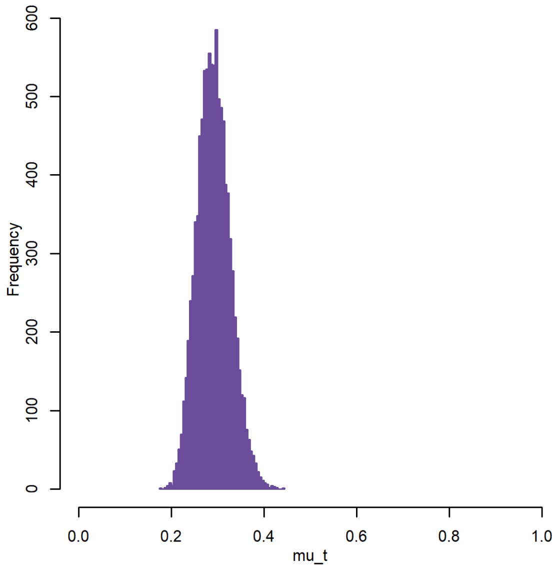


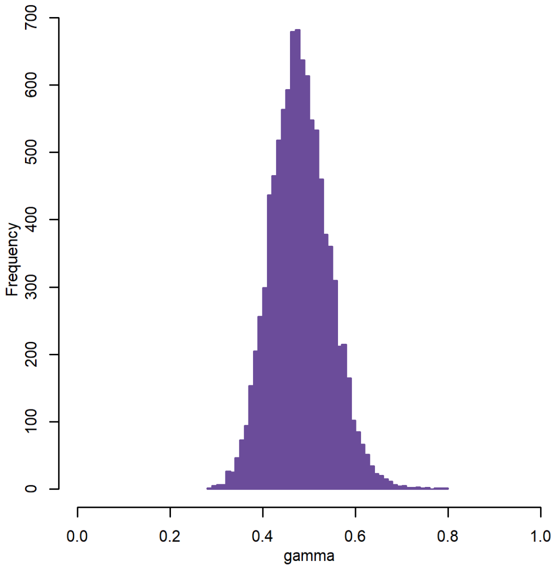

### Cohort 46

Author: Munchbach

Publication year: 1939 (from Berg et al.)

Patient recruitment years: 1926-1927

Cohort size: 497

Location: Germany

Type of TB (smear-status): positive

Details: Sanatorium patients with "open" bacillary TB. Female patients only.

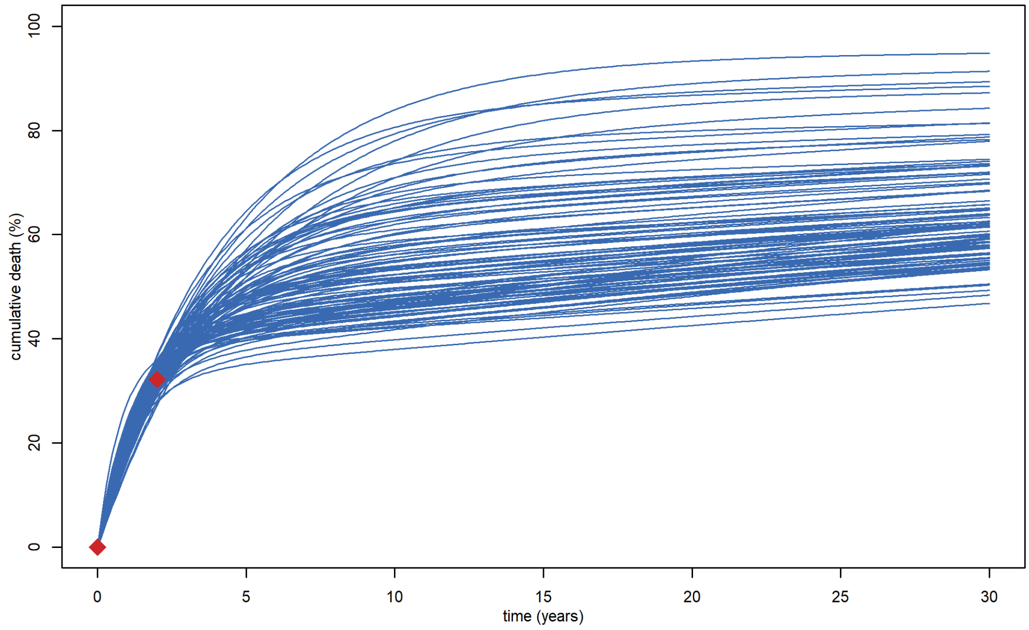


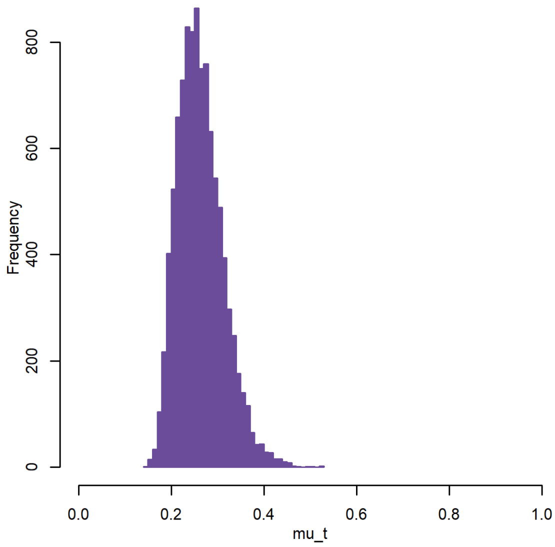


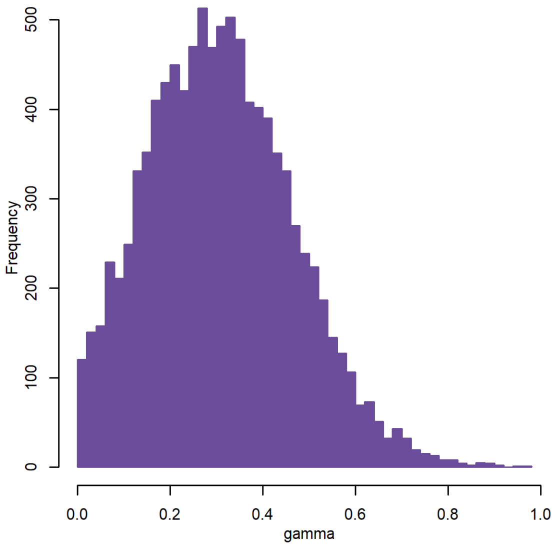

### Cohort 47

Author: Lindhart

Publication year: 1939

Patient recruitment years: 1930-1931

Cohort size: 1162

Location: Denmark

Type of TB (smear-status): positive

Details: Mortality of notified TB
cases in Denmark between 1925 and 1934.
Male patients only.

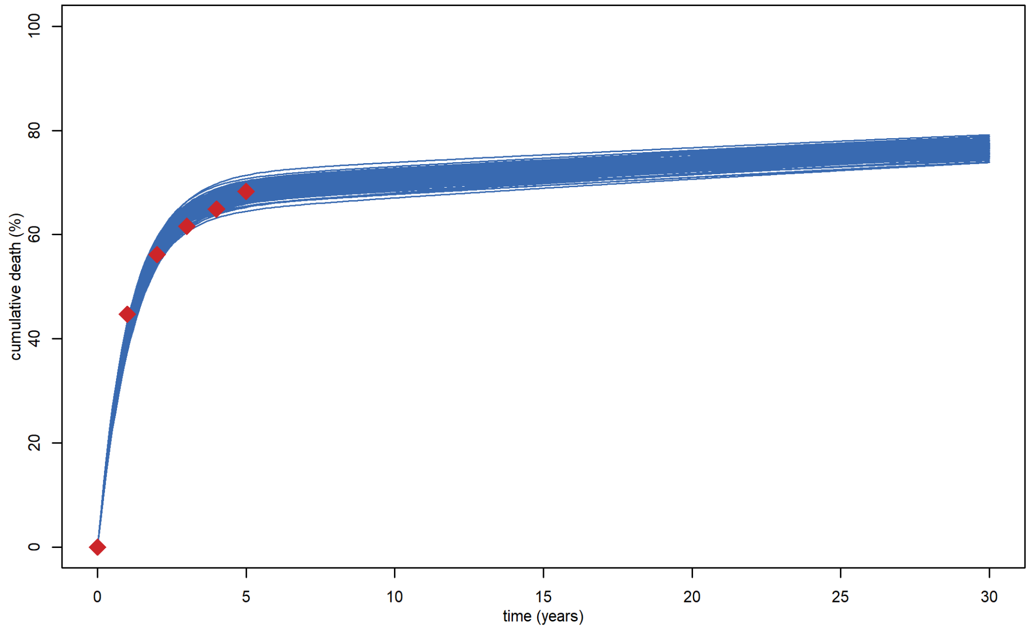


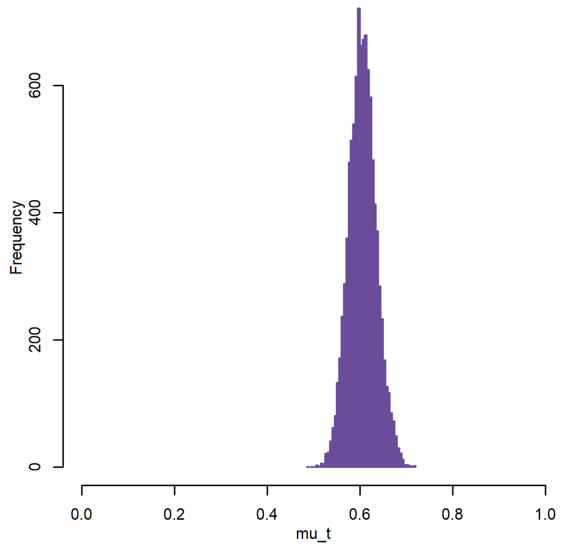


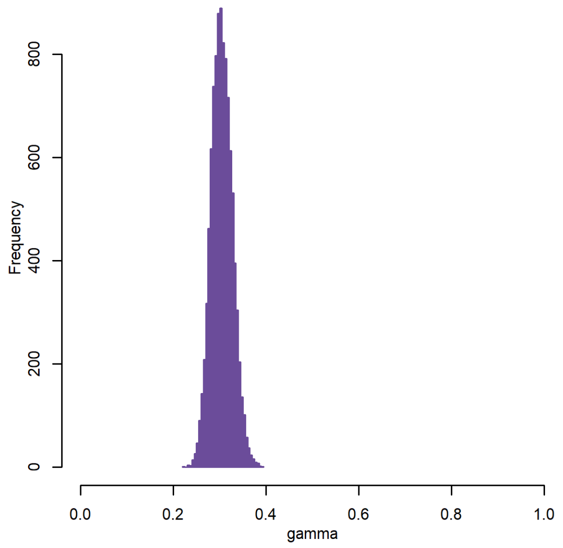

### Cohort 48

Author: Lindhart

Publication year: 1939

Patient recruitment years: 1931-1932

Cohort size: 1140

Location: Denmark

Type of TB (smear-status): positive

Details: Mortality of notified TB cases in Denmark between 1925 and 1934. Male patients only.

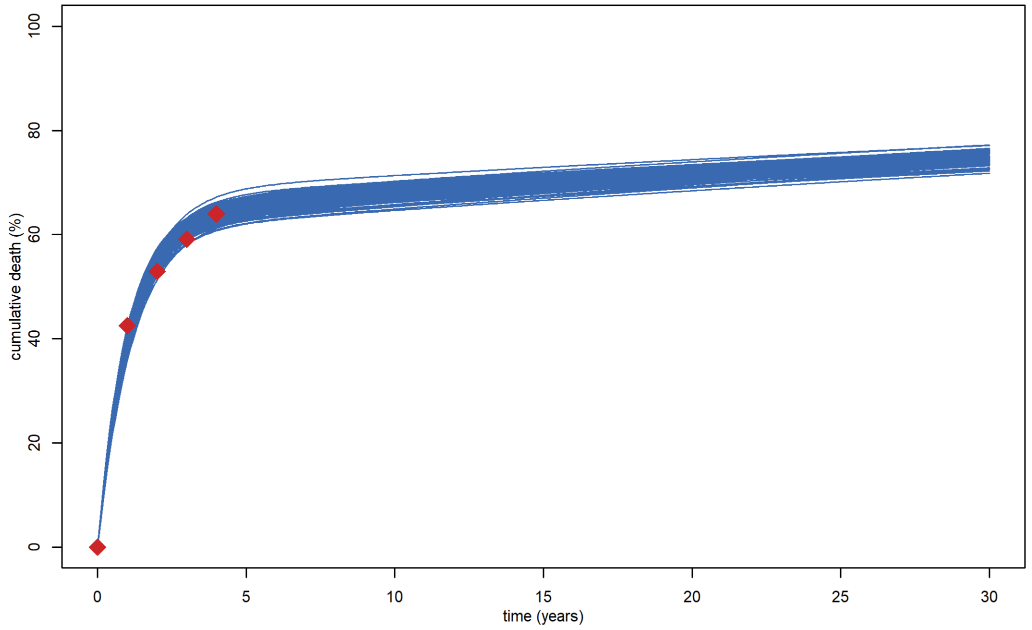


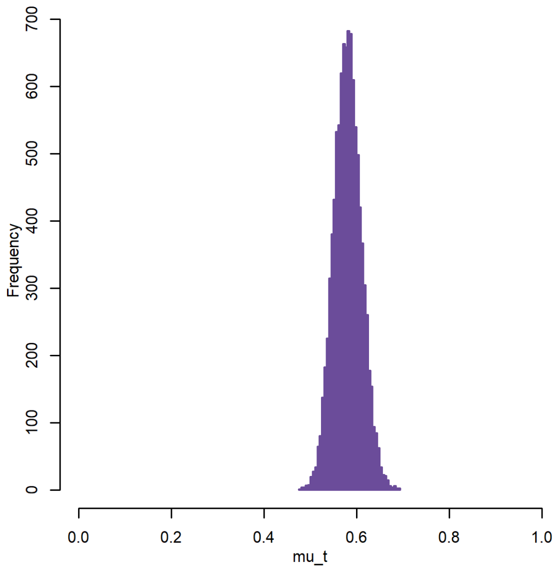


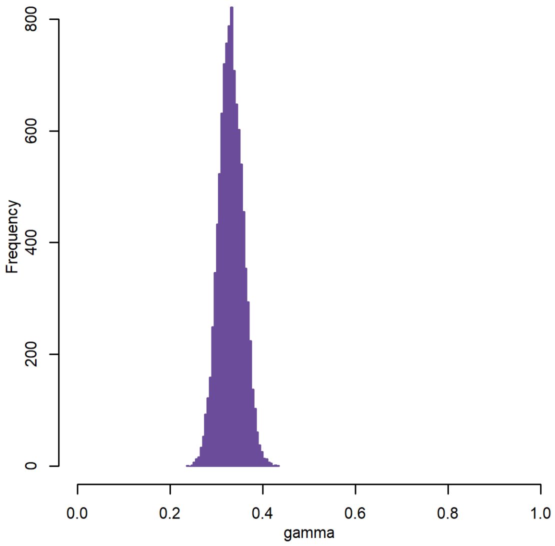

### Cohort 49

Author: Lindhart

Publication year: 1939

Patient recruitment years: 1932-1933

Cohort size: 1138

Location: Denmark

Type of TB (smear-status): positive

Details: Mortality of notified TB cases in Denmark between 1925 and 1934. Male patients only.

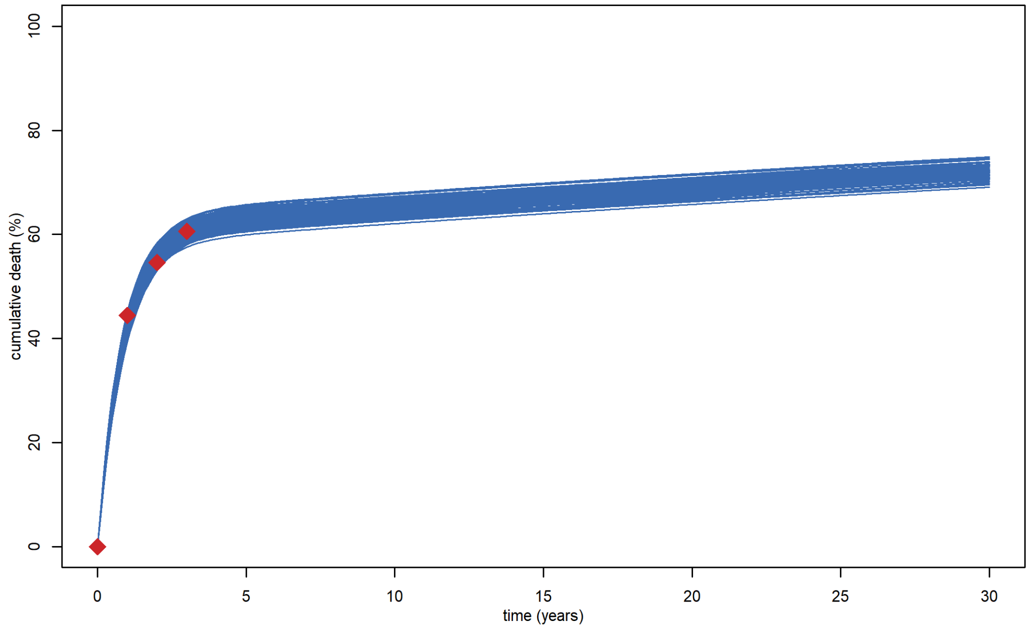


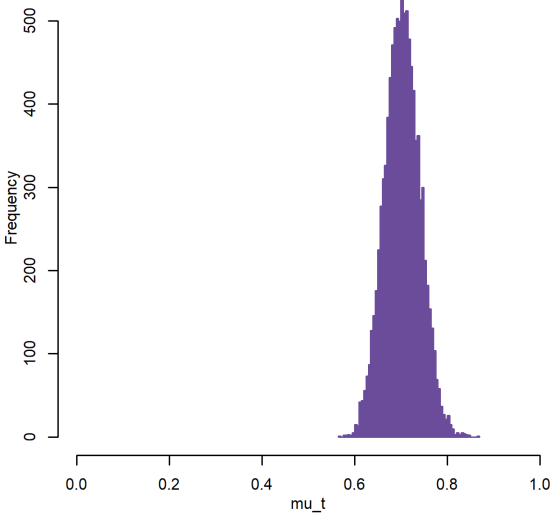


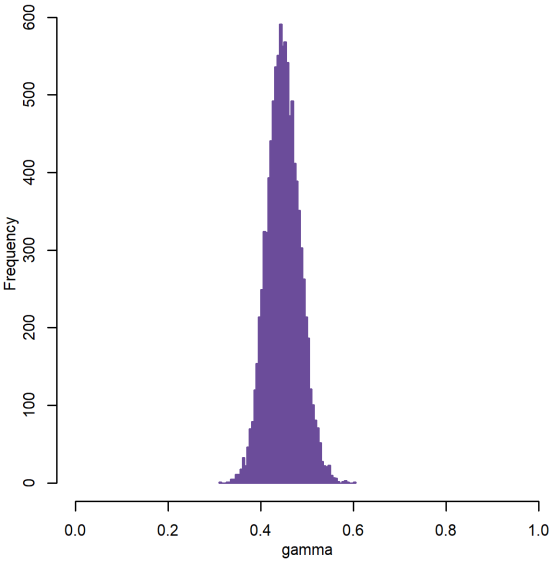

### Cohort 50

Author: Lindhart

Publication year: 1939

Patient recruitment years: 1933-1934

Cohort size: 1004

Location: Denmark

Type of TB (smear-status): positive

Details: Mortality of notified TB
cases in Denmark between 1925 and 1934.
Male patients only.

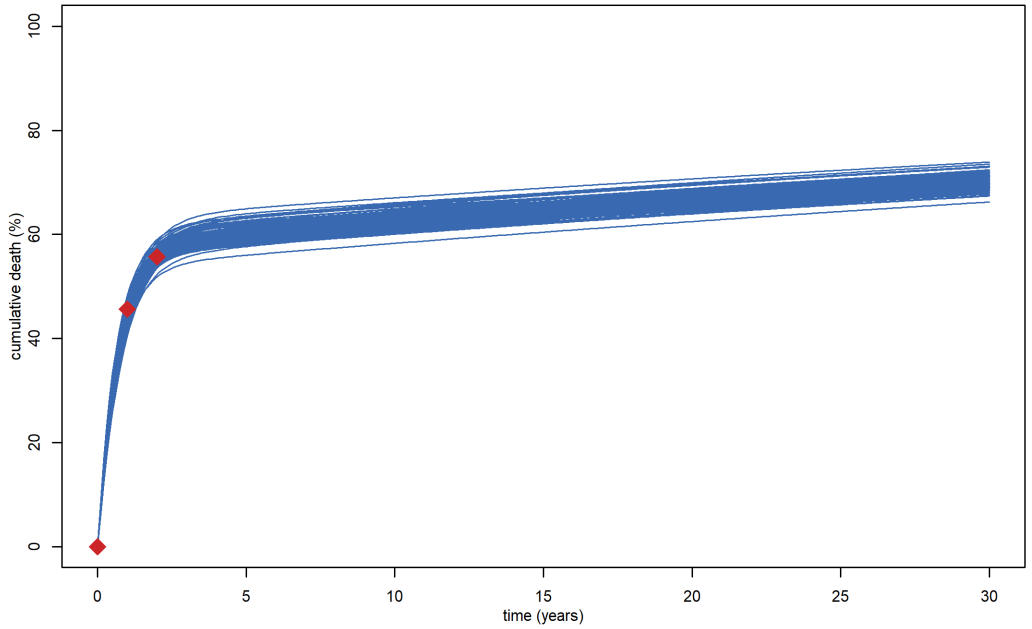


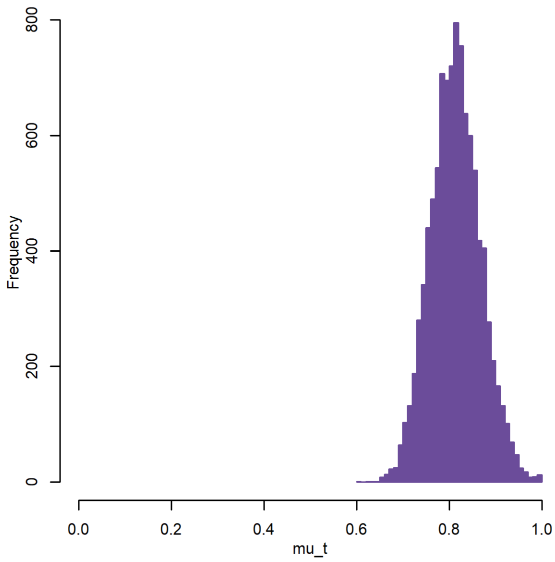


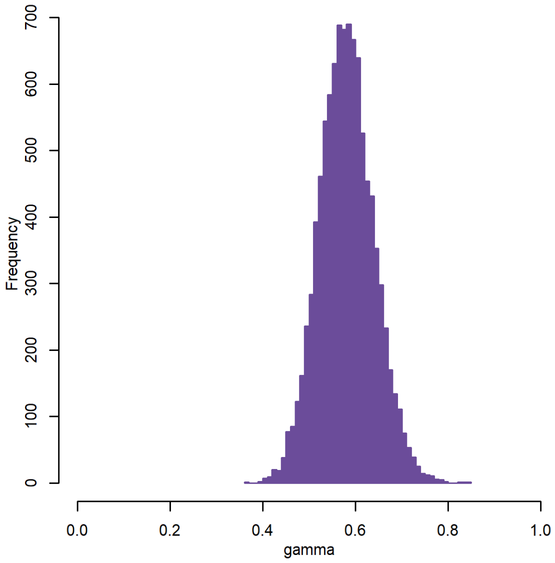

### Cohort 51

Author: Lindhart

Publication year: 1939

Patient recruitment years: 1934-1935

Cohort size: 968

Location: Denmark

Type of TB (smear-status): positive

Details: Mortality of notified TB cases in Denmark between 1925 and 1934. Male patients only.

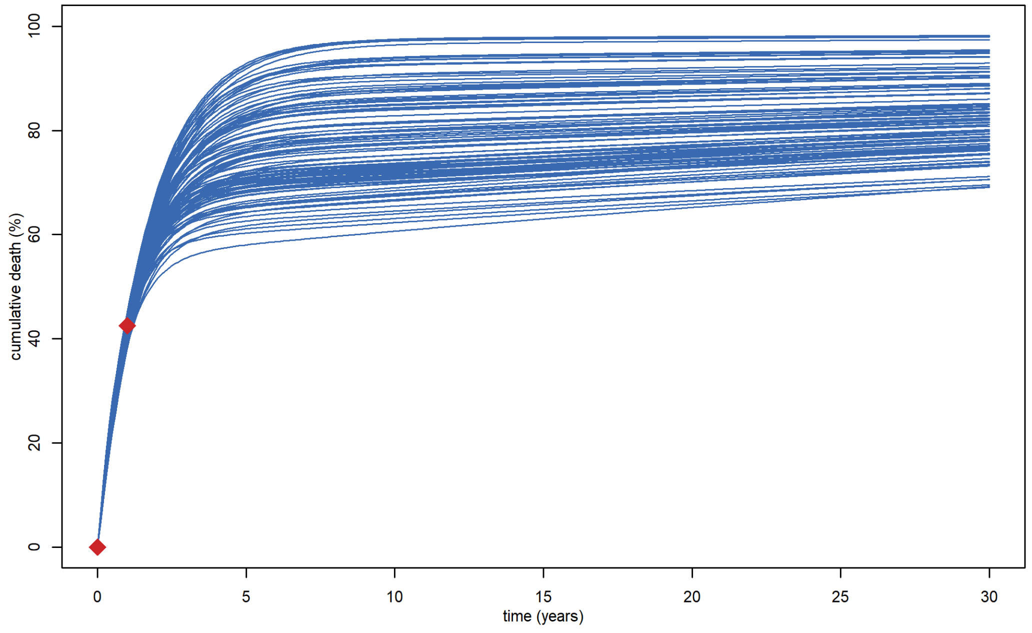


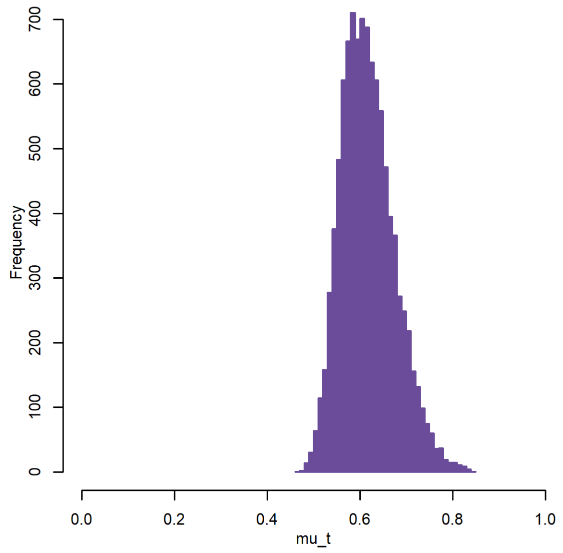


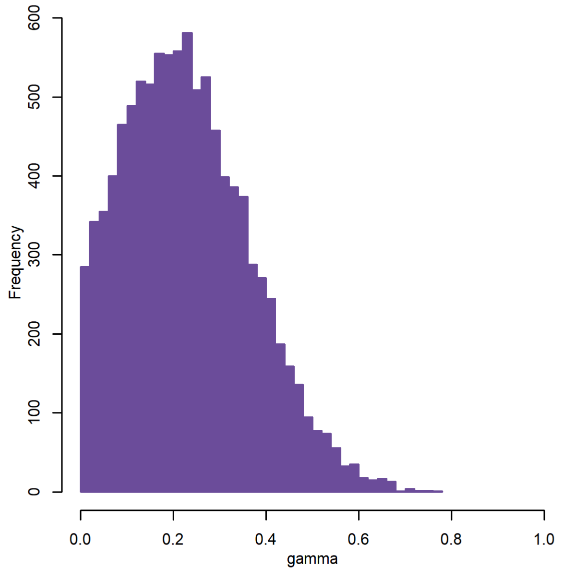

### Cohort 52

Author: Lindhart

Publication year: 1939

Patient recruitment years: 1930-1931

Cohort size: 1383

Location: Denmark

Type of TB (smear-status): positive

Details: Mortality of notified TB cases in Denmark between 1925 and 1934. Female patients only.

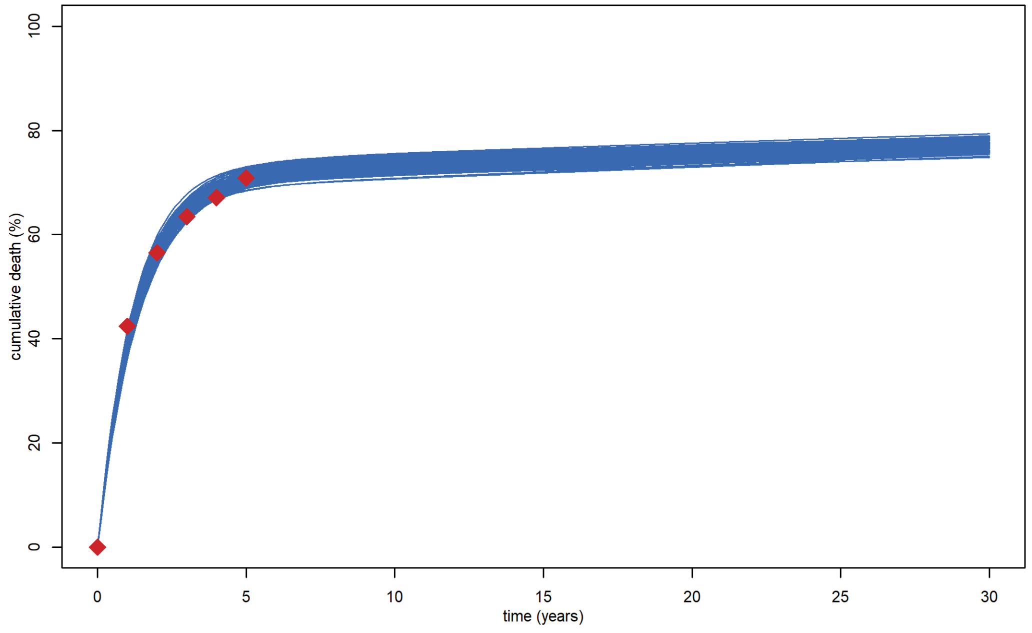


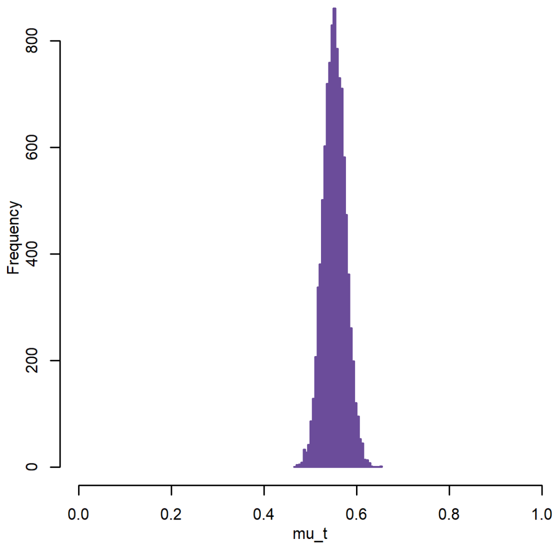


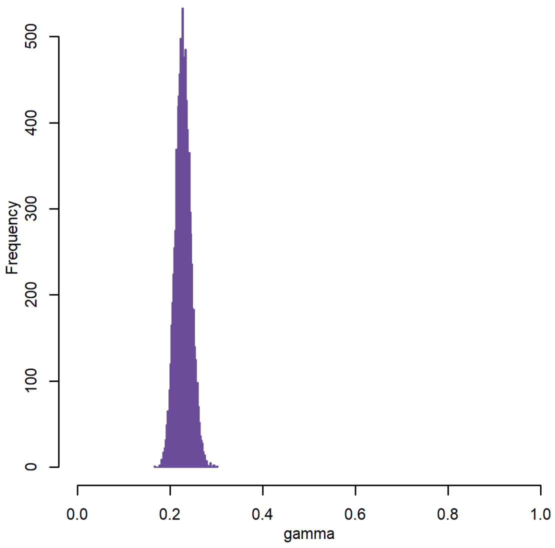

### Cohort 53

Author: Lindhart

Publication year: 1939

Patient recruitment years: 1931-1932

Cohort size: 1342

Location: Denmark

Type of TB (smear-status): positive

Details: Mortality of notified TB
cases in Denmark between 1925 and 1934.
Female patients only.

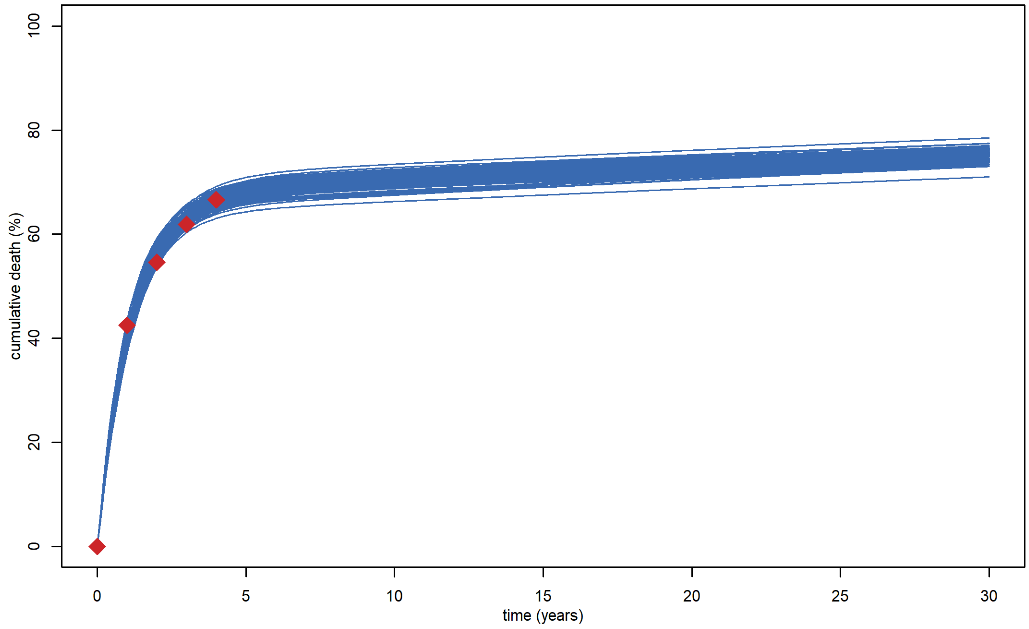


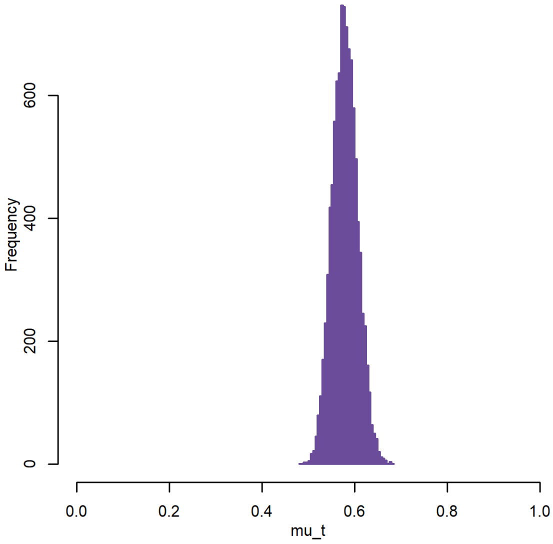


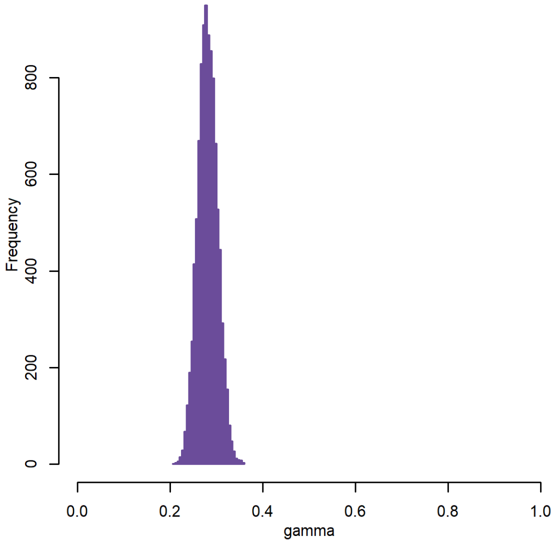

### Cohort 54

Author: Lindhart

Publication year: 1939

Patient recruitment years: 1932-1933

Cohort size: 1398

Location: Denmark

Type of TB (smear-status): positive

Details: Mortality of notified TB cases in Denmark between 1925 and 1934. Female patients only.

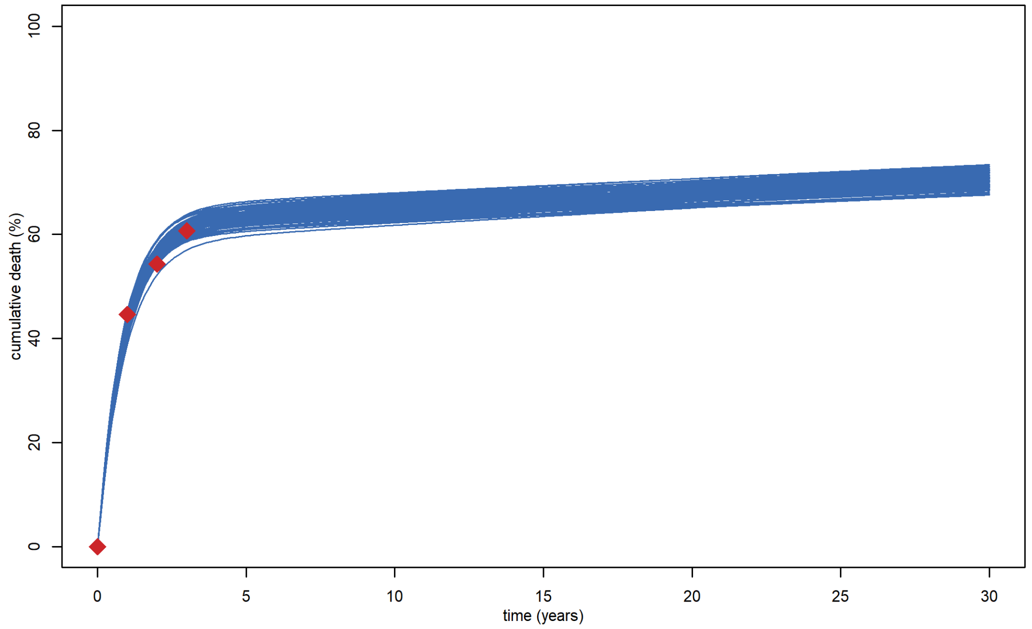


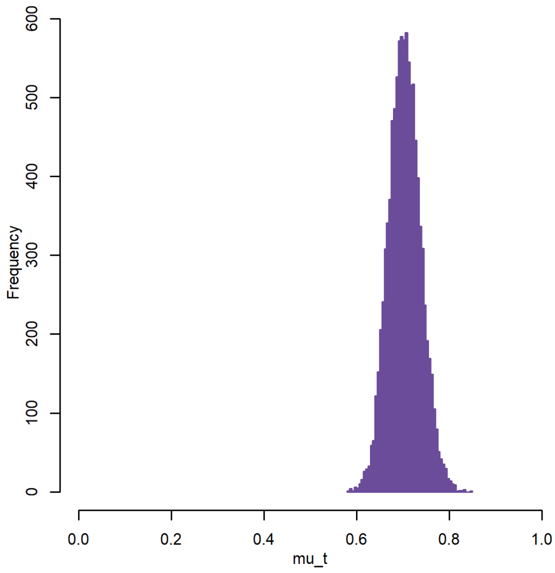


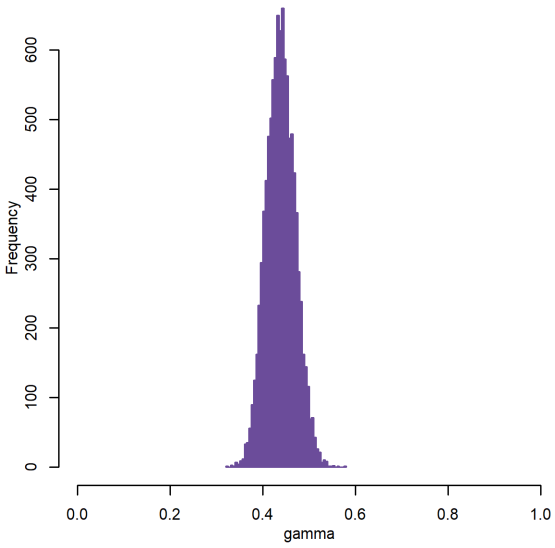

### Cohort 55

Author: Lindhart

Publication year: 1939

Patient recruitment years: 1933-1934

Cohort size: 1205

Location: Denmark

Type of TB (smear-status): positive

Details: Mortality of notified TB
cases in Denmark between 1925 and 1934.
Female patients only.

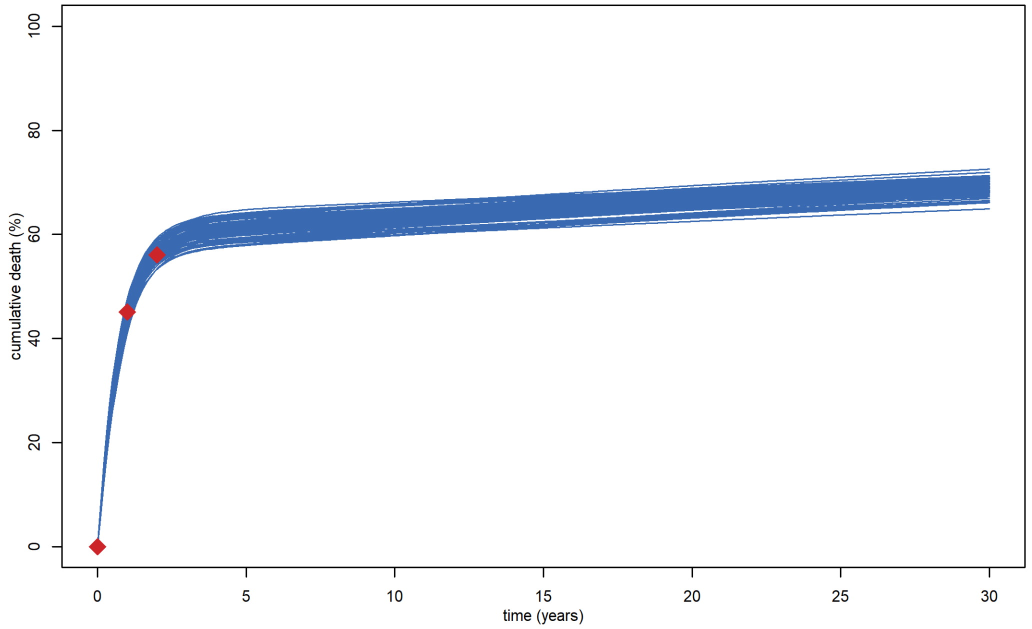


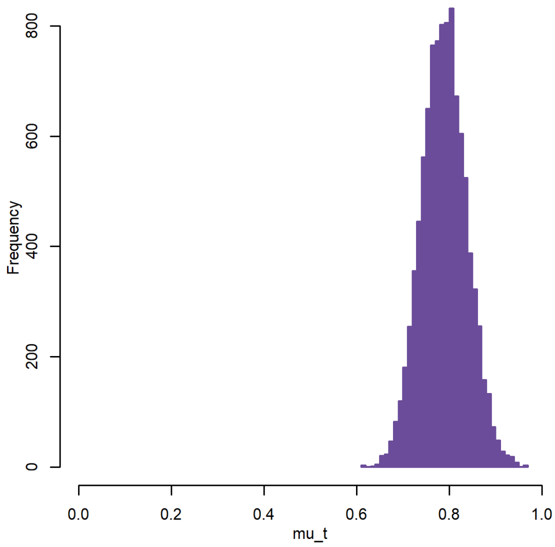


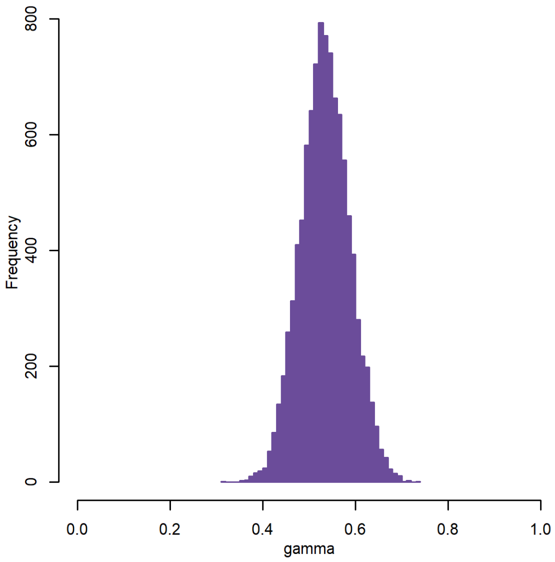

### Cohort 56

Author: Lindhart

Publication year: 1939

Patient recruitment years: 1934-1935

Cohort size: 1057

Location: Denmark

Type of TB (smear-status): positive

Details: Mortality of notified TB
cases in Denmark between 1925 and 1934.
Female patients only.

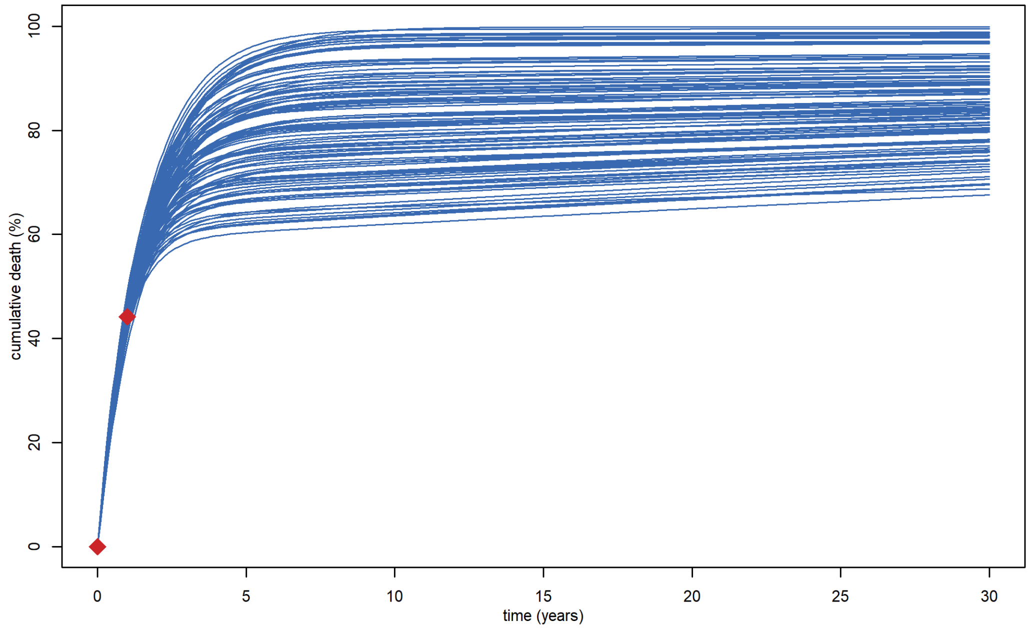


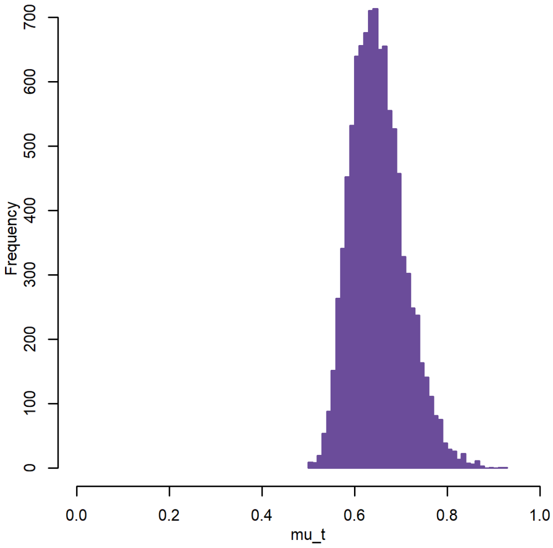


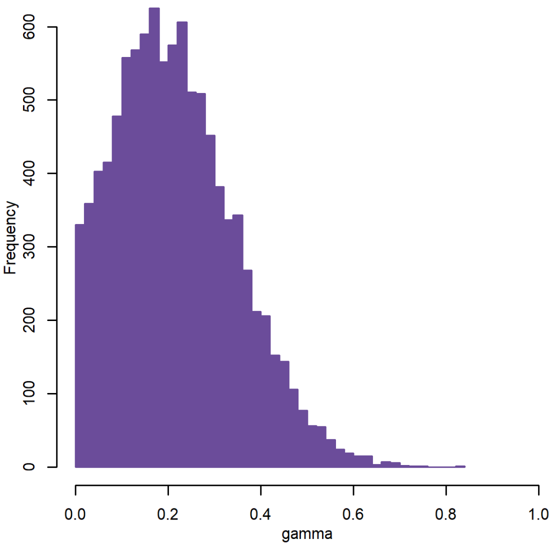

### Cohort 59

Author: Magnusson

Publication year: 1938

Patient recruitment years: 1916-1935

Cohort size: 166

Location: Iceland

Type of TB (smear-status): positive

Details: Cases admitted for treatment at the Vifillsstadir Sanatorium in Reykjavik. Male patients only.

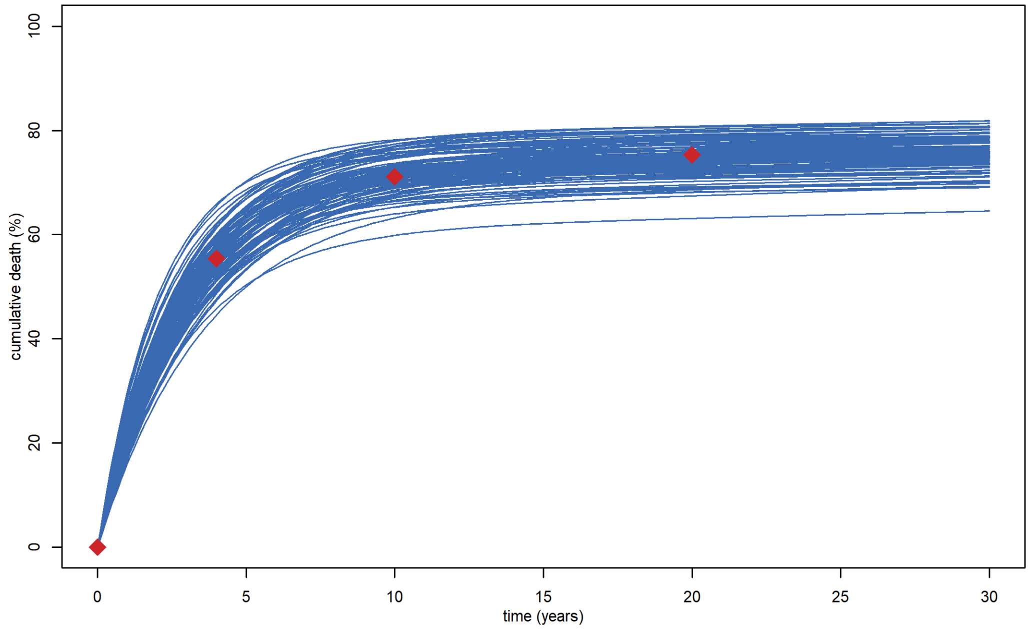


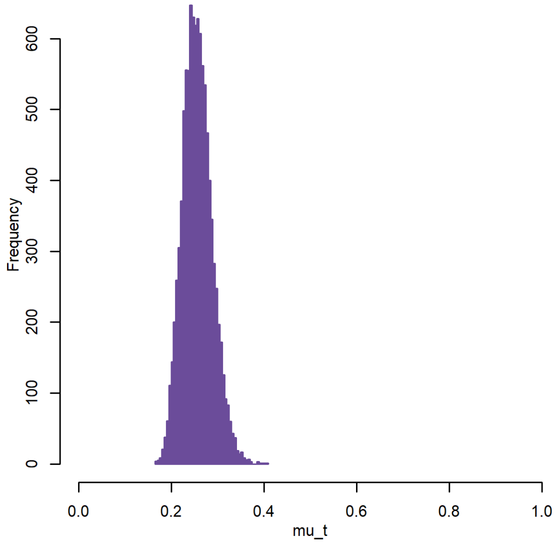


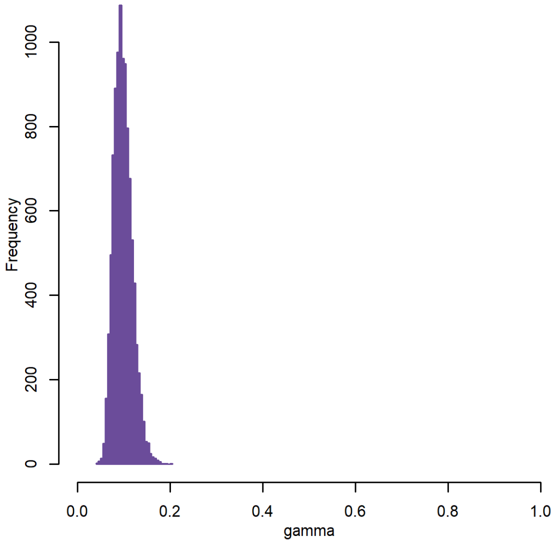

### Cohort 60

Author: Magnusson

Publication year: 1938

Patient recruitment years: 1916-1935

Cohort size: 213

Location: Iceland

Type of TB (smear-status): positive

Details: Cases admitted for treatment at the Vifillsstadir Sanatorium in Reykjavik. Female patients only.

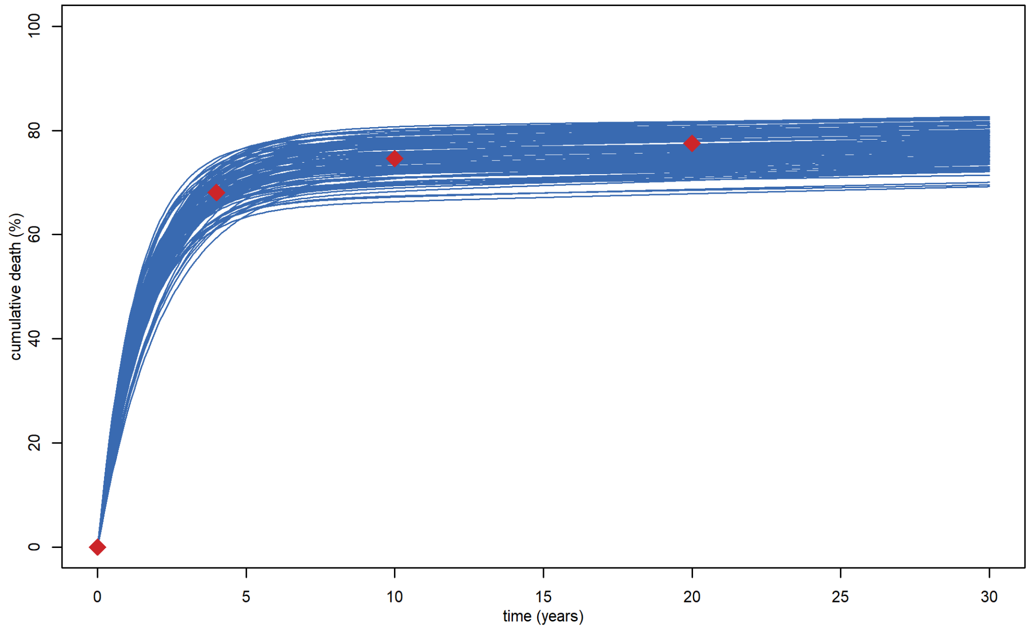


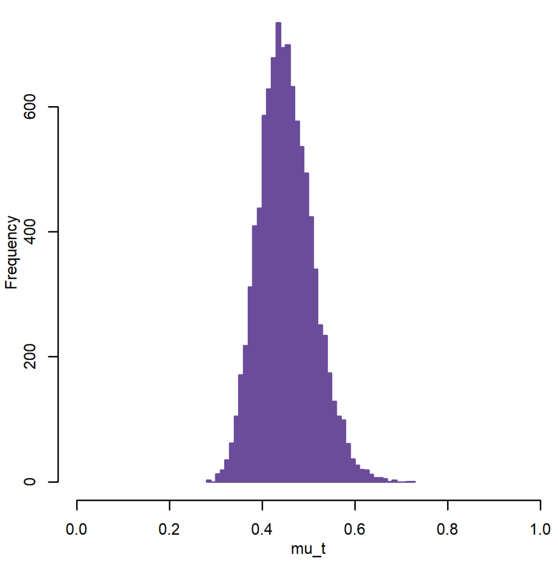


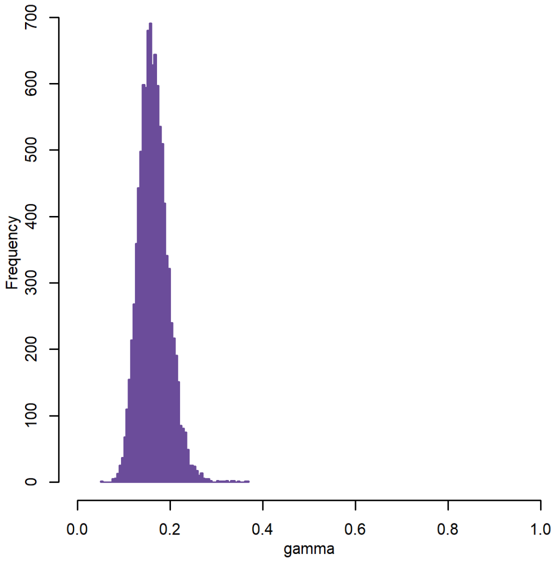
